# The limits to ecological limits to diversification

**DOI:** 10.1101/2022.05.13.491338

**Authors:** Rampal S. Etienne, Bart Haegeman, Álvaro Dugo-Cota, Carles Vilà, Alejandro Gonzalez-Voyer, Luis Valente

## Abstract

While the theory of micro-evolution by natural selection assigns a crucial role to competition, its role in macroevolution is less clear. Phylogenetic evidence for a decelerating accumulation of lineages suggests a feedback of lineage diversity on diversification, i.e., ecological limits to diversification. However, does this feedback only occur between close relatives, or do distant relatives also influence their diversification? In other words: are there phylogenetic limits to these ecological limits? Islands form ideal systems to answer these questions, because their boundedness facilitates an overview of all potential competitors. The DAISIE (Dynamic Assembly of Island biota through Speciation Immigration and Extinction) framework allows for testing the presence of diversity-dependence on islands given phylogenetic data on colonization and branching times. The current inference models in DAISIE assume that this diversity-dependence only applies within a colonizing clade, which we term clade-specific (CS) diversity-dependence. Here we introduce a new DAISIE model that assumes that diversity-dependence applies to all species regardless of their ancestry, i.e. diversity-dependence applies both to species within the same clade and between different clades. We call this island-wide (IW) diversity-dependence. Here we present a method to compute a likelihood for this model and develop a statistical procedure based on likelihood ratio bootstrapping to compare it to the likelihood of the CS model in order to overcome biases known for standard model selection. We apply it to the diversification of *Eleutherodactylus* frogs on Hispaniola. Across the Greater Antilles archipelago, this radiation shows repeated patterns of diversification in ecotypes which are similar across clades. This could be suggestive of overlapping niche space and hence between-clade interactions, i.e. IW diversity-dependence. But it could also be suggestive of only within-clade interactions, because between-clade interactions would have blocked the same ecotype re-appearing. We find that the CS model fits the data much better than the IW model, indicating that different colonizations, while resulting in similar ecotypes, are sufficiently distinct to avoid interacting strongly. We argue that non-overlapping distributions between clades (both spatially and in terms of ecotypes) cannot be used as evidence of CS diversity-dependence, because this pattern may be a consequence of IW diversity-dependence. By contrast, by using phylogenetic data rather than distributional data our method does allow for inferring the phylogenetic limits to ecological limits to diversification. We discuss how our new IW model advances our understanding also in other ways, ranging from identifying priority effects to modelling the spread of an epidemic in island-like systems, such as schools or hospitals.

## Introduction

“As species of the same genus have usually, though by no means invariably, some similarity in habits and constitution, and always in structure, the struggle will generally be more severe between species of the same genus, when they come into competition with each other, than between species of distinct genera”. This statement by Darwin in the Origin of Species (Darwin, 1859), known as the competition-relatedness hypothesis (Cahill et al., 2008) or the phylogenetic limiting similarity hypothesis (Violle et al., 2011), or Darwin’s naturalization hypothesis in the field of invasion biology (Proches et al., 2008), has been the subject of debate over the last decades, particularly in the field of phylogenetic community ecology (Mayfield and Levine, 2010; HilleRisLambers et al., 2012; Pigot and Etienne, 2015; Gerhold et al., 2015; Narwani et al., 2015; Germain et al., 2016; Cadotte et al., 2017; Wilcox et al., 2018). The consequences of the competition-relatedness hypothesis for macroevolution have received much less attention. Darwin (1859) formulated these consequences himself as “each new variety or species, during the progress of its formation, will generally press hardest on its nearest kindred, and tend to exterminate them.” This implies that with increasing diversity, speciation rates decline or extinction rates increase. This phenomenon has been referred to as diversity-dependent diversification (also somewhat confusingly called density-dependent diversification) since the 1970s (Raup et al., 1973; Walker and Valentine, 1984). Rabosky (2013) distinguishes Darwinian diversity-dependence, which does not imply an upper bound, from asymptotic diversity-dependence, which by definition does impose an upper bound on diversity. We leave the question aside whether an upper bound exists, and rather focus on the commonality of these types of diversity-dependence: that ecological interactions affect diversification. There have been many suggestions of how such ecological limits operate (Rabosky, 2013). Here we are interested whether there is a phylogenetic limit to the effect of these ecological limits, i.e. whether diversity-dependence only acts between closely related species or (also) between distantly related species. There is considerable support for diversity-dependence in clades of phylogenetically closely related species (Foote and Miller, 2006; Phillimore and Price, 2008; Rabosky and Glor, 2010; Etienne et al., 2012; Jønsson et al., 2012; Foote et al., 2018), but there is also some evidence that phylogenetically distantly related (but ecologically similar) taxa reduce each other’s diversification rates (Stanley, 1973; Sepkoski, 1996; Valkenburgh, 1999; Jablonski, 2008; Silvestro et al., 2015; Pires et al., 2017). However, the latter evidence is relatively scarce and comes mostly from fossil data. The question then presents itself whether molecular phylogenies can also inform us about the phylogenetic limits to these ecological limits to diversification.

We propose that islands are the ideal arena to study these questions, because they are clearly defined systems where (exceptional) radiations have occurred. Moreover, as islands tend to be depauperate, we see cases where species released from competition have radiated to fill niches usually occupied by a different clade, e.g., woodpecker finches in the Galápagos. In MacArthur & Wilson’s original work on island biogeography (MacArhur and Wilson, 1967) speciation receives little attention and therefore also diversity-dependent speciation, but colonization and extinction are diversity-dependent in their theory, as per capita colonization rates decrease and per capita extinction rates are assumed to increase with increasing island diversity. The General Dynamic Model of island biogeography (Whittaker et al., 2008) explicitly assumes that the island’s carrying capacity influences the diversification rates. However, neither of these classic works discusses the phylogenetic nature of the ecological limits to diversification. Here, we consider two types of diversity-dependence: the clade-specific (CS) level, where only species that descend from the same mainland species (possibly through multiple colonizations) reduce each other’s speciation rate and colonization rate, and the island-wide (IW) level, where all island species, that may descend from very different mainland ancestors, inhibit each other’s speciation and colonization. The CS scenario can be modelled by assuming a carrying capacity or upper limit to the number of species for each clade, while the IW model can modelled by assuming an island-wide carrying capacity or upper limit to the total number of species. Diversity-dependence in speciation rates and colonization rates has been incorporated in the DAISIE framework (Dynamic Assembly of Island biota through Speciation, Extinction and Immigration, Valente et al. (2015)) that allows estimating rates of colonization, speciation and extinction from phylogenetic data of the clades that colonized an island (or archipelago). In the first simulations in this framework, diversity-dependence was of the IW-type (Valente et al., 2014). For inference (i.e. parameter estimation), only the CS model was implemented (Valente et al., 2015), using insight from analyses on single clades of closely-related species (Rabosky and Lovette, 2008; Etienne et al., 2012), because the IW model presented technical difficulties. Here we overcome (some of) these technical difficulties by presenting a method to compute the likelihood of colonization and branching events under the IW model.

We illustrate our method with an application to the colonization of Hispaniola by five lineages of *Eleutherodactylus* frogs (genus *Eleutherodactylus*; Dugo-Cota et al., 2019), for which both CS and IW models can be verbally argued to apply. On the one hand, these lineages show, across the Greater Antillean archipelago, repeated patterns of diversification into a similar set of ecotypes (Dugo-Cota et al., 2019), suggesting a limited set of niches is available, which in turn implies that diversity-dependence acts, but no further than within each clade (CS). On the other hand, the relatively low geographic overlap in ecotypes between clades on Hispaniola suggests that diversity-dependence extends to all species on the island (IW) because species may have blocked colonization of the same ecotype regardless of their phylogenetic relatedness. Our analysis, using only phylogenetic data, shows that the CS model fits the data much better than the IW model. We discuss this result and provide suggestions for further research avenues.

## Methods

Under the original DAISIE inference model (Valente et al., 2015) and its subsequent extensions (Valente et al., 2017a,b, 2019) species can colonize an island at a rate *γ*, go extinct at a rate *μ*, and speciate via cladogenesis (when one island species splits into two, forming two new endemic species) at a rate *λ*^*c*^ or via anagenesis (when one island species diverges from its mainland ancestor becoming a new endemic species, without leading to an increase in diversity on the island) at a rate *λ*^*a*^. CS-type diversity-dependence is implemented by allowing for rates of cladogenesis and colonization to decline with increasing diversity within a clade, with the number of species within each clade being limited by a CS carrying capacity, *K*. The maximum-likelihood implementation of DAISIE allows *γ, μ, λ*^*c*^, *λ*^*a*^ and *K* to be estimated based on the distribution of times of island colonization and branching times within an island, extracted from divergence-dated molecular phylogenies. A diversity-independent model (DI) is also implemented, i.e. by fixing *K* to infinity so that *λ*^*c*^ and *γ* do not decline with diversity.

A logical alternative model to CS in the island context is the IW model, where instead of a *K* per clade there is an island-wide *K* that determines the maximum number of species that can coexist on an island across all clades. This model was implemented in the first version of DAISIE, but only in simulations (Valente et al., 2014). Until now, estimating parameters of an IW model has not been attempted, because a) the model equations are rather cumbersome to write down and implement, and b) parameter estimation is computationally demanding in terms of memory requirements and runtimes, even for small data sets. Here we take on these hurdles. We develop a method of estimating parameters of an IW model from phylogenetic data. The data requirements, parameters and simulation approach of the DAISIE IW model are the same as for CS, except that diversity-dependence in *λ*^*c*^ and *γ* is determined by an island-wide *K*, so that these rates decline with diversity of all island species rather than simply diversity of the colonist clade they belong to.

### Likelihood of colonization and branching data for the IW model

We compute the likelihood of the data, consisting of colonization and branching events, for the IW model using the *Q*-approach (Etienne et al., 2012; Laudanno et al., 2019). This approach is named after the quantity *Q*(*t*), which is the probability that a random realisation of the model is consistent with the data up to an arbitrary time *t*. In the supplementary material we construct the differential equations governing the dynamics of *Q*(*t*), and explain how these equations, which apply to the dynamics between colonization and branching events, are connected to one another across the colonization and branching events. By solving these equations from the island emergence time to the present, we obtain *Q*(*t*_*p*_), the quantity *Q*(*t*) evaluated at the present time *t*_*p*_, from which the likelihood can be extracted (see supplementary material for details).

Our computational procedure is based on the assumption that we have full information about the extant species. That is, we assume that the island phylogenies of the full set of extant species are known, together with the corresponding colonization times. This assumption simplifies not only the likelihood computation, but also the comparability with the CS likelihood. Indeed, in case of partial sampling from the phylogeny, the CS model distinguishes to what clade the missing species belong, while the IW model does not, making their likelihood incomparable. To guarantee full comparability we have also treated the likelihood of the CS model as a product of IW likelihoods with mainland pool size of 1 across the *M* mainland species. That is, CS and IW only differ in whether clades established by mainland ancestors are independent (CS) or are connected through each other’s diversity (IW).

### Model fitting

We fitted five DAISIE models: a model without diversity dependence (DI, 4 free parameters), a model with clade-specific diversity dependence (CS-DD, 5 free parameters), a model identical to CS-DD but without anagenesis (CS-DD-noA, 4 free parameters); a model with island-wide diversity dependence (IW-DD, 5 free parameters), and a model identical to IW-DD but without anagenesis (IW-DD-noA, 4 free parameters). In all DD models the per capita rates of cladogenesis 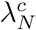 and colonization *γ*_*N*_ were assumed to linearly decline with diversity:

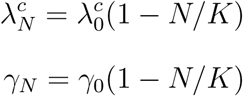

where *N* is the total number of species in a clade in the CS model, and the total number of species on the entire island in the IW model. *K* is the carrying capacity per clade for the CS model (hence the same for each clade) and for the entire island for the IW model. We follow the original DAISIE model (Valente et al., 2015) by assuming no diversity-dependence in extinction or anagenesis. Diversity-dependence in extinction (higher extinction rates for higher diversity) is conceivable, but seems unlikely because it causes a stronger pull-of-the-present in lineages-through-time plots contrary to what is commonly observed (Phillimore and Price, 2008; Etienne et al., 2012). For anagenesis, diversity-dependence seems biologically unlikely because speciation by anagenesis can occur through drift alone simply due to long-term isolation from the mainland. Even if local adaptation is the primary cause of anagenesis, competition with other species (and thus diversity-dependence in anagenesis) is unlikely to prevent anagenesis. In such a scenario the species will probably not establish at all (some adaptation seems necessary to survive in a new environment) which would be accounted for by diversity-dependence in the colonization rate.

### Phylogenetic data

We used the dated phylogeny of *Eleutherodactylus* frogs by Dugo-Cota et al. (2019), which is based on four mitochondrial and three nuclear genes. The data set comprises 152 species of the genus, including 148 Caribbean species, i.e. 89% of the Caribbean diversity, as well as four continental species. The divergence dated phylogeny was reconstructed in BEAST v1.8.2, using secondary time calibration points extracted from the wider eleutherodactyline phylogeny of Heinicke et al. (2007). Dugo-Cota et al. (2019) reconstructed the biogeographical history of Caribbean *Eleutherodactylus* using BioGEOBEARS (Matzke, 2013, 2014) with a time-stratified analysis and nine geographical regions. They inferred five colonizations of Hispaniola from the mainland and surrounding islands (which are collectively referred to as the mainland hereafter), each of which radiated on the island, to a great or lesser degree, producing five in situ radiations of 28, 21, 8, 5 and 3 species (Table 1).

**Table 1.**
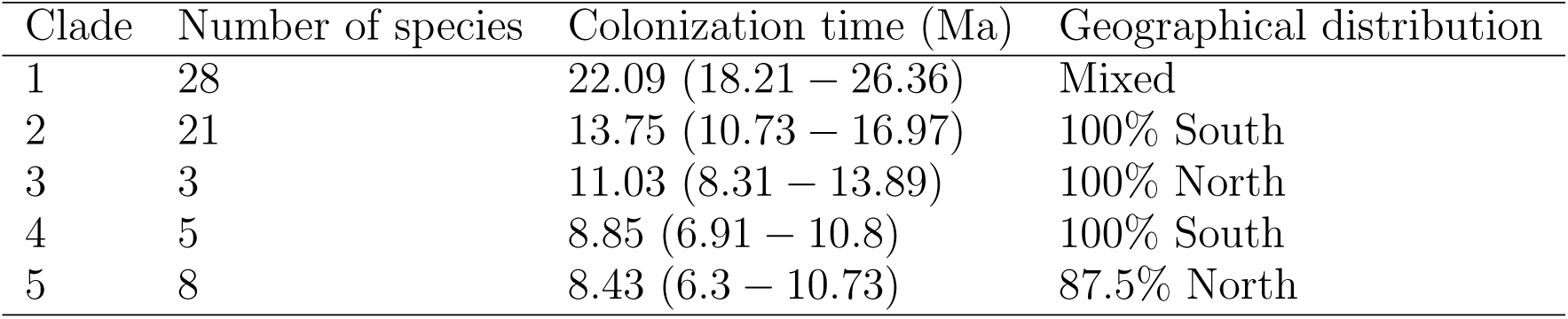
Characteristics of the five clades on Hispaniola: diversity, colonization times and geographical distribution within the island of Hispaniola.

The Dugo-Cota et al. (2019) phylogeny includes 57 of the 66 Hispaniola species. Because fitting the IW model assumes complete sampling of the extant species of the focal island, we inserted the missing Hispaniola species by assigning them at random locations within the Hispaniola subclades that they have been hypothesized to belong to. Information on the nine missing species and the detailed rationale for including them in a given subclade are given in supplementary Table S1. There is no genetic data available on GenBank for these missing species, because they have been recently described, are known from a single specimen or are possibly extinct. We used a set of functions from the phytools R package to assign missing species to clades (Revell, 2012). Four of the missing species were previously considered subspecies, and have recently been elevated to species, and we thus randomly inserted them 100 times at any height along the tip branch of the species they were previously assigned to. Four species have been proposed to belong to well-defined terminal clades based on morphology, and we randomly inserted them at any position and at any height within those clades (100 times). We repeated this procedure 100 times on the five clades from the maximum clade credibility tree from BEAST, producing 100 sets of five clades with complete sampling. The exact procedure is detailed in Table S1. One of the missing species, *E. neiba* was not added to the tree because there is no previous hypothesis regarding its phylogenetic position. We ran a sensitivity analysis including this species as a separate colonization, to assess whether in the unlikely case it formed a separate clade this would affect the results. These analyses showed that even if *E. neiba* formed an independent colonization, the same model would still be preferred. We therefore did not include it in the analyses, as it is unlikely to modify the main findings.

We extracted colonization and branching times for each of the five Hispaniola radiations from these data. Colonization times were assumed to be the stem ages of the Hispaniola clades. Information on each of the Hispaniola clades is given in Table 1 and the phylogeny is shown in Fig. 1.

**Figure 1.**
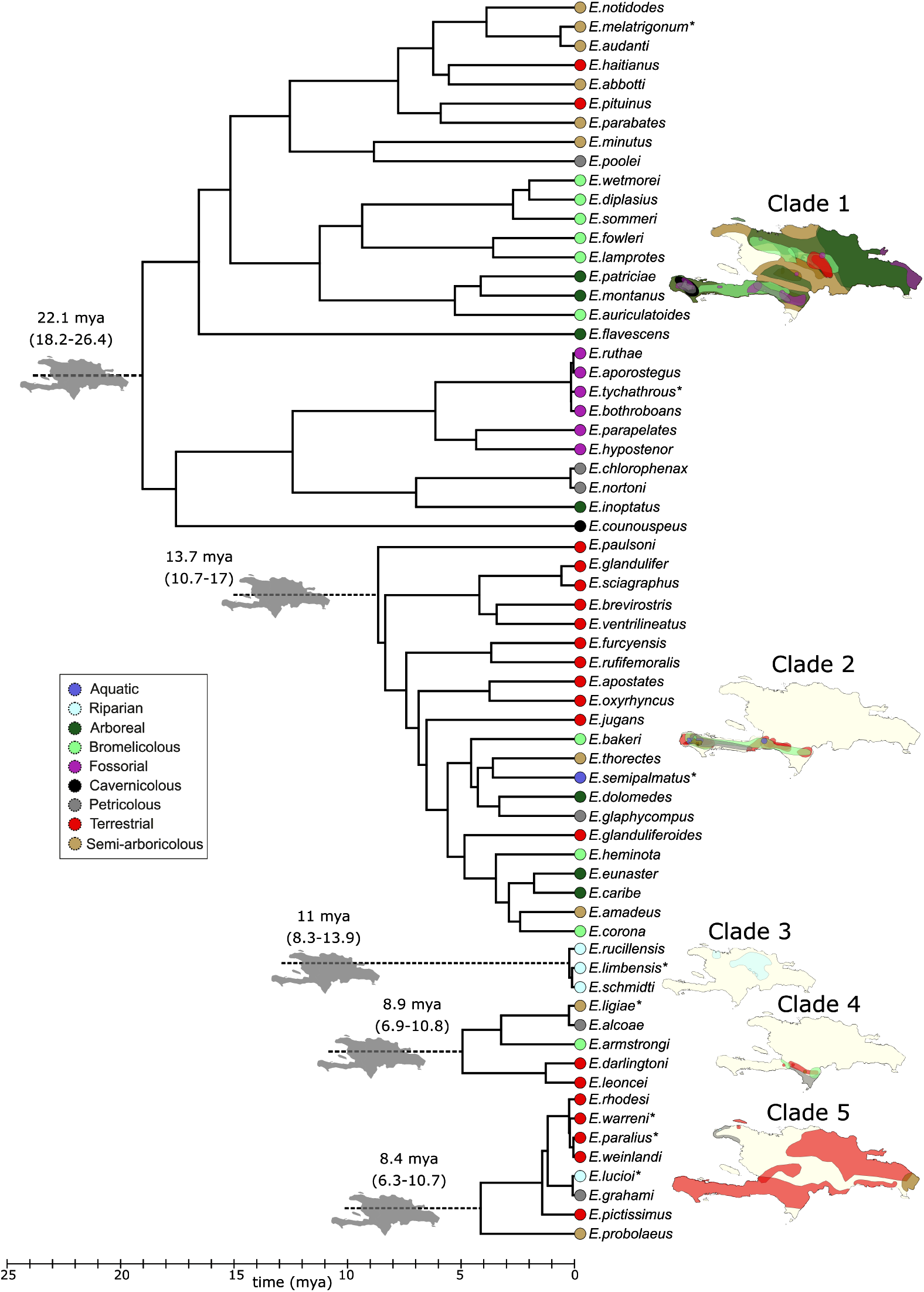
Phylogeny of the Hispaniola *Eleutherodactylus* frogs. A separate time-calibrated phylogenetic tree is shown for each of the five clades. Gray islands show the five inferred independent colonizations of Hispaniola. Colors at the tips of the phylogeny represent the species ecotypes (see legend). The same colors are used to show the species distributions for each independent colonization on the Hispaniola map (for visual clarity some transparency has been applied). The asterisk indicates where missing species have been added to phylogeny according to taxonomic information, see Table S1.

As the downstream DAISIE analyses are computationally demanding, we wanted to use only one data set for subsequent analyses. In order to perform an informed selection of the tree, we fitted the CS and the IW model with no anagenesis to each of the 100 sets of trees. The results of the analysis on the 100 sets of trees are shown in Table S2. The preferred model in all trees was CS. We thus used only tree set 52, which is the one with the highest likelihood for CS, for all subsequent analyses (hereafter ‘empirical data set’). This may seem to introduce a bias in favor of the CS model, but we note that all parameter estimates and all loglikelihood differences were very similar across the 100 sets of trees, and would have led to the same conclusions in our model comparison. Moreover, tree set 52 did not have the largest loglikelihood difference between CS and IW. All sets of trees and corresponding DAISIE colonization/branching time R objects are provided in the supplementary material. The simulation code, functions to compute the likelihood under the two models, and a tutorial on how to run simulations and perform model-fitting are available in the R package DAISIE on CRAN and on Github (https://github.com/rsetienne/DAISIE).

### Likelihood optimization on the empirical data

We fitted each of the five DAISIE models 5 times to the empirical data set using different random sets of starting parameters to avoid being trapped in local likelihood optima. We assumed an island age of 30 million years, consistent with the paleogeographical reconstruction of Iturralde-Vinent (2006) for when Hispaniola was isolated from other landmasses. The mainland pool size *M* was set to 1000 frog species. We note that this value is not crucial, because mainland pool size affects only the rate of colonization; the product of mainland pool size and the rate of colonization, i.e. the total rate of colonization, is practically constant (Valente et al., 2019). Indeed, parameter estimates were very similar for optimizations with *M* = 300.

Maximum likelihood optimizations were run on the high-performance (Peregrine) cluster of the University of Groningen. Optimization of DI and CS-type models generally converged in a few hours. IW-DD model optimizations took between a few hours up to 10 days to complete.

### Goodness-of-fit

We simulated 5,000 data sets using the maximum likelihood parameters of the preferred CS-type model (CS-DD no anagenesis) and preferred IW-type model (IW-DD no anagenesis), hereafter the CS and IW models. We then plotted relevant statistics from the simulated data sets and compared them to those in the empirical data to study how well the models fit the data.

### Bootstrap analysis

We computed the AIC and BIC values and weights for model comparison, but because model selection involving diversity-dependent models is known to be troublesome (Etienne et al., 2016), we developed a parametric bootstrap analysis of both CS and IW models in order to perform model selection using a likelihood ratio bootstrap test. In addition, this bootstrap analysis allowed us to assess bias and precision of parameter estimates. We chose the first 1000 out of the 5000 data sets from each of the CS and IW simulations. Not all 5000 simulated data sets were used, because the subsequent analyses on these data sets were computationally demanding. For each of the chosen 2000 data sets we fitted both CS and IW models, resulting in a total of 4000 maximum likelihood optimizations. We used as starting values the ML parameters obtained in the optimization analyses for the empirical data for the given model, unless this was not possible. For instance, if a clade in data sets generated under the IW model had more species than the value of *K* estimated for the CS model fitted to the empirical data, using that *K* as a starting value to fit the CS model to the IW-simulated data would give a likelihood of 0. Therefore we calculated the starting *K* for each data set using the largest value of either the *K* estimated from the empirical data for the given model being fitted, or the maximum number of species in a clade (CS model) or the total number of species on the island (IW model) in the simulated data set.

For the bootstrap likelihood ratio test we compared the logarithm of the likelihood ratio of CS and IW in the empirical data (i.e. loglikelihood difference, loglikelihood of the CS model *−* loglikelihood of the IW model) with the distribution of the logarithm of likelihood ratios from the data sets simulated under CS and under IW (1000 data sets each). We computed the 95th percentile of the distribution under the IW model. If the loglikelihood difference of the empirical data falls to the right of this value, then they are unlikely to be produced by the IW model, and if it is well within the distribution of the data generated under the CS model, the CS model is selected. We also computed the 5th percentile of the distribution under the CS model. If the loglikelihood difference of the empirical data falls to the left of this value, then they are unlikely to be produced by the CS model, and if it is well within the distribution for the data generated under the IW model, the IW model is selected. If the loglikelihood difference of the empirical data falls between the two percentiles, then no model can be selected decisively.

## Results

### Likelihood optimization on the empirical data

Convergence of the five independent optimizations per model to the empirical data set was very good, with all five runs finding the same maximum likelihood parameter set for each model. The preferred model using both AIC or BIC was CS-DD with no anagenesis (four free parameters) (Table 2). The loglikelihood difference between the best CS model and the best IW model (both diversity-dependent) was 6.43. This value points to the CS model as the best model also in the likelihood ratio bootstrap test (see below). The models without anagenesis had virtually the same parameter values as their counterparts allowing anagenesis to be different from 0. That is, the latter models had estimated rate of anagenesis that were very close to 0. This is to be expected because all five frog clades radiated and anagenesis (before cladogenesis) will then go largely unnoticed. The only signal of anagenesis in such a case could come from the observation of recolonizations of the same mainland species that established the clade(s). This is because the model assumes that recolonizations can only occur after speciation has taken place (if it happens before speciation takes place, the recolonization is assumed to reset the colonization time and is then not observed). As the data did not contain recolonizations, the maximum likelihood estimate of anagenesis is expected to be 0. Across the 100 sets of empirical trees the loglikelihood difference between the diversity-dependent CS and IW models ranged from 5.16 to 6.63, all of which suggest the CS model is highly preferred.

**Table 2.** Maximum likelihood parameter estimates and information criteria of the DAISIE models fitted to the empirical dataset. df is the number of free parameters. The asterisks indicate that the values are very close to 0, but not fixed at 0.

| model | $\lambda_0^c$ | $\mu$ | $K$ | $\gamma$ | $\lambda^a$ | LL | df | AIC | AIC weight | BIC | BIC weight |
| --- | --- | --- | --- | --- | --- | --- | --- | --- | --- | --- | --- |
| DI | 0.18 | 0.03 | ∞ | 0.0002 | 0* | -215.87 | 4 | 439.75 | 0.00 | 445.53 | 0.00 |
| DD-CS | 0.44 | 0.11 | 36.44 | 0.0002 | 0* | -208.67 | 5 | 427.34 | 0.27 | 434.58 | 0.15 |
| DD-CS-noA | 0.44 | 0.11 | 36.45 | 0.0002 | 0 | -208.67 | 4 | 425.34 | 0.73 | 431.13 | 0.85 |
| IW-CS | 0.40 | 0.17 | 131.89 | 0.0003 | 0* | -215.10 | 5 | 440.20 | 0.00 | 447.43 | 0.00 |
| IW-CS-noA | 0.40 | 0.17 | 131.96 | 0.0003 | 0 | -215.10 | 4 | 438.20 | 0.00 | 443.98 | 0.00 |

### Goodness-of-fit

Using the estimated parameters for the diversity-dependent CS and IW models we generated simulated data for which we computed several summary statistics. The distributions of the summary statistics across these simulations fitted well with the empirical data for both models, but somewhat better for the CS model. (Fig. 2, Fig. S1, Fig. S2), as the empirical statistics and the medians across the simulations are slightly more similar for this model.

**Figure 2.**
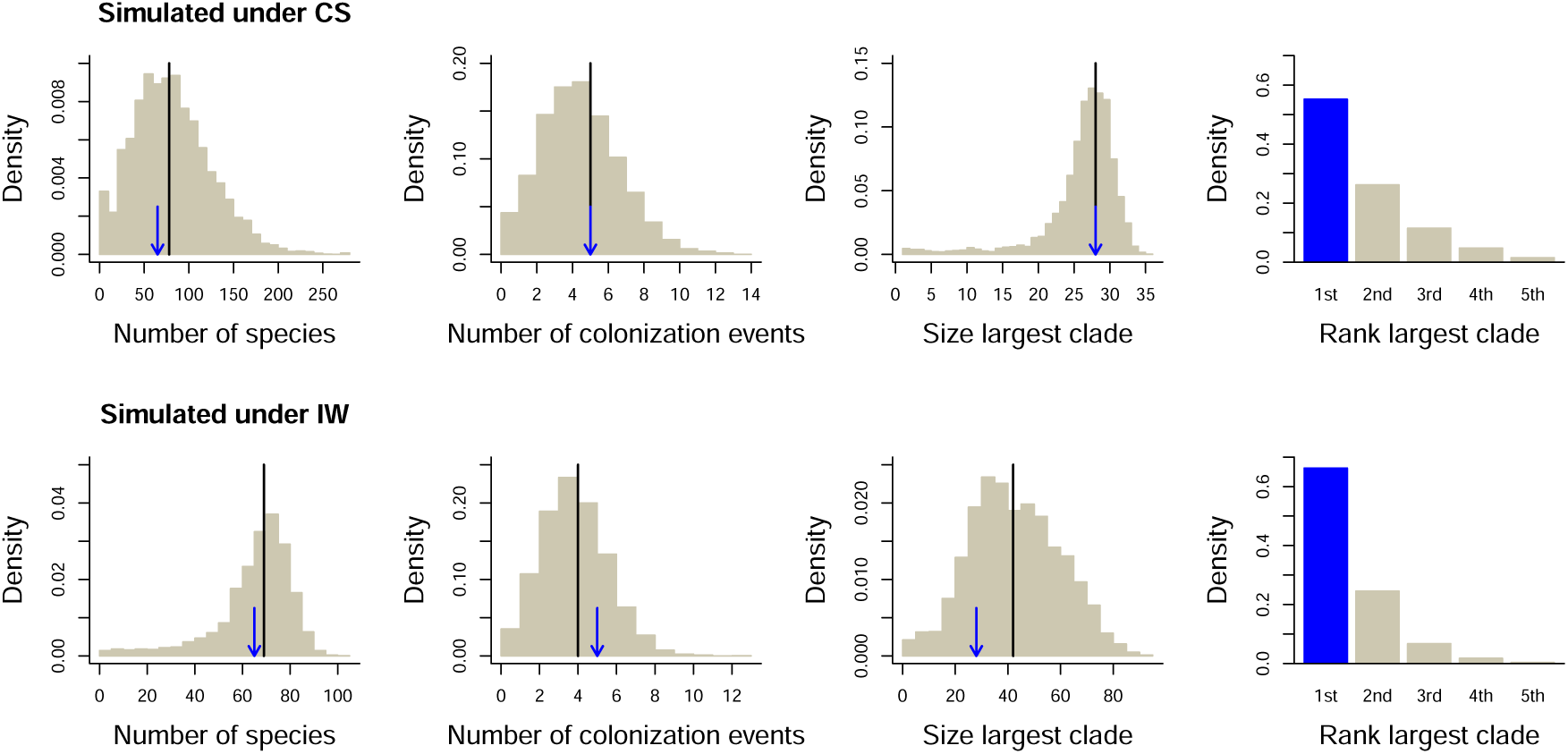
Goodness-of-fit plots. Distributions of relevant metrics (number of species, number of colonizations leading to extant clades, size of the largest clade, and rank of the largest clade when clades are ordered according to their colonization time, rank 1 corresponding to the first colonization) obtained from 5,000 data sets simulated with the maximum likelihood parameters of the CS (top row) and IW (bottom row) models. Black line: median value, blue arrow and blue bar: value in the empirical data.

### Bootstraps

The analyses fitting the CS model to each of 1000 CS and 1000 IW simulated data sets all completed successfully. For the analyses fitting the IW model to the same data sets, some runs runs could not be completed within the limit we set (10 days). For the CS data sets this was 0.8% of the simulations and for IW this was 1.6%. These were all data sets with little information (only a single clade) where the estimation procedure went to very high values of the rate of cladogenesis and colonization. The maximum IW loglikelihood is thus higher in these cases, although it may not be much higher. Even if we make the unlikely assumption that in all of these aberrant simulated data sets the IW is a better fit, they are so rare that our qualitative conclusion that the CS model is a better fit does not change.

Parameters were estimated with high precision and little bias under both models: the median and means of the distribution of parameters estimated under the CS and IW models for data sets simulated under those models closely matched the simulated values (Figs. 3 and 4). When fitting the CS model to IW simulations (Fig. S3) we observe that the *K* is estimated to be much higher than in the CS simulations (Fig. 3). This is because the IW simulations show more variability in clade sizes that can only be accommodated by the CS model by assuming a larger clade-level *K*. When fitting the IW model to CS simulations (Fig. S4), we do not observe such a discrepancy. Indeed, in this case the total number of species matters rather than the number of species per clade. All parameter estimates and corresponding loglikelihoods are available in Table S3.

**Figure 3.**
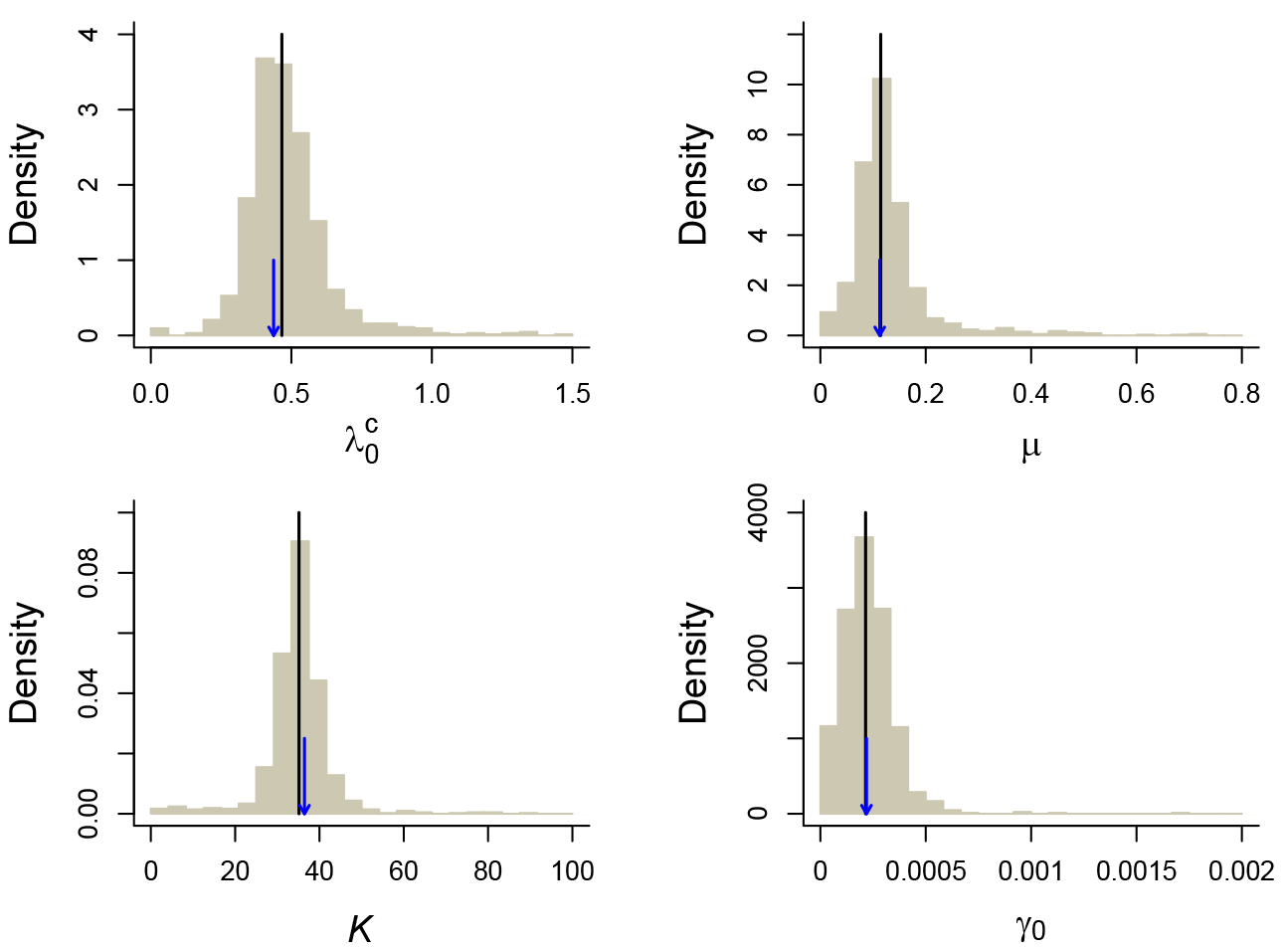
Bootstrap precision estimates of the parameters of the CS model. In a parametric bootstrap analysis the CS model was fitted to 1000 data sets simulated with the maximum likelihood parameters of the CS model for the empirical data. The panels show density histograms of the estimated parameters. The black lines indicate the median estimated values across all simulations and the blue arrows point to the values used in the simulations.

**Figure 4.**
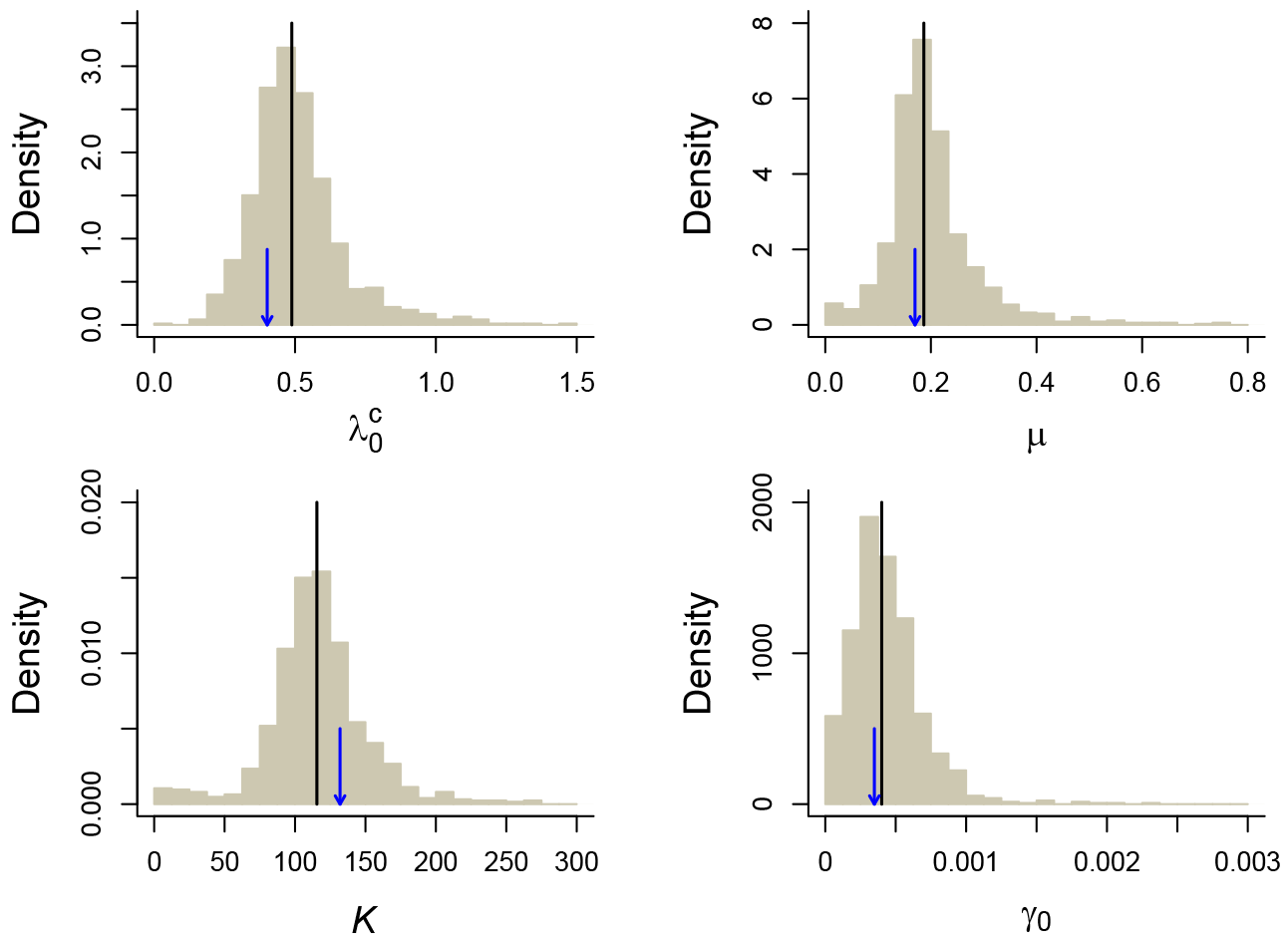
Bootstrap precision estimates of the parameters of the IW model. In a parametric bootstrap analysis the IW model was fitted to 1000 data sets simulated with the maximum likelihood parameters of the IW model for the empirical data. The panels show density histograms of the estimated parameters. The black lines indicate the median estimated values across all simulations and the blue arrows point to the values used in the simulations.

The simulated data can be used to check the reliability of model selection (because we know the generating process). When performing model selection by simply selecting the model with the highest likelihood, the IW model was incorrectly preferred over CS in 7.7% of data sets simulated under CS. The CS model was incorrectly preferred over IW in 19% of data sets simulated under IW. When imposing at least two log-units of difference before selecting a model, these numbers become 0.9% and 1.6%, respectively. The CS and IW model were then correctly selected in 77% and 40.6% of the corresponding data sets respectively, leaving 22.1% and 57.8% undecided between the two models. Because using highest likelihood or higher by at least two log-units is quite arbitrary, and still leads to either high type I error (highest likelihood) or low power (two log-units difference), we used the bootstrap likelihood ratio (or loglikelihood difference) distribution to set the permissible type I error to 5% (two left-most arrows in Fig. 5). This distribution of differences in loglikelihood between the CS and the IW model revealed that it was highly unlikely (*p <* 0.001) that the empirical loglikelihood difference (6.43) would have been found if the underlying model was IW, because the loglikelihood difference found in the empirical data (black arrow in Fig. 5) falls clearly beyond the tail of the distribution of loglikelihood differences obtained from data simulated under IW (higher than the largest likelihood), but falls right in the middle of the distribution of differences for data simulated under CS (at the 49.6th percentile). This suggests that the CS model is strongly supported as the best model for the empirical data.

**Figure 5.**
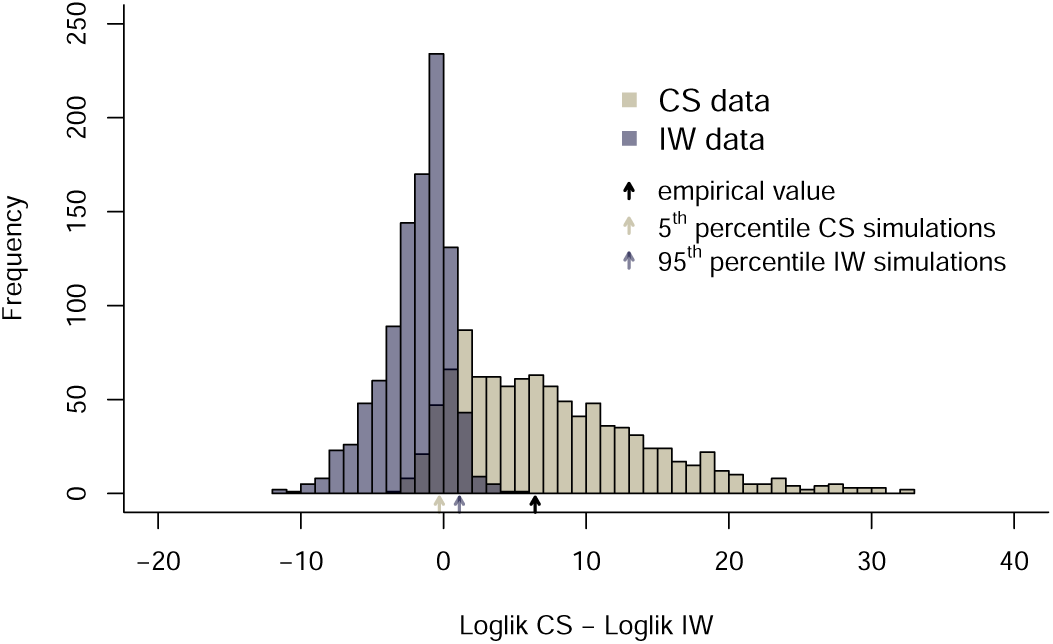
Likelihood ratio bootstrap test. Distribution of differences between the loglikelihood of the CS model and the loglikelihood of the IW model when fitting both models to data sets simulated under CS and IW. The rightmost (black) arrow shows the difference in the empirical data, while the leftmost and middle arrows indicate the 5th percentile and the 95th percentile of the distributions generated under the CS model and the IW model respectively. The black arrow falls well inside the distribution for data simulated under CS, and to the right of the 95th percentile (in fact even the maximum!) of the distribution for data generated under IW. Therefore, the CS model is strongly preferred.

The power to select the generating model is relatively high. The power to detect CS is 85% (part of distribution generated under CS model that is larger than the middle arrow in Fig. 5) whereas the power to detect IW is 72% (part of distribution under IW model that is smaller than the left-most arrow in Fig. 5). If the empirical data had had a loglikelihood ratio between -0.29 (the leftmost arrow in Fig. 5) and 1.11 (middle arrow in Fig. 5), model selection would have been indecisive.

## Discussion

We have developed a method to determine, using phylogenetic data on island colonization and branching times, whether diversity-dependence in rates of colonization and speciation is limited to species within a clade, or extends to species from different clades, or whether the information in the data is too limited to make a clear call. In Hispaniolan *Eleutherodactylus* frogs we find that models including diversity-dependence outperform models without a negative feedback of diversity on colonization and speciation rates, suggesting that ecological limits play an important role. Ecological limits operating at the clade-specific level (i.e. species from different colonizing clades do not interact) predominate over limits at the island-wide level (i.e. species from different clades reduce each other’s rate of colonization and speciation), because the model with clade-specific diversity-dependence clearly outperformed the model with island-wide diversity-dependence.

Although *Eleutherodactylus* frogs show repeated patterns of evolution into the same set of ecotypes (Dugo-Cota et al., 2019), these results suggest that these ecotypes do not interfere with each other across clades. One could argue that Fig. 1 already tells us this because the overlap in ecotypes and in ranges between the clades on Hispaniola is limited, and hence species do not seem to interact across clades. However, one can also explain this pattern as a *consequence* of interaction across clades, because under IW earlier clades block later ones from radiating into the same habitats (both ecologically and spatially). Our results do not support this explanation and lead us to conjecture that there has been sufficient (niche) space that IW diversity-dependence does not occur. Of course, this may change in time: if we wait millions of years, (niche) space may eventually become saturated, but currently there is no signal of IW diversity-dependence. In summary, present-day spatial distributions and ecological distributions into ecotypes cannot be taken as evidence that species from different clades do not interact, as these patterns may be a consequence of such interactions in the past. The approach we have taken in this paper is to infer such diversity-dependence from the phylogenetic branching pattern. We have shown that if IW diversity-dependence operates, we would often pick up its signal from the phylogenetic data. In our *Eleutherodactylus* frog example, we did not, as there is only 1% chance that the pattern we observed would be generated by an IW model (i.e. only 1 in 100 simulations of an IW model we would obtain a loglikelihood ratio between CS and IW models which is equal or higher than observed for the empirical data).

Our simulations were limited to the parameters estimated from the *Eleutherodactylus* frog data. To assess the more general ability of our approach to identify CS and IW when they are operating would require analyzing many more simulated data sets for a wide range of parameters sets. This is currently computationally unfeasible, because the likelihood maximizations, although performed with highly optimized code, take quite a bit of time (at least a few hours per data set), which bars extensive simulation studies across a sizeable number of replicates. Instead we suggest that researchers wishing to compare CS and IW models for their study system should fit these models to their data and take the estimated parameters to run simulations, just like we did here. This allows one to establish whether CS and IW models can be distinguished by plotting figures such as Fig. 5. We have shown that it is important to do so, because model selection based solely on AIC may be biased.

The CS model assumes the same carrying capacity *K* for each clade, which is a constraint to each clade’s size, and hence our model selection may be somewhat biased towards IW that only limits the overall number of species by its *K*. Because the CS model nevertheless outperforms the IW model, this is not an issue for this study, and it may be indicative of a similar *K* among clades, which is in line with ecotype space limiting the number of species equally in each clade. Still, models with different *K* values for each clade could in principle be fitted to the data to confirm this. In practice, however, this is not really feasible, because we are already estimating four or five parameters, and there may not be enough information in a data set of this size to allow for more parameters to be reliably estimated.

We have only considered two models of diversity-dependence: one where diversity-dependence only applies to species within the same clade, and one where it also applies to species of other clades establishing on an island. Various other models can be conceived. For example, diversity-dependence might apply to a higher taxonomic level, i.e. the number of clades, rather than the number of species within them may be limiting further colonization or diversification. The effect of phylogenetic relatedness may also differ for speciation, extinction and colonization. For instance, Pires et al. (2017) found that speciation is mostly affected by within-clade diversity-dependence, whereas extinction is mostly affected by between-clade diversity-dependence. Furthermore, one could define phylogenetic limits in terms of actual phylogenetic distances so that we move from a within- and between-clade dichotomy to a more continuous spectrum where some phylogenetically related clades may interact, but more distantly related clades do not. There are no likelihood methods for such models yet. One may have to resort to simulation-based approaches such as Approximate Bayesian Computation (Janzen et al., 2015). These methods need to integrate over all possible trajectories of the clades through time which is not trivial, because the space of these trajectories is extremely high-dimensional.

One may wonder what it is in the branching pattern that allows for selecting one model over the other. An intuitively obvious candidate may be the rank of the largest clade. The IW model can be expected to have the first clade as the largest, because later clades will be suffering from diversity-dependence and hence not be able to grow very large. However, we noticed that the first clade is also almost equally often the largest clade under the CS model (Fig. 2). Hence, a pattern we may put down to incumbency and interclade competition (Silvertown, 2004; Schenk et al., 2013) arises equally prominently under a model without interclade competition. Apparently, in our empirical example time since colonization is a more important determinant of the size of a clade than diversity-dependence. The IW or CS nature of the colonization and diversification process, and thus the presence or absence of priority effects at the macroevolutionary scale, is hidden in a more complex way in the phylogenetic branching pattern. However, we note that the estimate of the island-wide carrying capacity *K* (132) is quite a bit larger than the number of species present on the island (66), suggesting that the island is still far from saturation under the IW model. Other systems may have a lower *K* and the effect of priority effects may be relatively stronger. It is an interesting avenue to study whether there is a relationship between the magnitude of these priority effects and invasibility of islands, which may contribute to our understanding of biological invasions (Fraser et al., 2015).

The new IW model may also be applicable in other fields, such as epidemiology where it may serve as a tool in understanding spread of an infectious disease, e.g. a virus, in island-like systems such as schools or hospitals. In such systems there may be multiple sources of infections which spread through the local population which can be modelled as colonizations. The carrying capacity is the number of children or patients. The IW model would be the appropriate model if once the host is infected, it builds up immunity against all strains, thus hindering further colonization and diversification. This scenario is most likely if the colonizing strains are phylogenetically related. The CS model would be a better description if a host can be infected by multiple strains, but within each strain there is viral interference (see e.g. Ojosnegros et al. (2010)). This scenario is most likely if the colonizing strains are phylogenetically (and hence functionally) distinct.

We have provided a new model for island biogeography with diversity-dependent feedback on colonization and diversification occurring between all island species. Our implementation of this IW model has some computational limitations for islands with large numbers of colonizations, particularly if these are non-endemic, but typical insular data sets with a moderate number of colonizations or a high level of endemism (such as our *Eleutherodactylus* frogs) are perfectly feasible. Our single empirical example serves as an illustration to the empiricist how to explore the phylogenetic limits of ecological limits to diversification. Although this example showed a clearly better fit of the CS model, future applications may identify the IW model as the superior model indicating feedback between all colonizing lineages, and hence priority effects.

## Supporting information

Figures Tables and Supplementary Data and Scripts

## Acknowledgements

We thank Ally Phillimore for early discussions that provided the original motivation for this study. We thank Pedro Neves, Joshua Lambert, Ally Phillimore, Tanja Stadler and two anonymous reviewers for helpful comments on the manuscript, and Hanno Hildenbrandt for a major improvement in the performance of the DAISIE PACKAGE. We thank the Center for Information Technology of the University of Groningen for their support and for providing access to the Peregrine high performance computing cluster. RSE and LV thank the Netherlands Organization for Scientific Research (NWO) for financial support through NWO-VICI and NWO-VIDI grants, respectively. LV thanks the DFG for an Eigene Stelle grant (VA 1102/1-1). BH was supported by the Laboratoires d’Excellences (LABEX) TULIP (ANR-10-LABX-41). CV thanks the Spanish Government for funding project CGL2016-75227-P within which the data were collected. RSE and BH thank the bilateral French-Dutch Van Gogh Program for funding work visits.

## Supplementary Material

### Supplementary Text – Derivation of likelihood computation

We derive the likelihood of colonization and branching data using the *Q*-approach (Etienne et al., 2012; Laudanno et al., 2019). This approach is based on the quantity *Q*(*t*), defined as the probability that a model realization is consistent with the data from the initial time *t*_i_ to a given time *t*. We construct the differential equations governing the dynamics of *Q*(*t*), and explain how the likelihood can be extracted from *Q*(*t*_p_) at the present time *t*_p_.

A dataset consists of a number of trees, each associated with a mainland species. An example is shown in Fig. S5. Several trees can be associated with the same mainland species, if it has colonized the island several times (e.g., species B in the figure). For trees consisting of a single branch, the dataset also specifies whether speciation has taken place, i.e., whether the species at the present time is endemic to the island (compare species C and D).

To set up the likelihood computation, we partition the number of extant species on the island at any time *t* as follows:

> *k* the number of species present on the island at time *t* that are represented in the dataset, i.e., that survive until the present and are sampled;
>
> *m* the number of mainland species present on the island at time *t* that are not in the data (either because the lineage became extinct before the present or because it has not been sampled);
>
> *e* the number of endemic species at time *t* that are not in the data

Hence, the total number of extant species at time *t* is equal to *k* + *m* + *e*. We have *m ⩽M*, where *M* is the number of species in the mainland source pool. Note that *k* is determined by the data, while *m* and *e* are not. Instead they vary between model realizations, and are part of the system state in the *Q*-approach.

Fig. S6 shows one possible model realization consistent with the dataset of Fig. S5, together with the values that *k, m* and *e* take through time.

### Before first colonization recorded in data

Before any colonization event of still extant lineages (so *k* = 0) we keep track of the number of non-endemic (i.e., mainland) species *m* and the number of endemic species *e*. The dynamics of the probabilities *Q*_*m,e*_(*t*), where *m* and *e* can take any integer value, are described by the following set of differential equations:

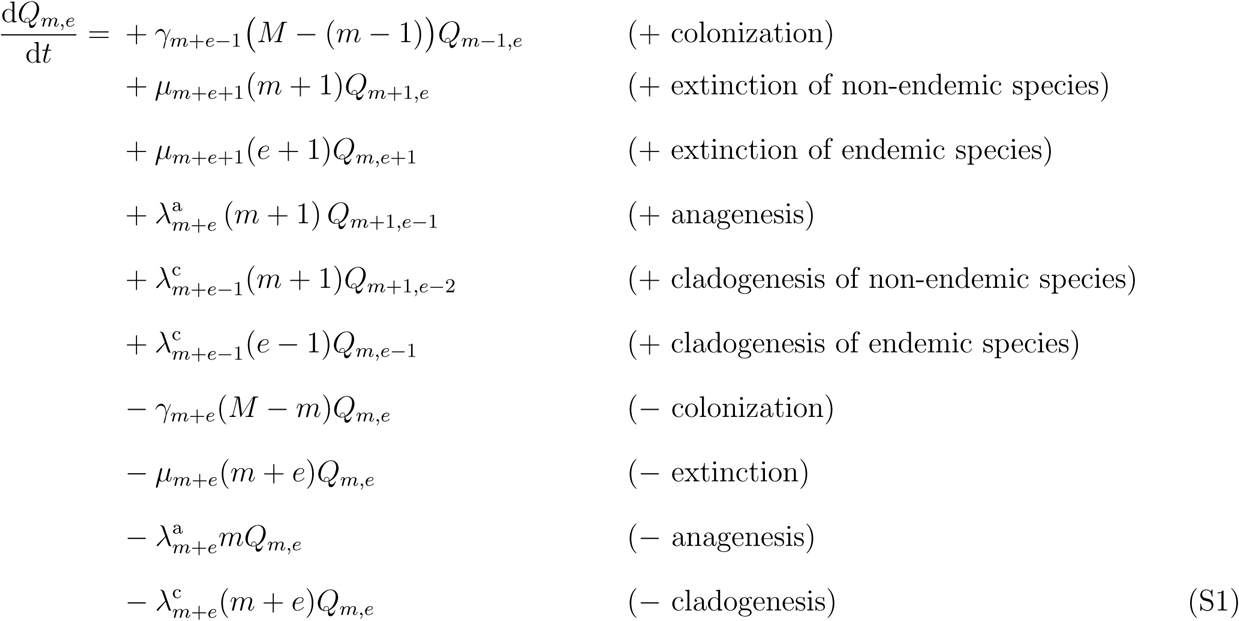

The positive terms on the right-hand side correspond to events that bring the system in the state with *m* non-endemic and *e* endemic species, and hence increase the probability *Q*_*m,e*_. These events are due to colonization, extinction of a non-endemic species, extinction of an endemic species, anagenesis (of a non-endemic species) and cladogenesis in a non-endemic and endemic species. The negative terms correspond to events that decrease the probability due to the same processes when in the state with *m* non-endemic and *e* endemic species, i.e., events that move the system away from this state.

The initial condition for *Q*_*m,e*_(*t*) at time *t*_i_ usually corresponds to an empty island, so

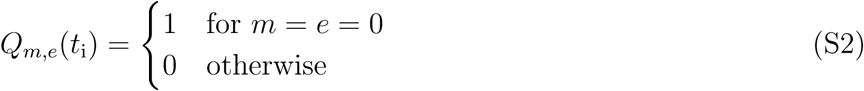

Note that because *k* = 0, there are no lineages yet with which the realizations have to be consistent, and hence equations (S1) are identical to the master equations for the probabilities *P*_*m,e*_(*t*) to have *m* non-endemic and *e* endemic species.

### After first colonization recorded in data

At the first colonization event recorded in the data, which we call *t*_1_, one of the extant species is represented in the phylogeny (so *k* = 1). From this time onwards, *Q*_*m,e*_(*t*) becomes different from *P*_*m,e*_(*t*), because the realizations contributing to *Q*_*m,e*_(*t*) not only have *m* non-endemic species and *e* endemic species at time *t*, but should also be consistent with the colonization at time *t*_1_.

After the first colonization event, recolonization of the same mainland species is not allowed until the colonizing species undergoes speciation, because it would change the data, namely the time of colonization. This implies that we have to keep track of whether the colonizing species has undergone speciation. We therefore expand the set of equations (S1) to two sets. The first set describes the probabilities where speciation has not (yet) taken place. The probabilities in this set are denoted by 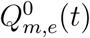, where the superscript 0 indicates that speciation has not (yet) taken place. The second set describes the probabilities where speciation has taken place and recolonization of the same species can therefore occur without changing the data. The probabilities in this set are denoted by 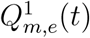, where the superscript 1 indicates that speciation has taken place.

The dynamical equations for 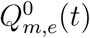 are

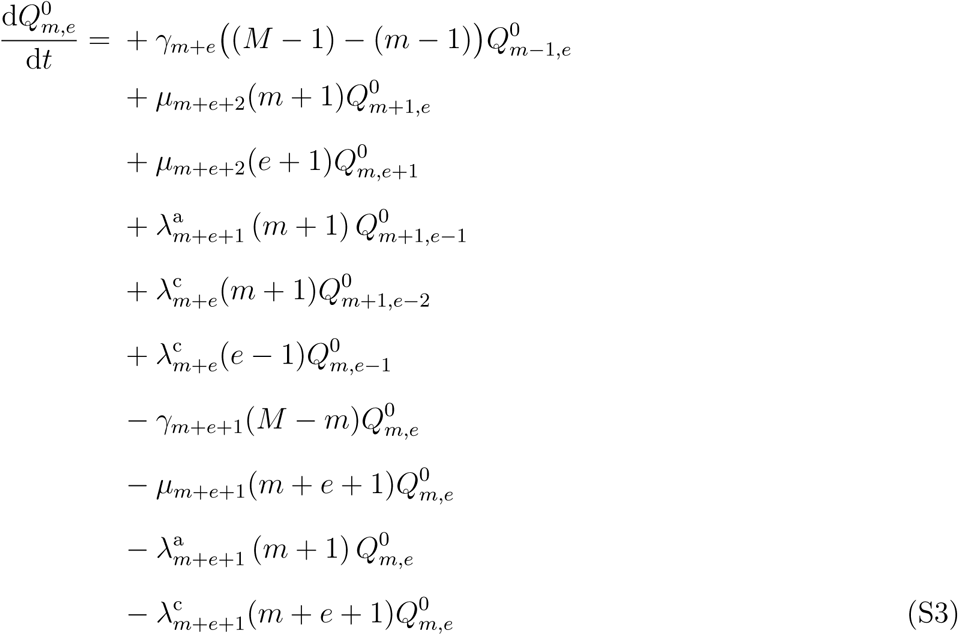

Importantly, these equations guarantee that the realizations contributing to 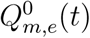 are consistent with a colonization event at time *t*_1_ in the data. In particular, the mainland species that has colonized does not recolonize (first line, factor (*M −* 1) *−* (*m −* 1) instead of *M −* (*m −* 1)) and the colonizing species does not become extinct (second line, factor *m* + 1 instead of *m* + 2). If the colonizing species undergoes anagenesis or cladogenesis, 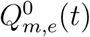 does not increase (fourth and fifth line, factor *m* + 1), but 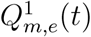 increases (see below).

The dynamical equations for 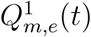 are

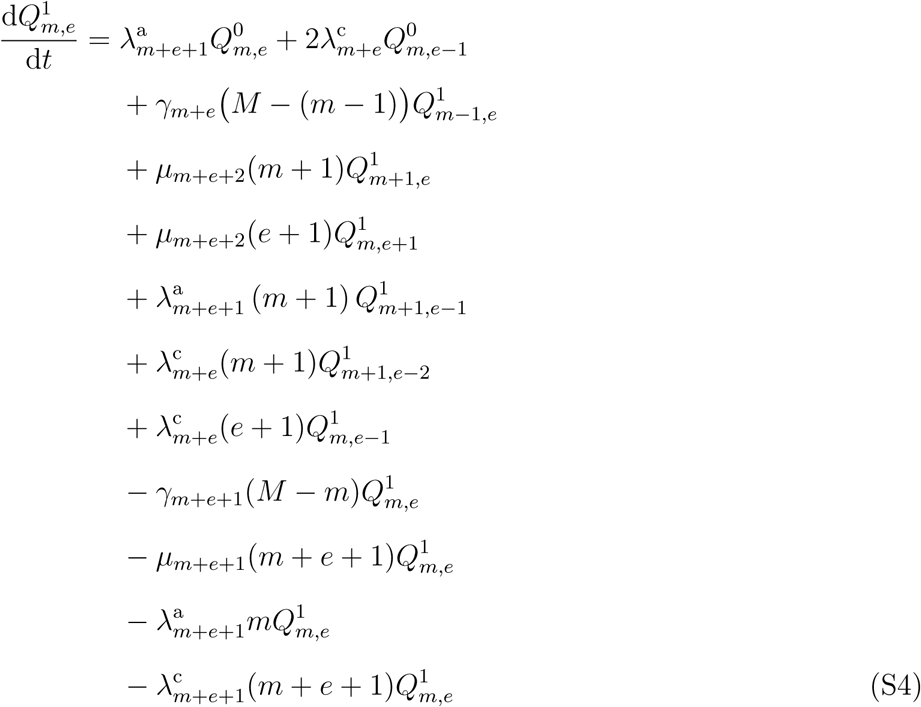

The first line on the right-hand side corresponds to a speciation of the colonizing species, either anagenesis or cladogenesis. Observe the factor 2 in the cladogenesis term. This is a combinatorial factor required in the *Q*-approach, which can be understood intuitively by noting that either of the two daughter species can be represented in the phylogeny (see Laudanno et al. (2019) for a detailed justification).

Equations (S4) guarantee that the lineage that originates at colonization time *t*_1_ does not become extinct (fourth line, factor *e* + 1 instead of *e* + 2). In contrast to equations (S3), the mainland species that colonized is allowed to recolonize (because it does not change the data; second line, factor *M −* (*m −* 1)). The term corresponding to cladogenesis of endemic species (seventh line) has a factor *e* + 1, because there are *e* endemic species in total and the species represented in the phylogeny (i.e., the species that descends from the colonization event at time *t*_1_) should be counted twice, due to the combinatorial factor 2 we mentioned above.

The initial condition for equations (S3–S4) at time *t*_1_ impose the colonization of the appropriate mainland species,

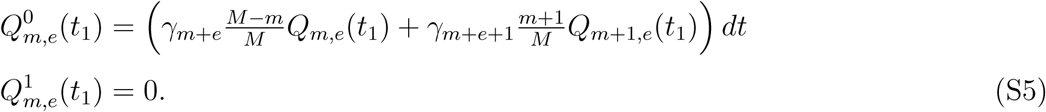

The first term in the brackets corresponds to the case where the colonizing mainland species is not yet present on the island at time *t*_1_; the second term to the case where the species is already present. The infinitesimal d*t* makes clear that we are looking at the probability of colonization at precisely time *t*_1_, i.e., between *t*_1_ and *t*_1_ + d*t*. The conditions (S5) connect equations (S1) before and equations (S3–S4) after the colonization event.

### After second colonization recorded in data

Suppose the next event in the data is a new colonization of another mainland species, which occurs at time *t*_2_. Again, we have to keep track of whether this colonizing species has undergone speciation. We proceed as before, and expand the set of equations (S3–S4) to two sets: a first set for the probabilities where speciation has not (yet) taken place, and a second set for the probabilities where speciation has taken place. This leads to four sets of dynamical variables, 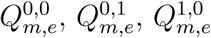 and 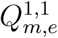, where the first and second bit indicate the speciation status of the first and second colonization, respectively.

The corresponding dynamical equations are

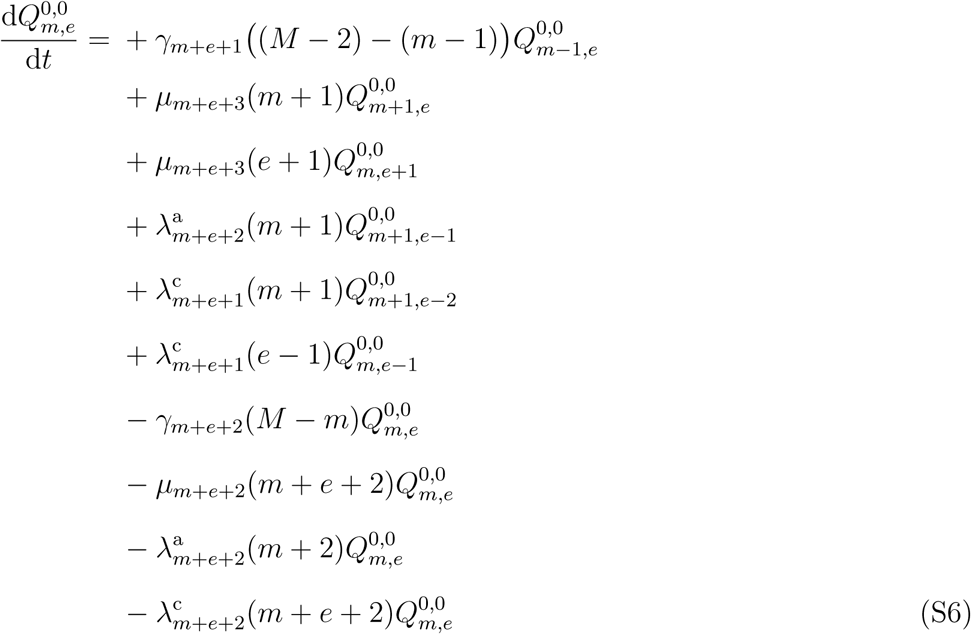

where we exclude the recolonization of the two colonizing mainland species (first line, factor (*M −* 2) *−* (*m −* 1) instead of *M −* (*m −* 1));

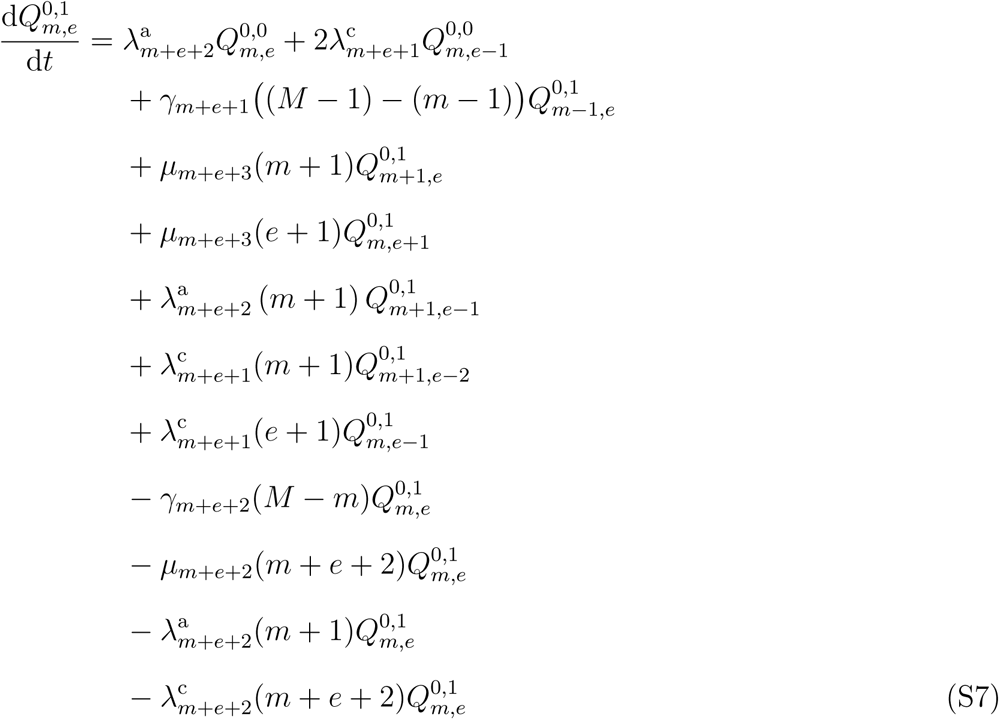

where we exclude the recolonization of the first colonizing mainland species;

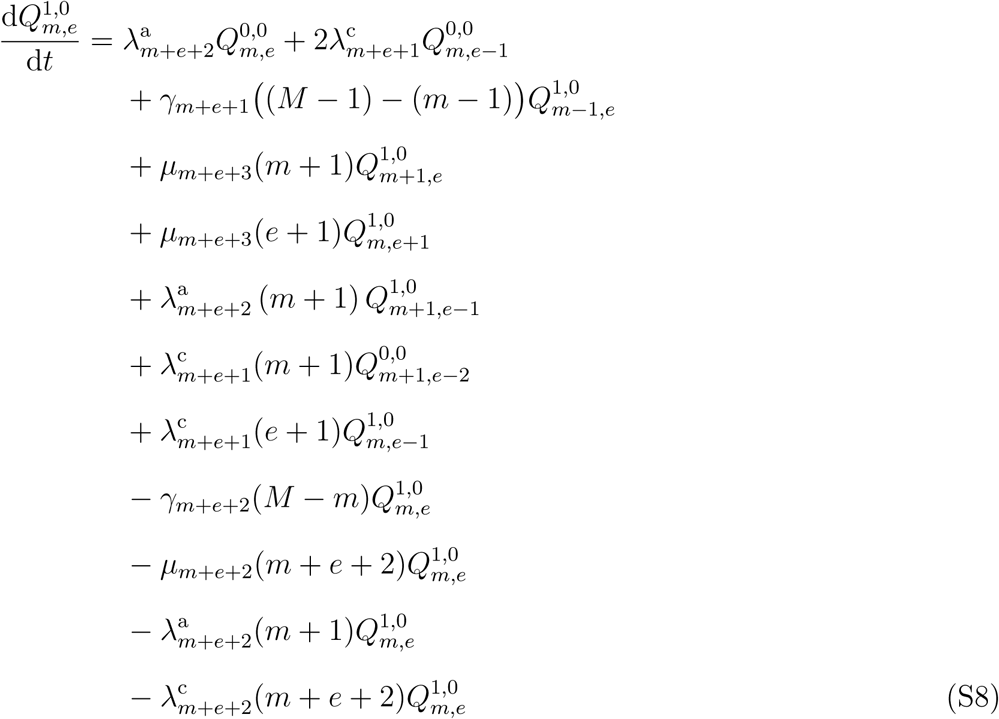

where we exclude the recolonization of the second colonizing mainland species;

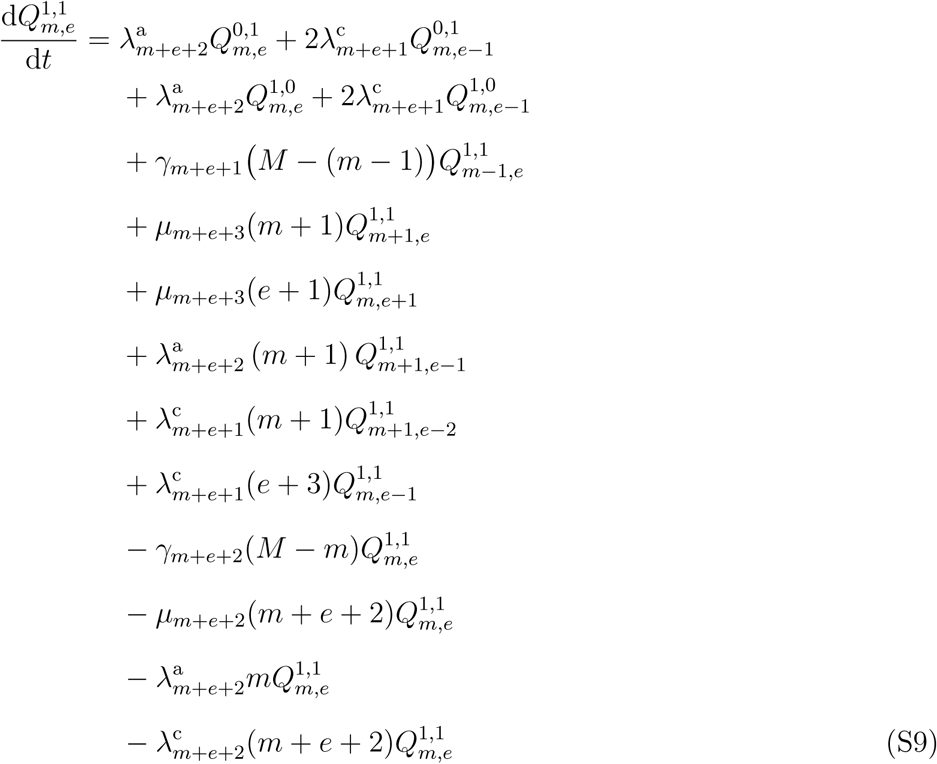

Note the factor *e* + 3 in the term corresponding to cladogenesis of endemic species (eighth line): there are *e* + 1 endemic species and the two species represented in the phylogeny (the species descending from the two colonization events) should be counted twice.

The initial condition at colonization time *t*_2_ connects the solutions before (equations (S3–S4)) and after (equations (S6–S9)) the colonization event,

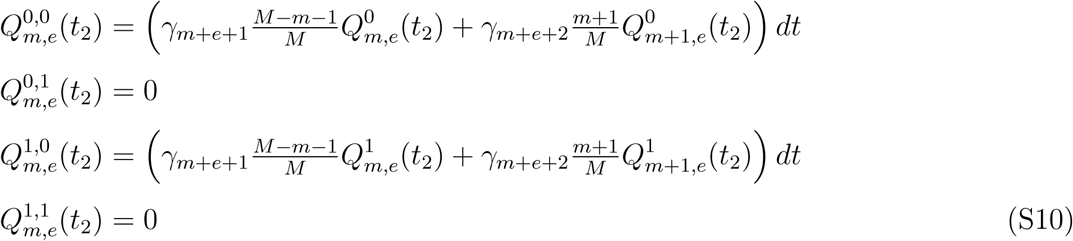

### After 𝓁-th colonization recorded in data

Now we generalize equations (S3–S4) and (S6–S9) to an arbitary number of colonization events recorded in the data. We define the number *𝓁* as the number of colonizations from which originate the species at time *t* represented in the phylogeny (see Fig. S6 for an illustration). Recall that there are *k* such species, so we have *𝓁 ⩽ k*.

For each of the *𝓁* colonizations we have to keep track of whether speciation has taken place in the corresponding lineage (see previous sections for the cases *𝓁* = 1 and *𝓁* = 2). Therefore, we introduce a binary vector 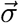 of length *𝓁*, the *i*-th component of which indicates whether speciation has taken place (*σ*_*i*_ = 1) or not (*σ*_*i*_ = 0) in the lineage that originates from the *i*-th colonization event. We add this vector to the system state, so that it consists of the number of non-endemic species *m*, the number of endemic species *e* and the speciation state 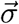 of the *𝓁* colonization events. We denote the corresponding dynamical variable by 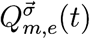.

We need some additional notation to formulate the dynamical equations. We denote by *𝓁*^0^ the number of colonizations recorded in the data for which speciation has not (yet) taken place, equal to the number of zeros in 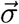 (see Fig. S6 for an illustration). For a given vector 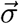, we define 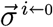 as the vector with the same components except *σ*_*i*_ = 0, and 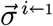 as the vector with the same components except *σ*_*i*_ = 1. The vector 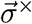 stands for the vector with the last component removed (i.e., a vector of length *𝓁 −* 1). For example, for 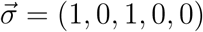, we have 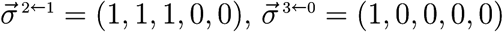 and 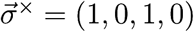.

Then, the dynamical equations are

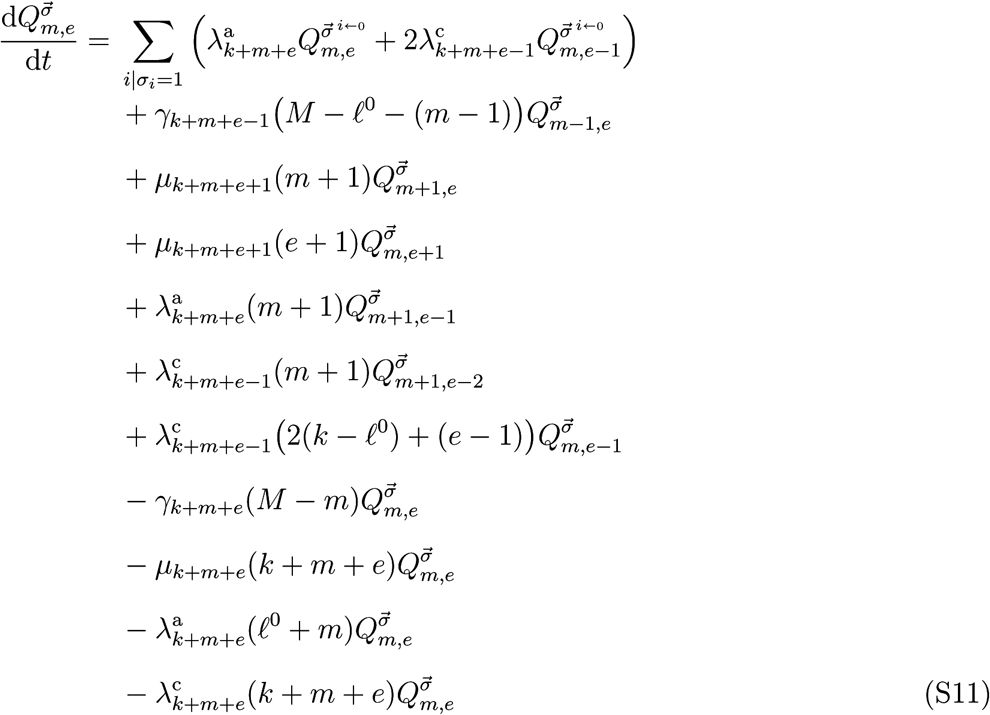

It can be verified that these equations generalize equations (S3–S4) and equations (S6–S9). In particular, on the second line we exclude the recolonization of the *𝓁*^0^ mainland species that have not yet undergone speciation on the island. On the seventh line, when counting the endemic species that can undergo cladogenesis, we count twice the *k − 𝓁*^0^ endemic species that are represented in the phylogeny.

Equations (S11) govern the dynamics in the time interval where *k*, the number of species in the data, and *𝓁*, the number of colonizing lineages, take fixed values. This time interval starts with the *k*-th event in the data, at time *t*_*k*_ (note that *k* increases by one at each event in the data, so that we can use *k* to count the events). This event is either a new colonization recorded in the data, or a cladogenesis in a lineage present in the data that colonized at an earlier time. To formulate the initial conditions, we distinguish the following cases:

- The event at time *t*_*k*_ is a colonization of a new mainland species, that is, a mainland species that has no colonization in the data before *t*_*k*_. The initial condition, connecting the solutions of equations (S11) before and after *t*_*k*_, is

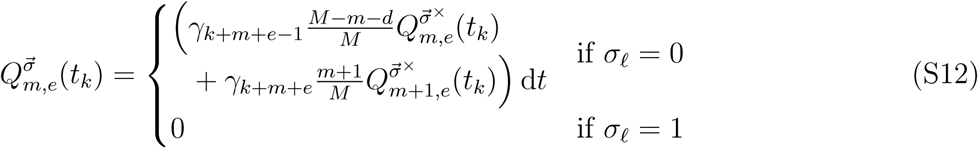

where *d* is the number of mainland species that have at least one colonization in the data before *t*_*k*_. Note that the values of *k* and *𝓁* increase by one at a colonization; the values used in equation (S12) are those after the colonization event.
- The event at time *t*_*k*_ is a colonization of a mainland species that is already present in the data (i.e., it has one or more colonizations in the data before *t*_*k*_), and the lineage initiated at the last colonization of this mainland species before *t*_*k*_ has not branched before *t*_*k*_. Denote the index of the latter colonization by *i*. We have to require that the lineage initiated at the *i*-th colonization has undergone speciation before *t*_*k*_ (otherwise the new colonization would modify the time of the *i*-th colonization). That is, we require that *σ*_*i*_ = 1. Hence, the initial condition is

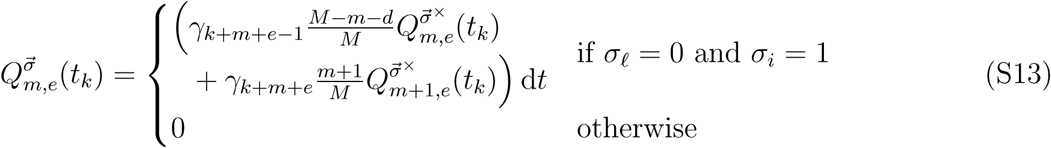
- The event at time *t*_*k*_ is a colonization of a mainland species that is already present in the data, and the lineage initiated at the last colonization of this mainland species before *t*_*k*_ has at least one branching event before *t*_*k*_. Denoting the index of the latter colonization by *i*, we know from the presence of a branching event that immediately before *t*_*k*_ the corresponding component *σ*_*i*_ = 1 (more precisely, 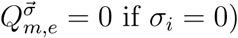. Hence, the initial condition is still given by equation (S12).
- The event at time *t*_*k*_ is a cladogenesis. Denoting by *i* the colonization that initiated the clade in which this cladogenesis event occurs, the initial condition is

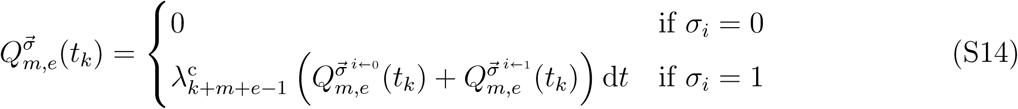

We see that the two sets of equations for *σ*_*i*_ = 0 and *σ*_*i*_ = 1 are merged (or collapsed) into one set of equations for *σ*_*i*_ = 1. Note that at a cladogenesis event the value of *𝓁* does not change, but the value of *k* increases by one.

From a computational point of view it is important to note that we do not have to track probabilities 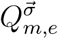 for all possible values of the vector 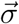. First, component *σ*_*i*_ is only defined from the *i*-th colonization time onwards (undefined values are indicated by *×*-marks in the table of Fig. S6). Second, for a colonization that initiates a clade with branching events, the corresponding component *σ*_*i*_ has to be kept only until the first branching event. After this first branching, component *σ*_*i*_ is identical to one (see *σ*_A_ and 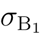 in Fig. S6; the numbers 1 printed in grey are imposed by the data). Third, for a colonization that initiates a branch that remains non-endemic until the present time, the corresponding component *σ*_*i*_ can be discarded entirely, because it is identical to zero from the *i*-th colonization time to the present time *t*_p_ (see 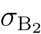 and *σ*_C_ in Fig. S6; the numbers 0 printed in grey are imposed by the data).

### Likelihood from solution of Q-equation

We continue the above procedure, until reaching the present time *t*_p_. The likelihood can then be extracted from components 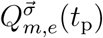 at the present time *t*_p_.

To do so, we assume that we have complete information about the extant species. That is, we assume that there are no other species on the island at the present time than those represented in the dataset. This allows us to set the number of missing non-endemic and endemic species to zero, i.e., we impose that *m* = *e* = 0 at time *t*_p_. The data also allow us to determine the vector 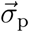 at time *t*_p_. We set *σ*_*i*_ = 0 for colonizations that correspond to lineages that are still non-endemic at the present time (represented by dashed lines in Fig. S5). For all other colonizations we set *σ*_*i*_ = 1. Finally, the likelihood is given by 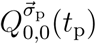.

In practice we often do not have complete information. For example, there might be large uncertainty about colonization times, or there might be missing species at the present time, i.e., species that are not represented in the dataset. Some types of missing information can be easily dealt with, while other types requires a more thorough adaptation of the computational procedure.

**Figure S1.**
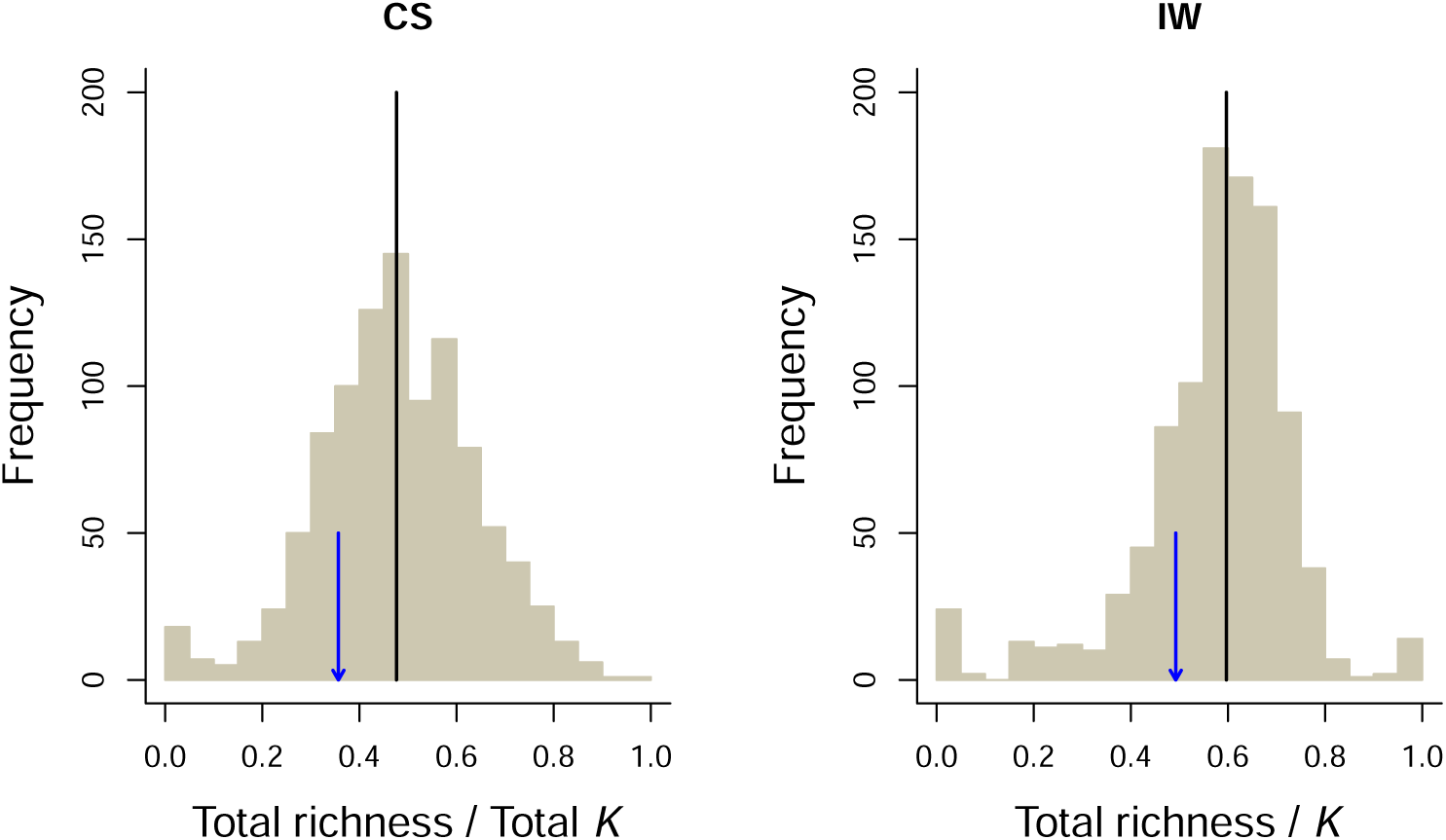
Histograms showing the distribution of different saturation metrics from data sets simulated under CS (left) and IW (right). The black line shows the median value. The blue arrow points to the empirical value (i.e. the total number of species in the empirical data set divided by the estimated value of the carrying capacity. In the CS case the carrying capacity is the number of colonizations multiplied by the estimated clade-specific *K*, in the IW case it is just the estimated island-wide *K*.

**Figure S2.**
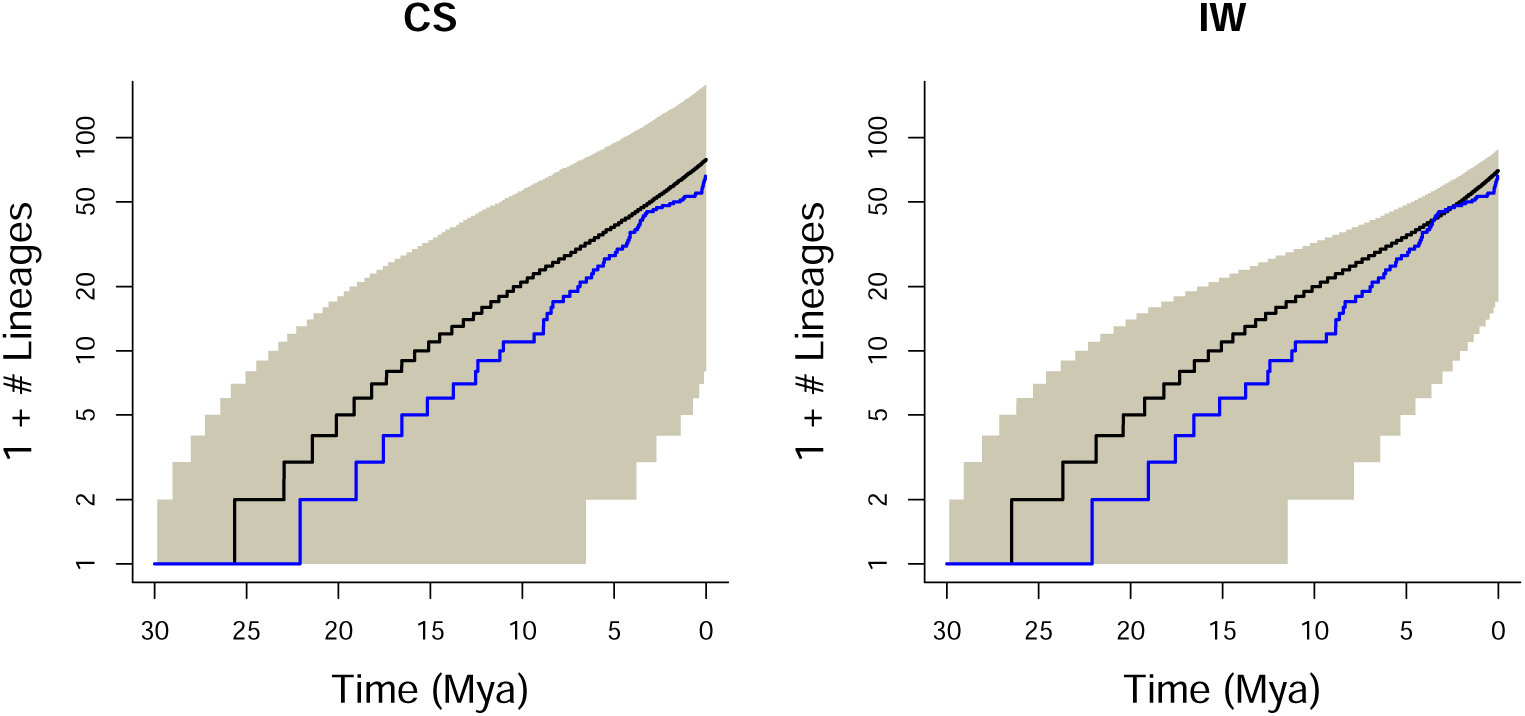
Number of species through time for 5,000 data sets simulated under CS (left) and IW (right). The black line shows the median value, and the shaded area the 95% confidence interval. The blue line shows the empirical curve.

**Figure S3.**
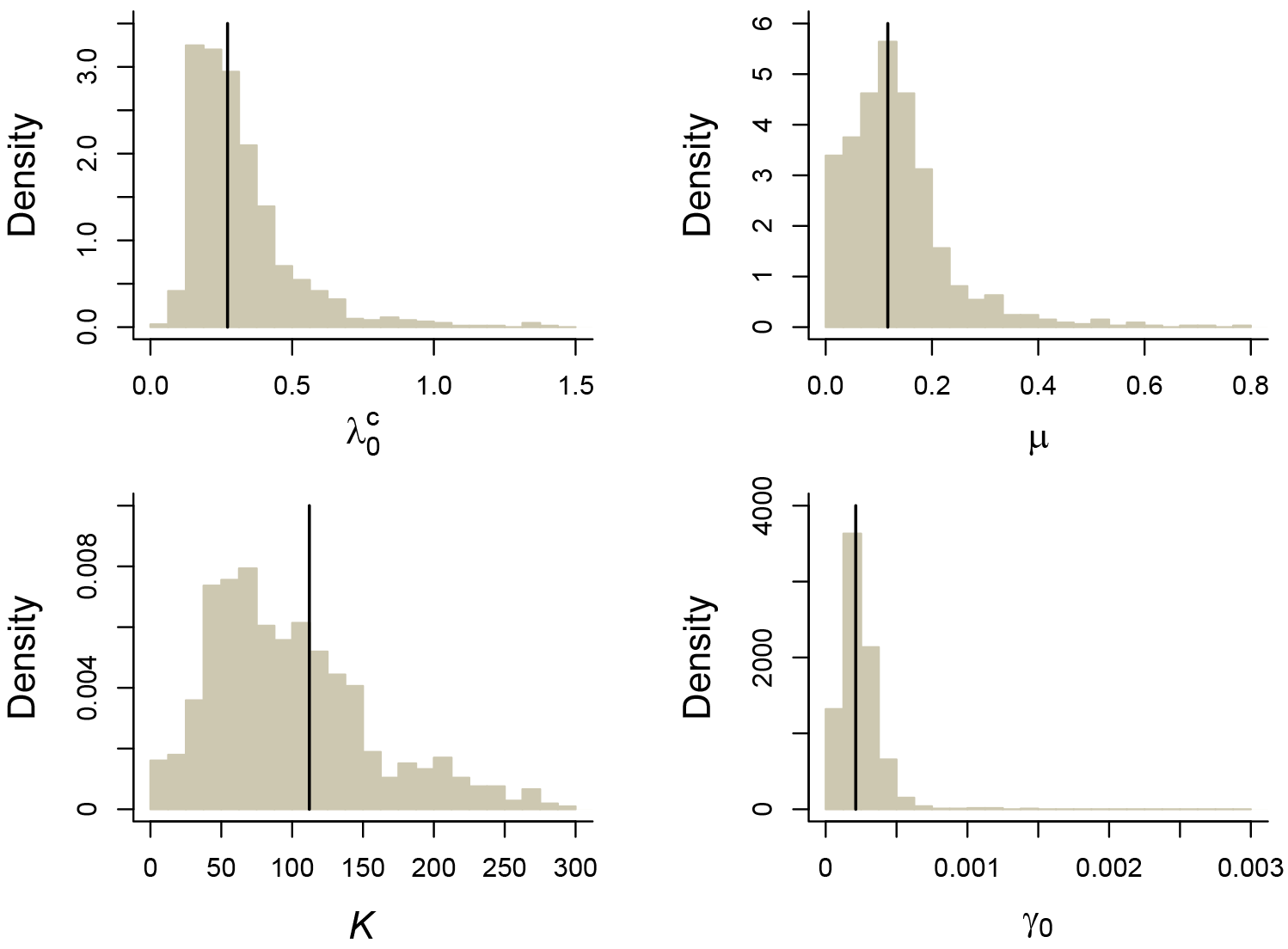
Histograms showing the CS model parameters fitted to IW model simulations. The black line shows the median value. There are no true values to compare with, because the generating and inference model differ. Note that the *K* estimated by the CS model (i.e. this is a *K* per clade) under IW simulations is approximately three times higher than that estimated under CS simulations (see Fig. 3). IW simulations (with a generating island-wide *K* of 132) will form clades with a larger variation in clade size than CS simulations (see Fig. 2), and this is best fitted by a higher clade-level *K* in the CS model.

**Figure S4.**
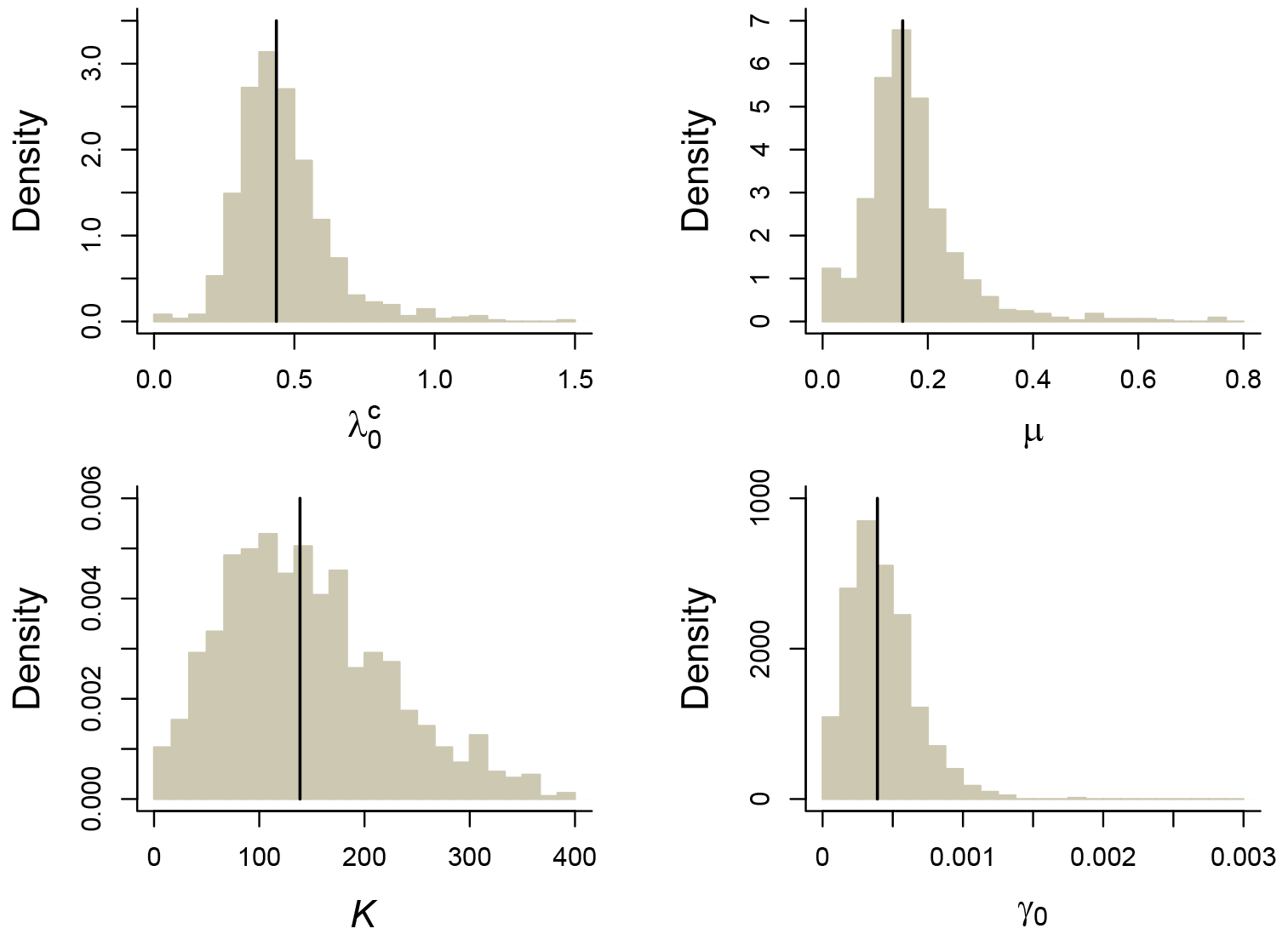
Histograms showing the IW model parameters fitted to CS model simulations. The black line shows the median value. There are no true values to compare with, because the generating and inference model differ.

**Figure S5.**
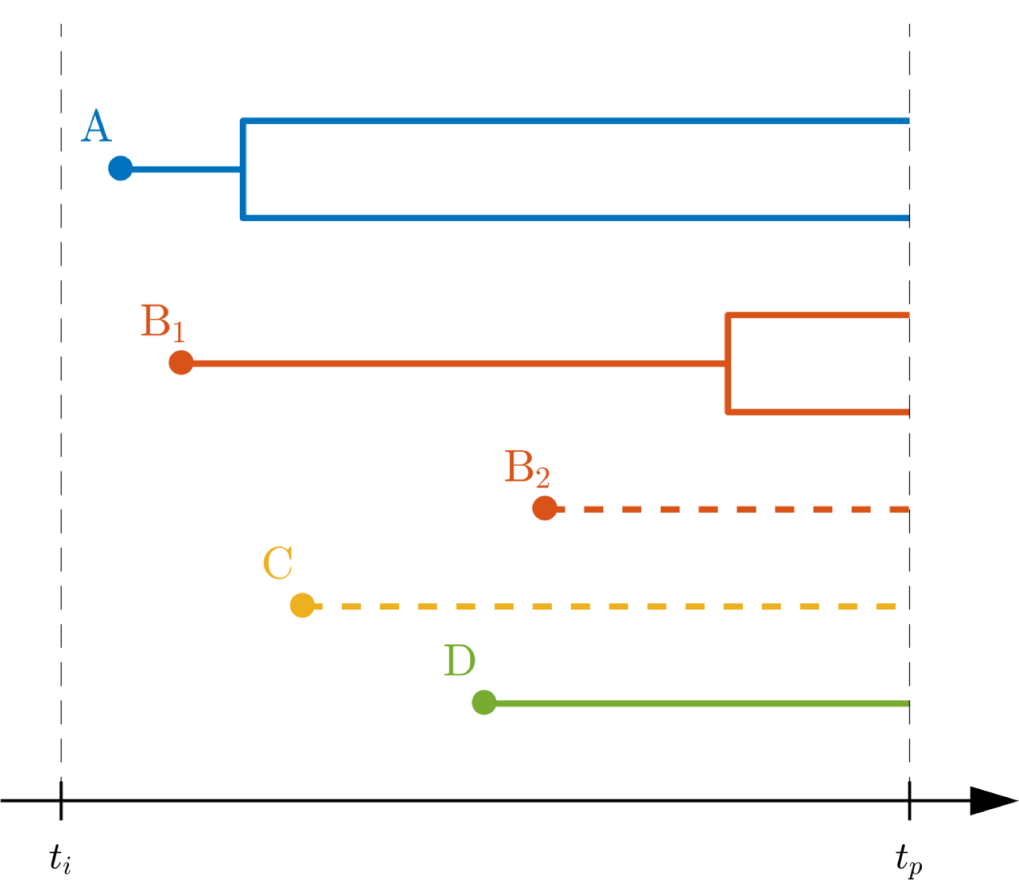
Example data set for the likelihood computation. Different colors represent trees associated with different mainland species (here species A, B, C and D). Filled circles indicate colonization events. Note that mainland species B has two colonization events in the data set; we distinguish the two corresponding trees with a subscript, B_1_ and B_2_. Full lines correspond to trees in which the colonizing mainland species has speciated on the island; dashed lines indicate lineages that have not yet speciated and are still non-endemic at the present time. Time *t*_*i*_ is the emergence time of the island; time *t*_p_ is the present time.

**Figure S6.**
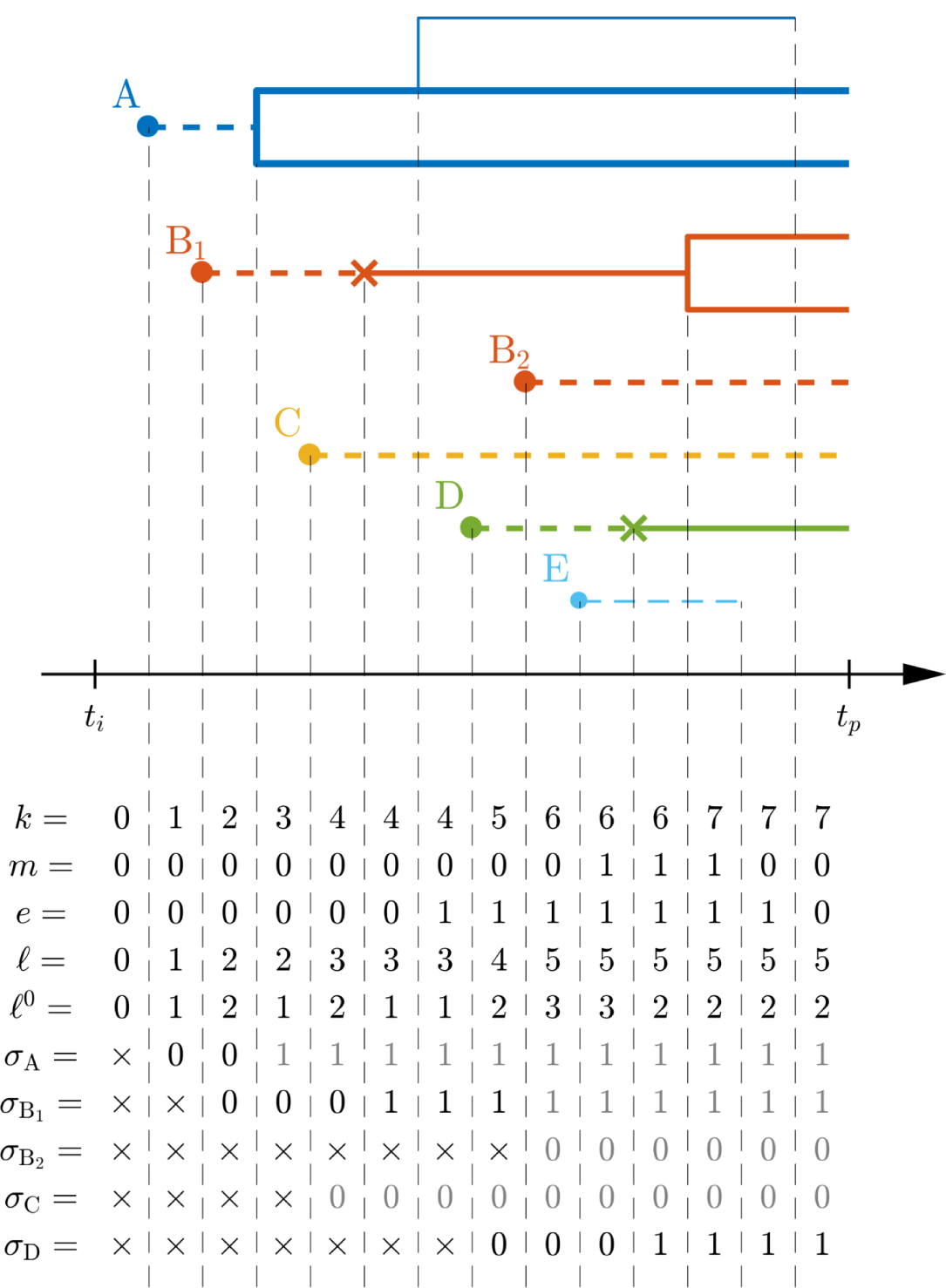
A full set of events leading to the data set represented in Fig. S5. Full lines indicate endemic species; dashed lines indicate non-endemic species. Thick lines correspond to species that are represented in the data set; thin lines correspond to lineages that become extinct before the present (and hence are not part of the data). Filled circles are colonizations; *×*-marks are anagenesis events. The lower part of the figure shows the values the different variables take during the likelihood computation, from the initial time *t*_i_ to the present time *t*_p_.

## Supplementary files

The following files are available at Dryad at doi:10.5061/dryad.t1g1jwt3n.

- Table S1 – File `Table S1.xlsx` – List of missing species added to the phylogenetic trees and justification for the topological assignation.
- Table S2 – File `out_empirical_M1000.txt` – Parameter estimates of the CS and IW model for the 100 sets of trees with added missing species.
- Table S3 – File `out_simulation_M1000.txt` – Parameter estimates of the CS and IW model for 1000 simulated sets of trees.
- `Frogs_Alignment.nex` – DNA concatenated sequence alignment.
- `Frogs_BEAST2_script.xml` – XML script to perform phylogenetic reconstruction in BEAST2.
- `100trees_with_missing_species_added.tre` – One hundred trees with missing species added in Newick format. The 52nd tree is used for bootstrap analyses.
- `Frogs_100_30_miss_spec_added_M1000.Rdata` – One hundred sets of trees with missing species added as an R object following the DAISIE data format. The 52nd tree is used for bootstrap analyses.
- `CSvsIW_bootstrap.R` – R script to fit models to simulated and empirical data.
- `Frogs_sim_5000.R` – R script to simulate data sets.
- `out empirical M1000.txt`– Parameter estimates of the CS and IW model for the 100 sets of trees with added missing species.
- `out_simulation_M1000.txt`– Parameter estimates of the CS and IW model for 1000 simulated sets of trees.
- `Plot_Goodness_of_fit_frogs.R` – R script to plot the figures related to the goodness-of-fit of the CS and IW models to the empirical data.
- `Plot_CSvsIW.R` – R script to plot the figures related to the inference of the CS and IW models.

## References

Cadotte, M. W., T. J. Davies, and P. Peres-Neto. 2017. Why phylogenies do not always predict ecological differences. Ecological Monographs 87:535–551.

Cahill, J. F., S. W. Kembel, E. G. Lamb, and P. A. Keddy. 2008. Does phylogenetic relatedness influence the strength of competition among vascular plants? Perspectives in Plant Ecology, Evolution and Systematics 10:41–50.

Darwin, C. 1859. On the origin of species by means of natural selection, or preservation of favoured races in the struggle for life. John Murray, London.

Dugo-Cota, A., C. Vilá, A. Rodríguez, and A. Gonzalez-Voyer. 2019. Ecomorphological convergence in Eleutherodactylus frogs: a case of replicate radiations in the Caribbean. Ecology Letters 22:884–893.

Etienne, R. S., B. Haegeman, T. Stadler, T. Aze, P. N. Pearson, A. Purvis, and A. B. Phillimore. 2012. Diversity-dependence brings molecular phylogenies closer to agreement with the fossil record. Proceedings of the Royal Society of London B 279:1300–1309.

Etienne, R. S., A. L. Pigot, and A. B. Phillimore. 2016. How reliably can we infer diversity-dependent diversification from phylogenies? Methods in Ecology and Evolution 7:1092–1099.

Foote, M., R. A. Cooper, J. S. Crampton, and P. M. Sadler. 2018. Diversity-dependent evolutionary rates in early Palaeozoic zooplankton. Proceedings of the Royal Society of London B: Biological sciences 285:20180122.

Foote, M. and A. I. Miller. 2006. Principles of psaleontology. W.H. Freeman, New York.

Fraser, C. I., S. C. Banks, and J. M. Waters. 2015. Priority effects can lead to underestimation of dispersal and invasion potential. Biological Invasions 17:1–8.

Gerhold, P., J. F. C. Jr, M. Winter, I. V. Bartish, and A. Prinzing. 2015. Phylogenetic patterns are not proxies of community assembly mechanisms (they are far better). Functional Ecology 29:600–614.

Germain, R. M., J. T. Weir, and G. Benjamin. 2016. Species coexistence: macroevolutionary relationships and the contingency of historical interactions. Proceedings of the Royal Society B: Biological Sciences 283:20160047.

Heinicke, M. P., W. E. Duellman, and S. B. Hedges. 2007. Major Caribbean and Central American frog faunas originated by ancient oceanic dispersal. Proceedings of the National Academy of Sciences 104:10092–10097.

HilleRisLambers, J., P. B. Adler, W. S. Harpole, J. M. Levine, and M. M. Mayfield. 2012. Rethinking community assembly through the lens of coexistence theory. Annual Review of Ecology, Evolution, and Systematics 43:227–248.

Iturralde-Vinent, M. A. 2006. Meso-cenozoic Caribbean paleogeography: Implications for the historical biogeography of the region. International Geology Review 48:791–827.

Jablonski, D. 2008. Biotic interactions and macroevolution: Extensions and mismatches across scales and levels. Evolution 62:715–739.

Janzen, T., S. Höhna, and R. S. Etienne. 2015. Approximate bayesian computation of diversification rates from molecular phylogenies: introducing a new efficient summary statistic, the nltt. Methods in Ecology and Evolution 6:566–575.

Jønsson, K. A., P.-H. Fabre, S. A. Fritz, R. S. Etienne, R. E. Ricklefs, T. B. Jørgensen, J. Fjeldså, C. Rahbek, P. G. P. Ericson, F. Woog, E. Pasquet, and M. Irestedt. 2012. Ecological and evolutionary determinants for the adaptive radiation of the Madagascan vangas. Proceedings of the National Academy of Sciences 109:6620–6625.

Laudanno, G., B. Haegeman, and R. S. Etienne. 2019. Additional analytical support for a new method to compute the likelihood of diversification models. bioRxiv.

MacArhur, R. H. and E. O. Wilson. 1967. Island Biogeography. Princeton University Press, Princeton, NJ.

Matzke, N. J. 2013. Probabilistic historical biogeography: new models for founder-event speciation, imperfect detection, and fossils allow improved accuracy and model-testing. Frontiers of Biogeography 5:242–248.

Matzke, N. J. 2014. Model Selection in Historical Biogeography Reveals that Founder-Event Speciation Is a Crucial Process in Island Clades. Systematic Biology 63:951–970.

Mayfield, M. M. and J. M. Levine. 2010. Opposing effects of competitive exclusion on the phylogenetic structure of communities. Ecology Letters 13:1085–1093.

Narwani, A., B. Matthews, J. Fox, and P. Venail. 2015. Using phylogenetics in community assembly and ecosystem functioning research. Functional Ecology 29:589–591.

Ojosnegros, S., N. Beerenwinkel, T. Antal, M. A. Nowak, C. Escarmís, and E. Domingo. 2010. Competition-colonization dynamics in an rna virus. Proceedings of the National Academy of Sciences 107:2108–2112.

Phillimore, A. B. and T. D. Price. 2008. Density-dependent cladogenesis in birds. PLoS Biology 6:0483–0489.

Pigot, A. L. and R. S. Etienne. 2015. A new dynamic null model for phylogenetic community structure. Ecology Letters 18:153–163.

Pires, M. M., D. Silvestro, and T. B. Quental. 2017. Interactions within and between clades shaped the diversification of terrestrial carnivores. Evolution 71:1855–1864.

Proches, S., J. R. U. Wilson, D. M. Richardson, and M. Rejmnek. 2008. Searching for phylogenetic pattern in biological invasions. Global Ecology and Biogeography 17:5–10.

Rabosky, D. L. 2013. Diversity-dependence, ecological speciation, and the role of competition in macroevolution. Annual Review of Ecology, Evolution, and Systematics 44:481–502.

Rabosky, D. L. and R. E. Glor. 2010. Equilibrium speciation dynamics in a model adaptive radiation of island lizards. Proceedings of the National Academy of Sciences 107:22178–22183.

Rabosky, D. L. and I. J. Lovette. 2008. Density-dependent diversification in North American wood warblers. Proceedings of the Royal Society of London B 275:2363–2371.

Raup, D. M., S. J. Gould, T. J. M. Schopf, and D. S. Simberloff. 1973. Stochastic models of phylogeny and the evolution of diversity. Journal of Geology 81:525–542.

Revell, L. J. 2012. phytools: an R package for phylogenetic comparative biology (and other things). Methods in Ecology and Evolution 3:217–223.

Schenk, J. J., K. C. Rowe, and S. J. Steppan. 2013. Ecological Opportunity and Incumbency in the Diversification of Repeated Continental Colonizations by Muroid Rodents. Systematic Biology 62:837–864.

Sepkoski, J. 1996. Competition in macroevolution: the double wedge revisited. Pages 211–255 in Evolutionary Paleobiology (J.L.D. Jablosinki, D.H. Erwin, ed.). University of Chicago Press.

Silvertown, J. 2004. The ghost of competition past in the phylogeny of island endemic plants. Journal of Ecology 92:168–173.

Silvestro, D., A. Antonelli, N. Salamin, and T. B. Quental. 2015. The role of clade competition in the diversification of North American canids. Proceedings of the National Academy of Sciences 112:8684–8689.

Stanley, S. M. 1973. An explanation for cope’s rule. Evolution 27:1–26.

Valente, L., R. S. Etienne, and L.M. Dávalos. 2017a. Recent extinctions disturb path to equilibrium diversity in Caribbean bats. Nature Ecology & Evolution 1:0026.

Valente, L., R. S. Etienne, and J.C. Garcia-R. 2019. Deep macroevolutionary impact of humans on New Zealand’s unique avifauna. Current Biology 29:2563–2569.e4.

Valente, L., J. C. Illera, K. Havenstein, T. Pallien, R. S. Etienne, and R. Tiedemann. 2017b. Equilibrium bird species diversity in Atlantic islands. Current Biology 27:1660–1666.e5.

Valente, L. M., R. S. Etienne, and A. B. Phillimore. 2014. The effects of island ontogeny on species diversity and phylogeny. Proceedings of the Royal Society B: Biological Sciences 281:20133227.

Valente, L. M., A. B. Phillimore, and R. S. Etienne. 2015. Equilibrium and non-equilibrium dynamics simultaneously operate in the Galápagos islands. Ecology Letters 18:844–852.

Valkenburgh, B. V. 1999. Major patterns in the history of carnivorous mammals. Annual Review of Earth and Planetary Sciences 27:463–493.

Violle, C., D. R. Nemergut, Z. Pu, and L. Jiang. 2011. Phylogenetic limiting similarity and competitive exclusion. Ecology Letters 14:782–787.

Walker, T. D. and J. W. Valentine. 1984. Equilibrium models of evolutionary species diversity and the number of empty niches. American Naturalist 124:887–899.

Whittaker, R. J., K. A. Triantis, and R. J. Ladle. 2008. A general dynamic theory of oceanic island biogeography. Journal of Biogeography 35:977–994.

Wilcox, T. M., M. K. Schwartz, and W. H. Lowe. 2018. Evolutionary community ecology: Time to think outside the (taxonomic) box. Trends in Ecology & Evolution 33:240 – 250.

