## Supplementary material for "The limits to ecological limits to diversification": Figures Tables and Supplementary Data and Scripts: CSvsIW_bootstrap.R

```
CSvsIW_bootstrap <- function(the_dataset,the_island,model)
{
  cluster_folder <- '/home/tece/Rampal/CSvsIW'
  res <- 200
  ## load data
  if (the_dataset == 1)
  {
    the_dataset <- "CSdata"
    load(paste(cluster_folder,'/frogs_sim_CS_T30_M1000_R5000.Rdata',sep=""))
  }
  if (the_dataset == 2)
  {
    the_dataset <- "IWdata"
    load(paste(cluster_folder,'/frogs_sim_IW_T30_M1000_R5000.Rdata',sep=""))
  }
  if (the_dataset == 3)
  {
    the_dataset <- "empdata"
    load(paste(cluster_folder,'/Frogs_100_30_miss_spec_added_M1000.Rdata',sep=""))
    frogs_sims <- frogs
  }
  
  not_present <- c()
  for(i in 1:length(frogs_sims))
  {
     if(frogs_sims[[i]][[1]]$not_present == 1000)
     {
        not_present <- append(not_present, i)
     } 
  }
  if(!is.null(not_present)) frogs_sims <- frogs_sims[-not_present]

  datalist <- frogs_sims[[the_island]]

  island_age <- 30
  M <- 1000
  tol <- c(1E-4, 1E-6, 1E-7)
  optimmethod <- 'simplex'
  cond <- 1
  methode <- 'odeint::runge_kutta_fehlberg78'

  ## CS DD model no ANA
  if(model == '3') {
    ddmodel <- 11
    idparsopt <- c(1,2,3,4)
    idparsnoshift <- 6:10
    parsfix <- 0
    idparsfix <- 5
    clade_sizes <- unlist(lapply(datalist[-1], function(x) length(x$branching_times[-1])))
    miss_specs <- unlist(lapply(datalist[-1], function(x) x$missing_species))
    K <- max(c(3.645000e+01,clade_sizes + miss_specs))
    initparsopt <- c(4.368768e-01, 1.126373e-01, K, 2.187464e-04)
    model_type <- 'CS'
  }

  ## IW model ANA 0
  if(model == '5') {
    ddmodel <- 11
    idparsopt <- c(1,2,3,4)
    parsfix <- 0
    idparsfix <- 5
    K <- max(1.319595e+02,sum(datalist[[1]]$stt_all[nrow(datalist[[1]]$stt_all),c(2,3,4)]))
    initparsopt <- c(4.012298e-01, 1.699521e-01, K, 3.487955e-04)
    verbose <- 1
    model_type <- 'IW'
  }

  outputfilename <- paste(cluster_folder,'/results_',the_dataset,'_',the_island,'_',model,'.txt',sep = '')
  if(file.exists(outputfilename))
  {
    stop('This jobs has been done already')
  }

  cat(paste('model fitted = ',model_type,' \n',sep = ' '))
  cat(paste('fitted to dataset =', the_dataset,' \n',sep = ' '))
  cat(paste('initial parameters = '))
  cat(initparsopt)
  cat('\n')

  ##### RUN1 ML Optimization
  if(model_type == 'CS')
  {
    lik_res <- DAISIE_ML(datalist = datalist,
                         initparsopt = initparsopt,
                         idparsopt = idparsopt,
                         parsfix = parsfix,
                         idparsfix = idparsfix,
                         ddmodel = ddmodel,
                         methode = methode,
                         CS_version = 0,
                         cond = cond,
                         res = res,
                         tol = tol,
                         optimmethod = optimmethod)
  }
  if(model_type == 'IW' )
  {
    lik_res <- DAISIE_ML_IW(datalist = datalist,
                            initparsopt = initparsopt,
                            idparsopt = idparsopt,
                            parsfix = parsfix,
                            idparsfix = idparsfix,
                            ddmodel = ddmodel,
                            methode = methode,
                            cond = cond,
                            res = res,
                            verbose = verbose,
                            tol = tol,
                            optimmethod = optimmethod)
  }

  lik_res <- as.matrix(lik_res)
  to.write <- c(the_island,
                the_dataset,
                island_age,
                M,
                the_island,
                model,
                model_type,
                methode,
                "R1",
                as.matrix(lik_res),
                initparsopt,
                cond)
  write(to.write,file = outputfilename,ncol = length(to.write),append = TRUE,sep = "\t")
}
```
