## Supplementary material for "The limits to ecological limits to diversification": Figures Tables and Supplementary Data and Scripts: CsvsIW_tutorial_FOR_PAPER.html

Fitting clade-specific and island-wide diversity-dependence models in DAISIE


### Fitting clade-specific and island-wide diversity-dependence models in DAISIE

#### 22/04/2020

Load required package DAISIE

```
library(DAISIE)
```

##### 1. Load, prepare and visualize data

- If you want to start directly with model fitting you can skip this step and go to step 2.

Load the phylogenetic data for *Eleutherodacylus* frogs from the island of Hispaniola (used in Etienne et al. 2020). This dataset contains times of colonization and branching times for all species of *Eleutherodactylus* frogs found on the island. The data was extracted from the dated molecular phylogeny of Dugo-Cota et al 2019 (*Ecology Letters* 22:884–893). The dataset includes 65 extant species, which are the result of five independent colonisation events of the island of Hispaniola.

```
data(frogs_datatable, package = "DAISIE")
```

You can load your own data in a table/tibble format, making sure the table headers match the ones in the example.

Eleutherodactylus data table

| Clade\_name | Status | Missing\_species | Branching\_times |
| --- | --- | --- | --- |
| Clade1 | Endemic | 0 | 22.090381975,19.03474267,17.558899475,16.551943175,15.163574647,12.539890494,12.418166302,11.224140324,9.356313022,8.838345923,7.763917546,6.976872428,6.225535797,6.128306647,5.891490086,5.534313767,5.268246402,4.325862315,4.130254551,3.876175449,3.576820015,2.700899509,1.988973569,0.609739668,0.16651871,0.149491685,0.138726728,0.038275342 |
| Clade2 | Endemic | 0 | 13.746625455,8.647464943,8.334079223,7.411910463,6.861782812,6.516839176,5.584497643,4.839004311,4.549481508,4.254369184,4.188654945,3.73901781,3.667503378,3.593658733,3.460220365,3.419754656,3.30938242,2.883634139,2.387378752,1.77824651,0.568917703 |
| Clade3 | Endemic | 0 | 11.032464497,0.224004413,0.102526748 |
| Clade4 | Endemic | 0 | 8.430721468,4.130831021,1.420117489,1.171228154,0.231664348,0.197599408,0.075831995,0.040222432 |
| Clade5 | Endemic | 0 | 8.852578907,4.93343221,3.230048172,1.259294338,0.235900375 |

###### The table contains the following 4 columns (column headers need to be written exactly like this):

- `Clade_name` - Name of the clade/lineage on the island (e.g. clade code, genus name)
- `Status` - Endemicity status of the clade. Can be the following `Endemic`; `Non_endemic`; `Endemic_MaxAge`, `Non_Endemic_MaxAge` The latter two options are for cases when the time of colonisation is believed to be an overestimate. In these cases, DAISIE will assume colonisation happened any time between the colonisation time given in `Branching_times` and the present, or between the colonisation time given in `Branching_times` and the age of the first cladogenesis event in the lineage (if any). The MaxAge options can also be used when the colonization time is unknown, specifying “NA” in the column `Branching_times`, in which case DAISIE will assume that colonisation happened any time between the age of the island and the present.
- `Missing_species` - Number of extant species that belong to the clade but are not included in the phylogeny.
- `Branching_times` - First element is the colonisation time, subsequent elements are branching times within the island if there are any. e.g. c(colonisation\_time, branching\_time1; branching\_time2). If colonisation time is unknown, ‘NA’ can be specified. If branching times are unknown, do not replace them with ‘NA’ - just add the corresponding number of missing species to the column `Missing_species`.

##### 1.1. Create DAISIE datalist from the input table

In this case we are assuming an island age of 30 million years for Hispaniola and a mainland pool size of 300 species (this is the number of species that may colonize the island).

```
frogs_datalist <- DAISIE_dataprep(
        datatable = frogs_datatable,
        island_age = 30,
        M = 300)
```

##### 1.2. View DAISIE datalist

Just type:

```
frogs_datalist
```

The DAISIE datalist includes the data from the table `frogs` in the format of a DAISIE object that can be read in the subsequent maximum likelihood functions (e.g `DAISIE_ML`, `DAISIE_ML_IW` ). It includes the age of the island, the number of mainland species that are not currently present on the island and a list with all extant independent colonisation events, including their colonization and branching times.

##### 1.3. Visualise the data

```
DAISIE_plot_island(frogs_datalist)
```

This plot shows the different colonization events, their times of colonization and branching times (horizontal ticks). Dashed line shows the age of the island. n=number of species; m= number of species missing.

###### Plot age versus diversity

```
DAISIE_plot_age_diversity(frogs_datalist)
```

##### 2. Fit DAISIE models

If you skipped step 1, first load the Hispaniola *Eleutherodactylus* datalist.

```
data(frogs_datalist)
```

\_

We will fit five different DAISIE models to the phylogenetic data contained in `frogs_datalist`:

- **DI** - diversity-independent model. Model where K’ (carrying-capacity) is set to `Inf`.
- **CS** - clade-specific diversity-dependence model.
- **CS\_no\_ana** - clade-specific diversity-dependence model with no anagenesis. *Equivalent to the CS model in Etienne et al. 2020*.
- **IW** - island-wide diversity-dependence model.
- **IW\_no\_ana** - island-wide diversity-dependence model with no anagenesis. *Equivalent to the IW model in Etienne et al. 2020*.

We use the `DAISIE_ML` and `DAISIE_ML_IW` functions to optimise the likelihood. These are the most important settings to specify in these functions:

- `datalist` - The name of the DAISIE datalist (in this case `frogs_datalist`)
- `initparsopt` - these are the initial values from the parameters for which the likelihood will be optimised. In the examples below we will use parameters similar to the maximum likelihood parameters from the corresponding models in Etienne et al 2020. However, we recommend you try a variety of initial starting parameters to ensure the optimum is found. Note that certain combinations of initial starting values may fail, as parameters must be feasible (e.g. a K’ value lower than the number of species found in the data will not run). In DAISIE the parameters have the following position:
  1. Rate of cladogenesis (per species on the island per time unit)
  2. Rate of extinction (per species on the island per time unit)
  3. Carrying-capacity (K’) - maximum number of species any clade can reach on the island for CS models; maximum number of species on the island across all clades for IW models.
  4. Rate of colonisation (per species on the mainland per time unit)
  5. Rate of anagenesis (per species on the island per time unit)
- `idparsopt` - The position of the parameters to optimise (e.g. to optimize cladogenesis and anagenesis only c(1,5).
- `parsfix` - If parameters are being fixed, specify here the value (e.g. if fixing K’ to Inf and fixing anagenesis to 0 - c(Inf,0).
- `idparsfix` - The position of the paratemers fixed (e.g. to fix K’ and anagenesis c(3,5)).
- `ddmodel` - Set 0 for diversity-independent models; 11 for diversity-dependent models (IW and CS) where cladogenesis and colonisation decline linearly with diversity. There are other options.
- There are many other options, check `? DAISIE_ML`

> **The IW models (IW and IW\_no\_ana) use the island-wide version of DAISIE, which is computationally demanding. They require high memory and long run times. We recommend these are run on a cluster. The CS models can be run on a regular computer or laptop.**

##### 2.1 Fit DI model

This model contains 4 parameters:  
1 - cladogenesis  
2 - extinction  
4 - colonisation  
5 - anagenesis  
K’ is fixed to `Inf`  
Set `ddmodel=0`

```
DAISIE_ML(
  datalist = frogs_datalist,
  initparsopt = c(0.18,0.03,0.0006,2),
  idparsopt = c(1,2,4,5),
  ddmodel = 0,
  parsfix = Inf,
  idparsfix = 3
)
#> You are optimizing lambda_c mu gamma lambda_a 
#> You are fixing K 
#> Calculating the likelihood for the initial parameters. 
#> The loglikelihood for the initial parameter values is -209.8779 
#> Optimizing the likelihood - this may take a while. 
#> 
#>  Maximum likelihood parameter estimates: lambda_c: 0.179159, mu: 0.025533, K: Inf,
#>       gamma: 0.000627, lambda_a: 495.568077 
#>  Maximum loglikelihood: -209.863423
```

##### 2.2 Fit CS model

This model contains 5 parameters:  
1 - cladogenesis  
2 - extinction  
3 - K’  
4 - colonisation  
5 - anagenesis  
Set `ddmodel=11`

```
DAISIE_ML(
  datalist = frogs_datalist,
  initparsopt = c(0.44,0.11,36.44,0.0007,2),
  idparsopt = c(1,2,3,4,5),
  ddmodel = 11,
  parsfix = NULL,
  idparsfix = NULL
)
#> You are optimizing lambda_c mu K gamma lambda_a 
#> You are fixing nothing 
#> Calculating the likelihood for the initial parameters. 
#> The loglikelihood for the initial parameter values is -202.6693 
#> Optimizing the likelihood - this may take a while. 
#> 
#>  Maximum likelihood parameter estimates: lambda_c: 0.437117, mu: 0.112778, K: 36.451046,
#>       gamma: 0.000731, lambda_a: 871.024165 
#>  Maximum loglikelihood: -202.653199
```

##### 2.3 Fit CS\_no\_ana model

This model contains 4 parameters:  
1 - cladogenesis  
2 - extinction  
3 - K’  
4 - colonisation  
Set `ddmodel=11`

```
DAISIE_ML(
  datalist = frogs_datalist,
  initparsopt = c(0.44,0.11,36.44,0.0007),
  idparsopt = c(1,2,3,4),
  ddmodel = 11,
  parsfix = 0,
  idparsfix = 5
)
#> You are optimizing lambda_c mu K gamma 
#> You are fixing lambda_a 
#> Calculating the likelihood for the initial parameters. 
#> The loglikelihood for the initial parameter values is -202.6718 
#> Optimizing the likelihood - this may take a while. 
#> 
#>  Maximum likelihood parameter estimates: lambda_c: 0.437021, mu: 0.112473, K: 36.412717,
#>       gamma: 0.000731, lambda_a: 0.000000 
#>  Maximum loglikelihood: -202.656382
```

##### 2.4 Fit IW model

This model contains 5 parameters:  
1 - cladogenesis  
2 - extinction  
3 - K’  
4 - colonisation  
5 - anagenesis  
Set `ddmodel=11`

```
DAISIE_ML_IW(
  datalist = frogs_datalist,
  initparsopt = c(0.41, 0.17, 131.7, 0.0012, 2),
  idparsopt = c(1,2,3,4,5),
  ddmodel = 11,
  parsfix = NULL,
  idparsfix = NULL
)
```

Output not shown here (recommended to be run on a cluster).

##### 2.5 Fit IW\_no\_ana model

This model contains 4 parameters:  
1 - cladogenesis  
2 - extinction  
3 - K’  
4 - colonisation  
Set `ddmodel=11`

```
DAISIE_ML_IW(
  datalist = frogs_datalist,
  initparsopt = c(0.40, 0.17, 131.83, 0.0012),
  idparsopt = c(1,2,3,4),
  ddmodel = 11,
  parsfix = 0,
  idparsfix = 5
)
```

Output not shown here (recommended to be run on a cluster).

##### 3. Simulate islands under given DAISIE models

###### 3.1 Simulate islands with the parameters estimated from the best model for the Hispaniolan *Eleutherodacytlus* data

We use the `DAISIE_sim` function, which simulates diversity dynamics on an island from island birth until a specificied island age, based on a given set of parameters (cladogenesis, extinction, carrying-capacity (K’), colonisation, anagenesis). These are the most important settings to specify in `DAISIE_sim` function:

- `pars` - The values of the 5 parameters in the following order
  1. Rate of cladogenesis
  2. Rate of extinction
  3. Carrying-capacity (K’)
  4. Rate of colonisation
  5. Rate of anagenesis
- `replicates` - Number of replicates to simulate
- `time` - Time to run the simulation, for example 20 million years
- `M` - Number species in the mainland pool
- `divdepmodel` - set `CS` for clade-specific diversity-dependence; or `IW` for island-wide diversity-dependence

Simulate a CS model for 30 million years, 1000 replicates:

```
frog_sims_CS<-DAISIE_sim_constant_rate(
  time=30,
  M=300,
  pars=c(0.44,0.11,36.44,0.0007,0),
  divdepmodel = "CS",
  replicates= 1000,
  plot_sims = FALSE)
```

Simulate an IW model for 30 million years, 1000 replicates:

```
frog_sims_CS<-DAISIE_sim_constant_rate(
  time=30,
  M=300,
  pars=c(0.40,0.17,131.83,0.0012,0),
  divdepmodel = "IW",
  replicates= 1000,
  plot_sims = FALSE)
```

###### 3.2 Plot the species-through-time plots resulting from the simulations.

```
DAISIE_plot_sims(frog_sims_CS)
```
