## Supplementary material for "The limits to ecological limits to diversification": Figures Tables and Supplementary Data and Scripts: CsvsIW_tutorial_FOR_PAPER.pdf

### Fitting clade-specific and island-wide diversity-dependence models in DAISIE

22/04/2020

Load required package DAISIE

```r
library(DAISIE)
```

##### 1. Load, prepare and visualize data

- If you want to start directly with model fitting you can skip this step and go to step 2.

Load the phylogenetic data for *Eleutherodacylus* frogs from the island of Hispaniola (used in Etienne et al. 2020). This dataset contains times of colonization and branching times for all species of *Eleutherodactylus* frogs found on the island. The data was extracted from the dated molecular phylogeny of Dugo-Cota et al 2019 (*Ecology Letters* 22:884–893). The dataset includes 65 extant species, which are the result of five independent colonisation events of the island of Hispaniola.

```r
data(frogs_datatable, package = "DAISIE")
```

You can load your own data in a table/tibble format, making sure the table headers match the ones in the example.

| Clade_name | Status | Missing_species | Branching_times |
|---|---|---|---|
| Clade1 | Endemic | 0 | 22.090381975,19.03474267,17.558899475,16.551943175,15.163574647,12.539890[illegible] |
| Clade2 | Endemic | 0 | 13.746625455,8.647464943,8.334079223,7.411910463,6.861782812,6.516839176,5[illegible] |
| Clade3 | Endemic | 0 | 11.032464497,0.224004413,0.102526748 |
| Clade4 | Endemic | 0 | 8.430721468,4.130831021,1.420117489,1.171228154,0.231664348,0.197599408,0.[illegible] |
| Clade5 | Endemic | 0 | 8.852578907,4.93343221,3.230048172,1.259294338,0.235900375 |

##### 1.3. Visualise the data

```
DAISIE_plot_island(frogs_datalist)
```

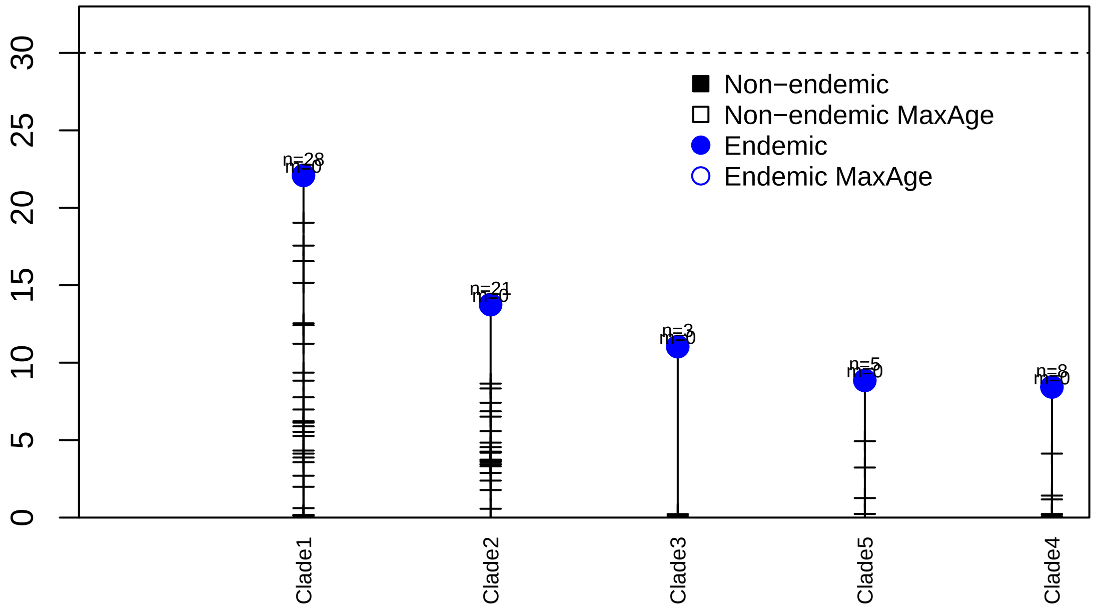


This plot shows the different colonization events, their times of colonization and branching times (horizontal ticks). Dashed line shows the age of the island. n=number of species; m= number of species missing.

##### Plot age versus diversity

```
DAISIE_plot_age_diversity(frogs_datalist)
```

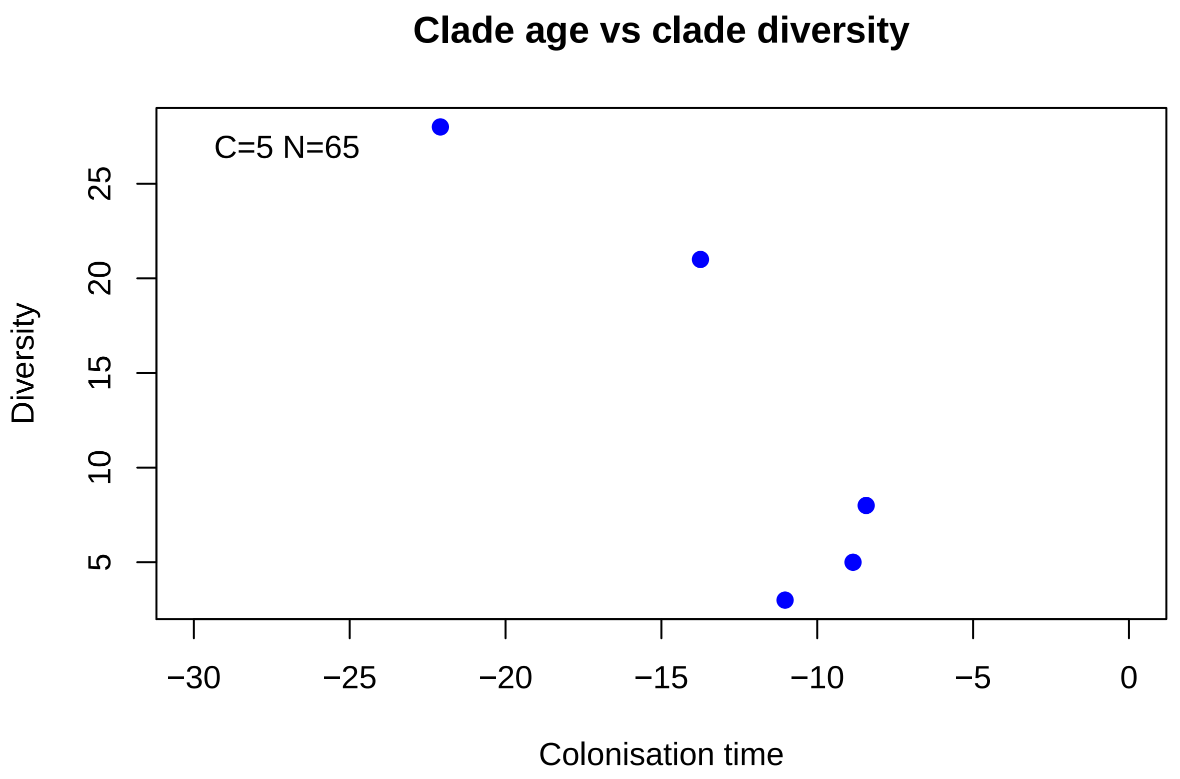


**2. Fit DAISIE models**

If you skipped step 1, first load the Hispaniola *Eleutherodactylus* datalist.

```
data(frogs_datalist)
```

—

We will fit five different DAISIE models to the phylogenetic data contained in `frogs_datalist`:

```
#>
#>  Maximum likelihood parameter estimates: lambda_c: 0.179159, mu: 0.025533, K: Inf,
#>       gamma: 0.000627, lambda_a: 495.568077
#>  Maximum loglikelihood: -209.863423
#>    lambda_c         mu   K        gamma lambda_a    loglik df conv
#> 1 0.1791595 0.02553254 Inf 0.0006268835 495.5681 -209.8634  4    0
```

##### 2.2 Fit CS model

This model contains 5 parameters:
1 - cladogenesis
2 - extinction
3 - K'
4 - colonisation
5 - anagenesis
Set `ddmodel=11`

```
DAISIE_ML(
  datalist = frogs_datalist,
  initparsopt = c(0.44,0.11,36.44,0.0007,2),
  idparsopt = c(1,2,3,4,5),
  ddmodel = 11,
  parsfix = NULL,
  idparsfix = NULL
)
#> You are optimizing lambda_c mu K gamma lambda_a
#> You are fixing nothing
#> Calculating the likelihood for the initial parameters.
#> The loglikelihood for the initial parameter values is -202.6693
#> Optimizing the likelihood - this may take a while.
#>
#>  Maximum likelihood parameter estimates: lambda_c: 0.437117, mu: 0.112778, K: 36.451046,
#>       gamma: 0.000731, lambda_a: 871.024165
#>  Maximum loglikelihood: -202.653199
#>   lambda_c        mu        K        gamma lambda_a    loglik df conv
#> 1 0.437117 0.1127778 36.45105 0.0007314647 871.0242 -202.6532  5    0
```

##### 2.3 Fit CS_no_ana model

This model contains 4 parameters:
1 - cladogenesis
2 - extinction
3 - K'
4 - colonisation
Set `ddmodel=11`

```r
DAISIE_ML(
  datalist = frogs_datalist,
  initparsopt = c(0.44,0.11,36.44,0.0007),
  idparsopt = c(1,2,3,4),
  ddmodel = 11,
  parsfix = 0,
  idparsfix = 5
)
#> You are optimizing lambda_c mu K gamma
#> You are fixing lambda_a
#> Calculating the likelihood for the initial parameters.
#> The loglikelihood for the initial parameter values is -202.6718
#> Optimizing the likelihood - this may take a while.
#>
#>  Maximum likelihood parameter estimates: lambda_c: 0.437021, mu: 0.112473, K: 36.412717,
#>       gamma: 0.000731, lambda_a: 0.000000
#>  Maximum loglikelihood: -202.656382
#>    lambda_c        mu        K        gamma lambda_a    loglik df conv
#> 1 0.4370211 0.1124732 36.41272 0.0007305765        0 -202.6564  4    0
```

##### 2.4 Fit IW model

This model contains 5 parameters:
1 - cladogenesis
2 - extinction
3 - K'
4 - colonisation
5 - anagenesis
Set `ddmodel=11`

```r
DAISIE_ML_IW(
  datalist = frogs_datalist,
  initparsopt = c(0.41, 0.17, 131.7, 0.0012, 2),
  idparsopt = c(1,2,3,4,5),
  ddmodel = 11,
  parsfix = NULL,
  idparsfix = NULL
)
```

Output not shown here (recommended to be run on a cluster).

##### 2.5 Fit IW_no_ana model

This model contains 4 parameters:
1 - cladogenesis
2 - extinction
3 - K'
4 - colonisation
Set `ddmodel=11`

```r
DAISIE_ML_IW(
  datalist = frogs_datalist,
  initparsopt = c(0.40, 0.17, 131.83, 0.0012),
  idparsopt = c(1,2,3,4),
  ddmodel = 11,
  parsfix = 0,
  idparsfix = 5
)
```

Output not shown here (recommended to be run on a cluster).

##### 3. Simulate islands under given DAISIE models

###### 3.1 Simulate islands with the parameters estimated from the best model for the Hispaniolan *Eleutherodacytlus* data

We use the `DAISIE_sim` function, which simulates diversity dynamics on an island from island birth until a specificied island age, based on a given set of parameters (cladogenesis, extinction, carrying-capacity (K'), colonisation, anagenesis). These are the most important settings to specify in `DAISIE_sim` function:

- `pars` - The values of the 5 parameters in the following order
  1. Rate of cladogenesis
  2. Rate of extinction
  3. Carrying-capacity (K')
  4. Rate of colonisation
  5. Rate of anagenesis
- `replicates` - Number of replicates to simulate
- `time` - Time to run the simulation, for example 20 million years
- `M` - Number species in the mainland pool
- `divdepmodel` - set `CS` for clade-specific diversity-dependence; or `IW` for island-wide diversity-dependence

Simulate a CS model for 30 million years, 1000 replicates:

```r
frog_sims_CS<-DAISIE_sim_constant_rate(
  time=30,
  M=300,
  pars=c(0.44,0.11,36.44,0.0007,0),
  divdepmodel = "CS",
  replicates= 1000,
  plot_sims = FALSE)
```

Simulate an IW model for 30 million years, 1000 replicates:

```r
frog_sims_CS<-DAISIE_sim_constant_rate(
  time=30,
  M=300,
  pars=c(0.40,0.17,131.83,0.0012,0),
  divdepmodel = "IW",
  replicates= 1000,
  plot_sims = FALSE)
```

**3.2 Plot the species-through-time plots resulting from the simulations.**

```
DAISIE_plot_sims(frog_sims_CS)
```

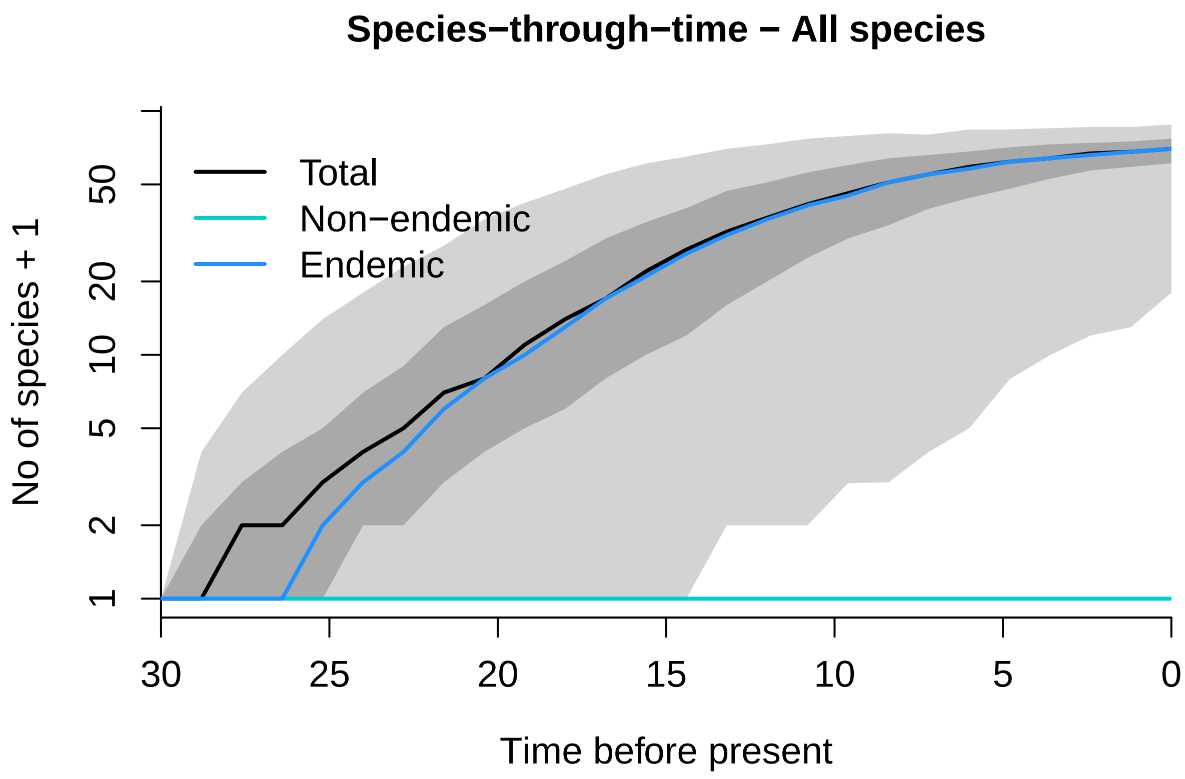
