## Supplementary material for "The limits to ecological limits to diversification": Figures Tables and Supplementary Data and Scripts: data_52_sum.tex

% latex table generated in R 4.2.0 by xtable 1.8-4 package
% Fri Nov 12 15:23:57 2021
\begin{table}[ht]
\centering
\caption{Maximum likelihood parameter estimates and information criteria of the DAISIE
models fitted to the empirical dataset. df is the number of free parameters. The asterisks indicate that the values are very close to 0, but not fixed at 0.\label{ML_estimates}}
\bigskip
\begin{tabular}{lrrrrlrrrrrr}
  \hline
model & $\lambda_0^c$ & $\mu$ & $K$ & $\gamma$ & $\lambda^a$ & LL & df & AIC & AIC weight & BIC & BIC weight \\ 
  \hline
DI & 0.18 & 0.03 & $\infty$ & 0.0002 & 0* & -215.87 & 4 & 439.75 & 0.00 & 445.53 & 0.00 \\ 
DD-CS & 0.44 & 0.11 & 36.44 & 0.0002 & 0* & -208.67 & 5 & 427.34 & 0.27 & 434.58 & 0.15 \\ 
DD-CS-noA & 0.44 & 0.11 & 36.45 & 0.0002 & 0 & -208.67 & 4 & 425.34 & 0.73 & 431.13 & 0.85 \\ 
IW-CS & 0.40 & 0.17 & 131.89 & 0.0003 & 0* & -215.10 & 5 & 440.20 & 0.00 & 447.43 & 0.00 \\ 
IW-CS-noA & 0.40 & 0.17 & 131.96 & 0.0003 & 0 & -215.10 & 4 & 438.20 & 0.00 & 443.98 & 0.00 \\ 
  \hline
\end{tabular}
\end{table}
