## Supplementary material for "The limits to ecological limits to diversification": Figures Tables and Supplementary Data and Scripts: likelihood_computation.tex

%%%%%%%%%%%%%%%%%%%%%%%%%%%%%%%%%%%%%%%%%%%%%%%%%%%%%%%%%%%%%%%%%%%%%%%%%%%%%%%
\section*{Supplementary Text -- Derivation of likelihood computation}
%%%%%%%%%%%%%%%%%%%%%%%%%%%%%%%%%%%%%%%%%%%%%%%%%%%%%%%%%%%%%%%%%%%%%%%%%%%%%%%

We derive the likelihood of colonization and branching data using the $Q$-approach \citep{Etienne-etal:2012, Laudanno-etal:2019}. This approach is based on the quantity $Q(t)$, defined as the probability that a model realization is consistent with the data from the initial time $t_\text{i}$ to a given time $t$.  We construct the differential equations governing the dynamics of $Q(t)$, and explain how the likelihood can be extracted from $Q(t_\text{p})$ at the present time $t_\text{p}$.

A dataset consists of a number of trees, each associated with a mainland species.  An example is shown in Fig.~\ref{figcomp1}.  Several trees can be associated with the same mainland species, if it has colonized the island several times (e.g., species~B in the figure).  For trees consisting of a single branch, the dataset also specifies whether speciation has taken place, i.e., whether the species at the present time is endemic to the island (compare species~C and D).  

To set up the likelihood computation, we partition the number of extant species on the island at any time $t$ as follows: \\[-18pt]
\begin{center}
\begin{tabular}{cp{.9\textwidth}}
 $k$ & the number of species present on the island at time $t$ that are represented in the dataset, i.e., that survive until the present and are sampled; \\[2pt]
 $m$ & the number of mainland species present on the island at time $t$ that are not in the data (either because the lineage became extinct before the present or because it has not been sampled); \\[2pt]
 $e$ & the number of endemic species at time $t$ that are not in the data \\[-6pt]
\end{tabular}
\end{center}
Hence, the total number of extant species at time $t$ is equal to $k+m+e$.  We have $m \leq M$, where $M$ is the number of species in the mainland source pool.  Note that $k$ is determined by the data, while $m$ and $e$ are not.  Instead they vary between model realizations, and are part of the system state in the $Q$-approach.

Fig.~\ref{figcomp2} shows one possible model realization consistent with the dataset of Fig.~\ref{figcomp1}, together with the values that $k$, $m$ and $e$ take through time.

%%%%%%%%%%%%%%%%%%%%%%%%%%%%%%%%%%%%%%%%%%%%%%%%%%%%%%%%%%%%%%%%%%%%%%%%%%%%%%%
\subsection*{Before first colonization recorded in data}
%%%%%%%%%%%%%%%%%%%%%%%%%%%%%%%%%%%%%%%%%%%%%%%%%%%%%%%%%%%%%%%%%%%%%%%%%%%%%%%

Before any colonization event of still extant lineages (so $k = 0$) we keep track of the number of non-endemic (i.e., mainland) species $m$ and the number of endemic species $e$.  The dynamics of the probabilities $Q_{m,e}(t)$, where $m$ and $e$ can take any integer value, are described by the following set of differential equations:
\begin{align}
 \frac{\diff Q_{m,e}}{\diff t} =\;
 & +\gamma_{m+e-1}\big(M-(m-1)\big)Q_{m-1,e}
   && \text{($+$ colonization)} \notag \\[-5pt]
 & +\mu_{m+e+1}(m+1)Q_{m+1,e}
   && \text{($+$ extinction of non-endemic species)} \notag \\
 & +\mu_{m+e+1}(e+1)Q_{m,e+1}
   && \text{($+$ extinction of endemic species)} \notag \\
 & +\lambda_{m+e}^{\text{a}}\left( m+1\right) Q_{m+1,e-1}
   && \text{($+$ anagenesis)} \notag \\
 & +\lambda_{m+e-1}^{\text{c}}(m+1)Q_{m+1,e-2}
   && \text{($+$ cladogenesis of non-endemic species)} \notag \\
 & +\lambda_{m+e-1}^{\text{c}}(e-1)Q_{m,e-1}
   && \text{($+$ cladogenesis of endemic species)} \notag \\
 & -\gamma_{m+e}(M-m) Q_{m,e}
   && \text{($-$ colonization)} \notag \\
 & -\mu_{m+e}(m+e) Q_{m,e}
   && \text{($-$ extinction)} \notag \\
 & -\lambda_{m+e}^{\text{a}}m Q_{m,e}
   && \text{($-$ anagenesis)} \notag \\
 & -\lambda_{m+e}^{\text{c}}(m+e) Q_{m,e}
   && \text{($-$ cladogenesis)}
 \label{eq:dynbefcol}
\end{align}
The positive terms on the right-hand side correspond to events that bring the system in the state with $m$ non-endemic and $e$ endemic species, and hence increase the probability $Q_{m,e}$.  These events are due to colonization, extinction of a non-endemic species, extinction of an endemic species, anagenesis (of a non-endemic species) and cladogenesis in a non-endemic and endemic species.  The negative terms correspond to events that decrease the probability due to the same processes when in the state with $m$ non-endemic and $e$ endemic species, i.e., events that move the system away from this state.

The initial condition for $Q_{m,e}(t)$ at time $t_\text{i}$ usually corresponds to an empty island, so
\begin{equation}
 Q_{m,e}(t_\text{i}) = \begin{cases}
 1 & \text{for } m=e=0 \\
 0 & \text{otherwise} \end{cases}
\end{equation}

Note that because $k = 0$, there are no lineages yet with which the realizations have to be  consistent, and hence equations~(\ref{eq:dynbefcol}) are identical to the master equations for the probabilities $P_{m,e}(t)$ to have $m$ non-endemic and $e$ endemic species.

%%%%%%%%%%%%%%%%%%%%%%%%%%%%%%%%%%%%%%%%%%%%%%%%%%%%%%%%%%%%%%%%%%%%%%%%%%%%%%%
\subsection*{After first colonization recorded in data}
%%%%%%%%%%%%%%%%%%%%%%%%%%%%%%%%%%%%%%%%%%%%%%%%%%%%%%%%%%%%%%%%%%%%%%%%%%%%%%%

At the first colonization event recorded in the data, which we call $t_1$, one of the extant species is represented in the phylogeny (so $k = 1$).  From this time onwards, $Q_{m,e}(t)$ becomes different from $P_{m,e}(t)$, because the realizations contributing to $Q_{m,e}(t)$ not only have $m$ non-endemic species and $e$ endemic species at time $t$, but should also be consistent with the colonization at time $t_1$.

After the first colonization event, recolonization of the same mainland species is not allowed until the colonizing species undergoes speciation, because it would change the data, namely the time of colonization.  This implies that we have to keep track of whether the colonizing species has undergone speciation.  We therefore expand the set of equations~(\ref{eq:dynbefcol}) to two sets. The first set describes the probabilities where speciation has not (yet) taken place. The probabilities in this set are denoted by $Q_{m,e}^{0}(t)$, where the superscript 0 indicates that speciation has not (yet) taken place. The second set describes the probabilities where speciation has taken place and recolonization of the same species can therefore occur without changing the data. The probabilities in this set are denoted by $Q_{m,e}^{1}(t)$, where the superscript 1 indicates that speciation has taken place.

The dynamical equations for $Q_{m,e}^0(t)$ are
\begin{align}
 \frac{\diff Q_{m,e}^{0}}{\diff t} =\;
 & +\gamma_{m+e}\big((M-1)-(m-1)\big)Q_{m-1,e}^{0} \notag \\[-3pt]
 & +\mu_{m+e+2}(m+1)Q_{m+1,e}^{0} \notag \\
 & +\mu_{m+e+2}(e+1)Q_{m,e+1}^{0} \notag \\
 & +\lambda_{m+e+1}^{\text{a}}\left( m+1\right) Q_{m+1,e-1}^{0} \notag \\
 & +\lambda_{m+e}^{\text{c}}(m+1)Q_{m+1,e-2}^{0} \notag \\
 & +\lambda_{m+e}^{\text{c}}(e-1)Q_{m,e-1}^{0} \notag \\
 & -\gamma_{m+e+1}(M-m) Q_{m,e}^{0} \notag \\
 & -\mu_{m+e+1}(m+e+1) Q_{m,e}^{0} \notag \\
 & -\lambda_{m+e+1}^{\text{a}}\left( m+1\right) Q_{m,e}^{0} \notag \\
 & -\lambda_{m+e+1}^{\text{c}}(m+e+1) Q_{m,e}^{0}
 \label{eq:dynaft1col1}
\end{align}
Importantly, these equations guarantee that the realizations contributing to $Q_{m,e}^0(t)$ are consistent with a colonization event at time $t_1$ in the data.  In particular, the mainland species that has colonized does not recolonize (first line, factor $(M-1)-(m-1)$ instead of $M-(m-1)$) and the colonizing species does not become extinct (second line, factor $m+1$ instead of $m+2$).  If the colonizing species undergoes anagenesis or cladogenesis, $Q_{m,e}^0(t)$ does not increase (fourth and fifth line, factor $m+1$), but $Q_{m,e}^1(t)$ increases (see below).

The dynamical equations for $Q_{m,e}^1(t)$ are
\begin{align}
 \frac{\diff Q_{m,e}^{1}}{\diff t} =\;
 & \lambda_{m+e+1}^{\text{a}}Q_{m,e}^{0}
   +2\lambda_{m+e}^{\text{c}}Q_{m,e-1}^{0} \notag \\
 & +\gamma_{m+e}\big(M-(m-1)\big)Q_{m-1,e}^{1} \notag \\
 & +\mu_{m+e+2}(m+1)Q_{m+1,e}^{1} \notag \\
 & +\mu_{m+e+2}(e+1)Q_{m,e+1}^{1} \notag \\
 & +\lambda_{m+e+1}^{\text{a}}\left(m+1\right) Q_{m+1,e-1}^{1} \notag \\
 & +\lambda_{m+e}^{\text{c}}(m+1)Q_{m+1,e-2}^{1} \notag \\
 & +\lambda_{m+e}^{\text{c}}(e+1)Q_{m,e-1}^{1} \notag \\
 & -\gamma_{m+e+1}(M-m) Q_{m,e}^{1} \notag \\
 & -\mu_{m+e+1}(m+e+1) Q_{m,e}^{1} \notag \\
 & -\lambda_{m+e+1}^{\text{a}}m Q_{m,e}^{1} \notag \\
 & -\lambda_{m+e+1}^{\text{c}}(m+e+1) Q_{m,e}^{1}
 \label{eq:dynaft1col2}
\end{align}
The first line on the right-hand side corresponds to a speciation of the colonizing species, either anagenesis or cladogenesis.  Observe the factor 2 in the cladogenesis term.  This is a combinatorial factor required in the $Q$-approach, which can be understood intuitively by noting that either of the two daughter species can be represented in the phylogeny (see \citet{Laudanno-etal:2019} for a detailed justification).

Equations~(\ref{eq:dynaft1col2}) guarantee that the lineage that originates at colonization time $t_1$ does not become extinct (fourth line, factor $e+1$ instead of $e+2$).  In contrast to equations~(\ref{eq:dynaft1col1}), the mainland species that colonized is allowed to recolonize (because it does not change the data; second line, factor $M-(m-1)$).  The term corresponding to cladogenesis of endemic species (seventh line) has a factor $e+1$, because there are $e$ endemic species in total and the species represented in the phylogeny (i.e., the species that descends from the colonization event at time $t_1$) should be counted twice, due to the combinatorial factor 2 we mentioned above.

The initial condition for equations (\ref{eq:dynaft1col1}--\ref{eq:dynaft1col2}) at time $t_1$ impose the colonization of the appropriate mainland species,
\begin{align}
 Q_{m,e}^{0}(t_1) &= \gamma_{m+e}\,\diff t\; Q_{m,e}(t_1) \notag \\
 Q_{m,e}^{1}(t_1) &= 0
 \label{eq:init1col}
\end{align}
where the infinitesimal $\diff t$ makes clear that we are looking at the probability of colonization at precisely time $t_1$, i.e., between $t_1$ and $t_1 + \diff t$.  Hence, the conditions~(\ref{eq:init1col}) connect equations~(\ref{eq:dynbefcol}) before and equations~(\ref{eq:dynaft1col1}--\ref{eq:dynaft1col2}) after the colonization event.

%%%%%%%%%%%%%%%%%%%%%%%%%%%%%%%%%%%%%%%%%%%%%%%%%%%%%%%%%%%%%%%%%%%%%%%%%%%%%%%
\subsection*{After second colonization recorded in data}
%%%%%%%%%%%%%%%%%%%%%%%%%%%%%%%%%%%%%%%%%%%%%%%%%%%%%%%%%%%%%%%%%%%%%%%%%%%%%%%

Suppose the next event in the data is a new colonization of another mainland species, which occurs at time $t_2$.  Again, we have to keep track of whether this colonizing species has undergone speciation.  We proceed as before, and expand the set of equations (\ref{eq:dynaft1col1}--\ref{eq:dynaft1col2}) to two sets: a first set for the probabilities where speciation has not (yet) taken place, and a second set for the probabilities where speciation has taken place.  This leads to four sets of dynamical variables, $Q_{m,e}^{0,0}$, $Q_{m,e}^{0,1}$, $Q_{m,e}^{1,0}$ and $Q_{m,e}^{1,1}$, where the first and second bit indicate the speciation status of the first and second colonization, respectively.

The corresponding dynamical equations are
\begin{align}
 \frac{\diff Q_{m,e}^{0,0}}{\diff t} =\;
 & +\gamma_{m+e+1}\big((M-2)-(m-1)\big)Q_{m-1,e}^{0,0} \notag \\[-3pt]
 & +\mu_{m+e+3}(m+1)Q_{m+1,e}^{0,0} \notag \\
 & +\mu_{m+e+3}(e+1)Q_{m,e+1}^{0,0} \notag \\
 & +\lambda_{m+e+2}^{\text{a}}(m+1)Q_{m+1,e-1}^{0,0} \notag \\
 & +\lambda_{m+e+1}^{\text{c}}(m+1)Q_{m+1,e-2}^{0,0} \notag \\
 & +\lambda_{m+e+1}^{\text{c}}(e-1)Q_{m,e-1}^{0,0} \notag \\
 & -\gamma_{m+e+2}(M-m)Q_{m,e}^{0,0} \notag \\
 & -\mu_{m+e+2}(m+e+2)Q_{m,e}^{0,0} \notag \\
 & -\lambda_{m+e+2}^{\text{a}}(m+2)Q_{m,e}^{0,0} \notag \\
 & -\lambda_{m+e+2}^{\text{c}}(m+e+2)Q_{m,e}^{0,0}
 \label{eq:dynaft2col1}
\end{align}
where we exclude the recolonization of the two colonizing mainland species (first line, factor $(M-2)-(m-1)$ instead of $M-(m-1)$);
\begin{align}
 \frac{\diff Q_{m,e}^{0,1}}{\diff t} =\;
 & \lambda_{m+e+2}^{\text{a}}Q_{m,e}^{0,0}
   +2\lambda_{m+e+1}^{\text{c}}Q_{m,e-1}^{0,0} \notag \\[-3pt]
 & +\gamma_{m+e+1}\big((M-1)-(m-1)\big)Q_{m-1,e}^{0,1} \notag \\
 & +\mu_{m+e+3}(m+1)Q_{m+1,e}^{0,1} \notag \\
 & +\mu_{m+e+3}(e+1)Q_{m,e+1}^{0,1} \notag \\
 & +\lambda_{m+e+2}^{\text{a}}\left(m+1\right) Q_{m+1,e-1}^{0,1} \notag \\
 & +\lambda_{m+e+1}^{\text{c}}(m+1)Q_{m+1,e-2}^{0,1} \notag \\
 & +\lambda_{m+e+1}^{\text{c}}(e+1)Q_{m,e-1}^{0,1} \notag \\
 & -\gamma_{m+e+2}(M-m)Q_{m,e}^{0,1} \notag \\
 & -\mu_{m+e+2}(m+e+2)Q_{m,e}^{0,1} \notag \\
 & -\lambda_{m+e+2}^{\text{a}}(m+1)Q_{m,e}^{0,1} \notag \\
 & -\lambda_{m+e+2}^{\text{c}}(m+e+2)Q_{m,e}^{0,1}
 \label{eq:dynaft2col2}
\end{align}
where we exclude the recolonization of the first colonizing mainland species;
\begin{align}
 \frac{\diff Q_{m,e}^{1,0}}{\diff t} =\;
 & \lambda_{m+e+2}^{\text{a}}Q_{m,e}^{0,0}
   +2\lambda_{m+e+1}^{\text{c}}Q_{m,e-1}^{0,0} \notag \\[-3pt]
 & +\gamma_{m+e+1}\big((M-1)-(m-1)\big)Q_{m-1,e}^{1,0} \notag \\
 & +\mu_{m+e+3}(m+1)Q_{m+1,e}^{1,0} \notag \\
 & +\mu_{m+e+3}(e+1)Q_{m,e+1}^{1,0} \notag \\
 & +\lambda_{m+e+2}^{\text{a}}\left(m+1\right) Q_{m+1,e-1}^{1,0} \notag \\
 & +\lambda_{m+e+1}^{\text{c}}(m+1)Q_{m+1,e-2}^{0,0} \notag \\
 & +\lambda_{m+e+1}^{\text{c}}(e+1)Q_{m,e-1}^{1,0} \notag \\
 & -\gamma_{m+e+2}(M-m)Q_{m,e}^{1,0} \notag \\
 & -\mu_{m+e+2}(m+e+2)Q_{m,e}^{1,0} \notag \\
 & -\lambda_{m+e+2}^{\text{a}}(m+1)Q_{m,e}^{1,0} \notag \\
 & -\lambda_{m+e+2}^{\text{c}}(m+e+2)Q_{m,e}^{1,0}
 \label{eq:dynaft2col3}
\end{align}
where we exclude the recolonization of the second colonizing mainland species;
\begin{align}
\frac{\diff Q_{m,e}^{1,1}}{\diff t} =\;
 & \lambda_{m+e+2}^{\text{a}}Q_{m,e}^{0,1}
   +2\lambda_{m+e+1}^{\text{c}}Q_{m,e-1}^{0,1} \notag \\[-3pt]
 & +\lambda_{m+e+2}^{\text{a}}Q_{m,e}^{1,0}
   +2\lambda_{m+e+1}^{\text{c}}Q_{m,e-1}^{1,0} \notag \\
 & +\gamma_{m+e+1}\big(M-(m-1)\big)Q_{m-1,e}^{1,1} \notag \\
 & +\mu_{m+e+3}(m+1)Q_{m+1,e}^{1,1} \notag \\
 & +\mu_{m+e+3}(e+1)Q_{m,e+1}^{1,1} \notag \\
 & +\lambda_{m+e+2}^{\text{a}}\left(m+1\right) Q_{m+1,e-1}^{1,1} \notag \\
 & +\lambda_{m+e+1}^{\text{c}}(m+1)Q_{m+1,e-2}^{1,1} \notag \\
 & +\lambda_{m+e+1}^{\text{c}}(e+3)Q_{m,e-1}^{1,1} \notag \\
 & -\gamma_{m+e+2}(M-m)Q_{m,e}^{1,1} \notag \\
 & -\mu_{m+e+2}(m+e+2)Q_{m,e}^{1,1} \notag \\
 & -\lambda_{m+e+2}^{\text{a}}mQ_{m,e}^{1,1} \notag \\
 & -\lambda_{m+e+2}^{\text{c}}(m+e+2)Q_{m,e}^{1,1}
 \label{eq:dynaft2col4}
\end{align}
Note the factor $e+3$ in the term corresponding to cladogenesis of endemic species (eighth line): there are $e+1$ endemic species and the two species represented in the phylogeny (the species descending from the two colonization events) should be counted twice.

The initial condition at colonization time $t_2$ connects the solutions before (equations (\ref{eq:dynaft1col1}--\ref{eq:dynaft1col2})) and after (equations (\ref{eq:dynaft2col1}--\ref{eq:dynaft2col4})) the colonization event,
\begin{align}
 Q_{m,e}^{0,0}(t_2)
 &= \gamma_{m+e+1}\,\diff t\; Q_{m,e}^{0}(t_2) \notag \\
 Q_{m,e}^{0,1}(t_2) &= 0 \notag \\
 Q_{m,e}^{1,0}(t_2)
 &=\gamma_{m+e+1}\,\diff t\; Q_{m,e}^{1}(t_2) \notag \\
 Q_{m,e}^{1,1}(t_2) &=0
 \label{eq:init2col}
\end{align}

%%%%%%%%%%%%%%%%%%%%%%%%%%%%%%%%%%%%%%%%%%%%%%%%%%%%%%%%%%%%%%%%%%%%%%%%%%%%%%%
\subsection*{After $\ell$-th colonization recorded in data}
%%%%%%%%%%%%%%%%%%%%%%%%%%%%%%%%%%%%%%%%%%%%%%%%%%%%%%%%%%%%%%%%%%%%%%%%%%%%%%%

Now we generalize equations (\ref{eq:dynaft1col1}--\ref{eq:dynaft1col2}) and (\ref{eq:dynaft2col1}--\ref{eq:dynaft2col4}) to an arbitary number of colonization events recorded in the data.  We define the number $\ell$ as the number of colonizations from which originate the species at time $t$ represented in the phylogeny (see Fig.~\ref{figcomp2} for an illustration).  Recall that there are $k$ such species, so we have $\ell \leq k$.

For each of the $\ell$ colonizations we have to keep track of whether speciation has taken place in the corresponding lineage (see previous sections for the cases $\ell=1$ and $\ell=2$).  Therefore, we introduce a binary vector $\vec\sigma$ of length $\ell$, the $i$-th component of which indicates whether speciation has taken place ($\sigma_i=1$) or not ($\sigma_i=0$) in the lineage that originates from the $i$-th colonization event.  We add this vector to the system state, so that it consists of the number of non-endemic species $m$, the number of endemic species $e$ and the speciation state $\vec\sigma$ of the $\ell$ colonization events.  We denote the corresponding dynamical variable by $Q_{m,e}^{\vec\sigma}(t)$.

We need some additional notation to formulate the dynamical equations.  We denote by $\ell^0$ the number of colonizations recorded in the data for which speciation has not (yet) taken place, equal to the number of zeros in $\vec\sigma$ (see Fig.~\ref{figcomp2} for an illustration).

For a given vector $\vec\sigma$, we define $\sgma{i}{0}$ as the vector with the same components except $\sigma_i=0$, and $\sgma{i}{1}$ as the vector with the same components except $\sigma_i=1$.  The vector $\vec\sigma^{\times}$ stands for the vector with the last component removed (i.e., a vector of length $\ell-1$).  For example, for $\vec\sigma = (1,0,1,0,0)$, we have $\sgma{\scaleto{2}{4pt}}{1} = (1,1,1,0,0)$, $\sgma{\scaleto{3}{4pt}}{0} = (1,0,0,0,0)$ and $\vec\sigma^{\times} = (1,0,1,0)$.

Then, the dynamical equations are
\begin{align}
 \frac{\diff Q_{m,e}^{\vec{\sigma}}}{\diff t} =\;
 & \sum_{i \mid \sigma_i=1}
   \Big( \lambda_{k+m+e}^{\text{a}}Q_{m,e}^{\sgmaa{i}{0}}
       +2\lambda_{k+m+e-1}^{\text{c}}Q_{m,e-1}^{\sgmaa{i}{0}} \Big) \notag \\[-3pt]
 & +\gamma_{k+m+e-1}\big(M-\ell^0-(m-1)\big)Q_{m-1,e}^{\vec\sigma} \notag \\
 & +\mu_{k+m+e+1}(m+1)Q_{m+1,e}^{\vec\sigma} \notag \\
 & +\mu_{k+m+e+1}(e+1)Q_{m,e+1}^{\vec\sigma} \notag \\
 & +\lambda_{k+m+e}^{\text{a}}(m+1)Q_{m+1,e-1}^{\vec\sigma} \notag \\
 & +\lambda_{k+m+e-1}^{\text{c}}(m+1)Q_{m+1,e-2}^{\vec\sigma} \notag \\
 & +\lambda_{k+m+e-1}^{\text{c}}\big(2(k-\ell^0)+(e-1)\big)Q_{m,e-1}^{\vec\sigma} \notag \\
 & -\gamma_{k+m+e}(M-m)Q_{m,e}^{\vec\sigma} \notag \\
 & -\mu_{k+m+e}(k+m+e)Q_{m,e}^{\vec\sigma} \notag \\
 & -\lambda_{k+m+e}^{\text{a}}(\ell^0+m)Q_{m,e}^{\vec\sigma} \notag \\
 & -\lambda_{k+m+e}^{\text{c}}(k+m+e)Q_{m,e}^{\vec\sigma}
 \label{eq:dynaftellcol}
\end{align}
It can be verified that these equations generalize equations~(\ref{eq:dynaft1col1}--\ref{eq:dynaft1col2}) and equations (\ref{eq:dynaft2col1}--\ref{eq:dynaft2col4}).  In particular, on the second line we exclude the recolonization of the $\ell^0$ mainland species that have not yet undergone speciation on the island.  On the seventh line, when counting the endemic species that can undergo cladogenesis, we count twice the $k-\ell^0$ endemic species that are represented in the phylogeny.

Equations~(\ref{eq:dynaftellcol}) govern the dynamics in the time interval where $k$, the number of species in the data, and $\ell$, the number of colonizing lineages, take fixed values.  This time interval starts with the $k$-th event in the data, at time $t_k$ (note that $k$ increases by one at each event in the data, so that we can use $k$ to count the events).  This event is either a new colonization recorded in the data, or a cladogenesis in a lineage present in the data that colonized at an earlier time.  To formulate the initial conditions, we distinguish the following cases: \\[-15pt]

\begin{itemize}
\item The event at time $t_k$ is a colonization of a new mainland species, that is, a mainland species that has no colonization in the data before $t_k$.  The initial condition, connecting the solutions of equations~(\ref{eq:dynaftellcol}) before and after $t_k$, is
\begin{equation}
 Q_{m,e}^{\vec\sigma}(t_k) =
 \begin{cases}
   \gamma_{k+m+e-1}\,\diff t\;
   Q_{m,e}^{\vec\sigma^{\times}}(t_k) & \text{if $\sigma_\ell=0$} \\
   0 & \text{if $\sigma_\ell=1$}
 \end{cases}
 \label{eq:initellcol1}
\end{equation}
where the values of $k$ and $\ell$ are those after the colonization event.  Note that the values of $k$ and $\ell$ increase by one at a colonization event.
\item The event at time $t_k$ is a colonization of a mainland species that is already present in the data (i.e., it has one or more colonizations in the data before $t_k$), and the lineage initiated at the last colonization of this mainland species before $t_k$ has not branched before $t_k$.  Denote the index of the latter colonization by $i$.  We have to require that the lineage initiated at the $i$-th colonization has undergone speciation before $t_k$ (otherwise the new colonization would modify the time of the $i$-th colonization).  That is, we require that $\sigma_i=1$.  Hence, the initial condition is
\begin{equation}
 Q_{m,e}^{\vec\sigma}(t_k) =
 \begin{cases}
   \gamma_{k+m+e-1}\,\diff t\;
   Q_{m,e}^{\vec\sigma^{\times}}(t_k) & \text{if $\sigma_\ell=0$ and $\sigma_i=1$} \\
   0 & \text{otherwise}
 \end{cases}
 \label{eq:initellcol2}
\end{equation}
\item The event at time $t_k$ is a colonization of a mainland species that is already present in the data, and the lineage initiated at the last colonization of this mainland species before $t_k$ has at least one branching event before $t_k$.  Denoting the index of the latter colonization by $i$, we know from the presence of a branching event that immediately before $t_k$ the corresponding component $\sigma_i=1$ (more precisely, $Q_{m,e}^{\vec\sigma}=0$ if $\sigma_i=0$).  Hence, the initial condition is still given by equation~(\ref{eq:initellcol1}).
\item The event at time $t_k$ is a cladogenesis.  Denoting by $i$ the colonization that initiated the clade in which this cladogenesis event occurs, the initial condition is
\begin{equation}
 Q_{m,e}^{\vec\sigma}(t_k) = 
 \begin{cases}
   0 & \text{if $\sigma_i=0$} \\
   \lambda_{k+m+e-1}^{\text{c}}\,\diff t\;
   \Big( Q_{m,e}^{\sgmaa{i}{0}}(t_k)
        +Q_{m,e}^{\sgmaa{i}{1}}(t_k) \Big) & \text{if $\sigma_i=1$}
 \end{cases}
 \label{eq:initellcol3}
\end{equation}
We see that the two sets of equations for $\sigma_i=0$ and $\sigma_i=1$ are merged (or collapsed) into one set of equations for $\sigma_i=1$. Note that at a cladogenesis event the value of $\ell$ does not change, but the value of $k$ increases by one.
\end{itemize}

From a computational point of view it is important to note that we do not have to track probabilities $Q^{\vec\sigma}_{m,e}$ for all possible values of the vector $\vec\sigma$.  First, component $\sigma_i$ is only defined from the $i$-th colonization time onwards (undefined values are indicated by $\times$-marks in the table of Fig.~\ref{figcomp2}).  Second, for a colonization that initiates a clade with branching events, the corresponding component $\sigma_i$ has to be kept only until the first branching event.  After this first branching, component $\sigma_i$ is identical to one (see $\sigma_\text{A}$ and $\sigma_{\text{B}_1}$ in Fig.~\ref{figcomp2};  the numbers 1 printed in grey are imposed by the data).  Third, for a colonization that initiates a branch that remains non-endemic until the present time, the corresponding component $\sigma_i$ can be discarded entirely, because it is identical to zero from the $i$-th colonization time to the present time $t_\text{p}$ (see $\sigma_{\text{B}_2}$ and $\sigma_\text{C}$ in Fig.~\ref{figcomp2};  the numbers 0 printed in grey are imposed by the data).


%%%%%%%%%%%%%%%%%%%%%%%%%%%%%%%%%%%%%%%%%%%%%%%%%%%%%%%%%%%%%%%%%%%%%%%%%%%%%%%
\subsection*{Likelihood from solution of $Q$-equation}
%%%%%%%%%%%%%%%%%%%%%%%%%%%%%%%%%%%%%%%%%%%%%%%%%%%%%%%%%%%%%%%%%%%%%%%%%%%%%%%

We continue the above procedure, until reaching the present time $t_\text{p}$.  The likelihood can then be extracted from components $Q_{m,e}^{\vec\sigma}(t_\text{p})$ at the present time $t_\text{p}$.

To do so, we assume that we have complete information about the extant species.  That is, we assume that there are no other species on the island at the present time than those represented in the dataset.  This allows us to set the number of missing non-endemic and endemic species to zero, i.e., we impose that $m=e=0$ at time $t_\text{p}$.  The data also allow us to determine the vector $\vec\sigma_\text{p}$ at time $t_\text{p}$.  We set $\sigma_i=0$ for colonizations that correspond to lineages that are still non-endemic at the present time (represented by dashed lines in Fig.~\ref{figcomp1}).  For all other colonizations we set $\sigma_i=1$.  Finally, the likelihood is given by $Q_{0,0}^{\vec\sigma_\text{p}}(t_\text{p})$.

In practice we often do not have complete information.  For example, there might be large uncertainty about colonization times, or there might be missing species at the present time, i.e., species that are not represented in the dataset.  Some types of missing information can be easily dealt with, while other types requires a more thorough adaptation of the computational procedure.
