## Supplementary material for "The limits to ecological limits to diversification": Figures Tables and Supplementary Data and Scripts: Plot_CSvsIW.R

```
rm(list = ls())
dir <- 'd:/data/tex/Etienne et al 2019/'
source('d:/data/ms/ownfun/ICs.R')
library(latex2exp)
library(xtable)

pictype <- 'pdf'

pic <- function(name,pictype,width = 7,height = 5)
{
  if(pictype == 'png')
  {
    png(filename = paste0(name,'.png'))
  }
  if(pictype == 'pdf')
  {
    pdf(file = paste0(name,'.pdf'), width = width, height = height)
  }
  return(invisible(NULL))
}

# empirical data
data_52 <- read.csv(paste0(dir,'frogs_ML_results52_M1000.txt'),sep = '\t')
colnames(data_52) <- c('T','M','model','lam_c','mu','K','gam','lam_a',
                       'LL','df','NR','NR','rep','init_lam_c','init_mu',
                       'ínit_K','init_gam','init_lam_a')
data_52_sum <- NULL
data_52_DI <- subset(data_52, model == 'CS' & K == Inf)
data_52_sum <- rbind(data_52_sum,data_52_DI[which.max(data_52_DI[,'LL']),])
data_52_DD_CS <- subset(data_52, model == 'CS' & K != Inf & lam_a != 0)
data_52_sum <- rbind(data_52_sum,data_52_DD_CS[which.max(data_52_DD_CS[,'LL']),])
data_52_DD_CS_noA <- subset(data_52, model == 'CS' & K != Inf & lam_a == 0)
data_52_sum <- rbind(data_52_sum,data_52_DD_CS_noA[which.max(data_52_DD_CS_noA[,'LL']),])
data_52_DD_IW <- subset(data_52, model == 'IW' & K != Inf & lam_a != 0)
data_52_sum <- rbind(data_52_sum,data_52_DD_IW[which.max(data_52_DD_IW[,'LL']),])
data_52_DD_IW_noA <- subset(data_52, model == 'IW' & K != Inf & lam_a == 0)
data_52_sum <- rbind(data_52_sum,data_52_DD_IW_noA[which.max(data_52_DD_IW_noA[,'LL']),])
data_52_sum[,'AIC'] <- AIC(data_52_sum[,'LL'], data_52_sum[,'df'])
data_52_sum[,'AIC_weight'] <- ICweights(data_52_sum[,'LL'], data_52_sum[,'df'], 5, 'AIC')
data_52_sum[,'BIC'] <- BIC(data_52_sum[,'LL'], data_52_sum[,'df'],5)
data_52_sum[,'BIC_weight'] <- ICweights(data_52_sum[,'LL'], data_52_sum[,'df'], 5, 'BIC')
model <- c('DI','DD-CS','DD-CS-noA','IW-CS','IW-CS-noA')
data_52_sum_for_printing <- cbind(model,data_52_sum[,c(4:10,19:22)])
data_52_sum_for_printing[data_52_sum_for_printing[,'lam_a'] < 1E-4 & data_52_sum_for_printing[,'lam_a'] > 0,'lam_a'] = '0*'
data_52_sum_for_printing[data_52_sum_for_printing[,'K'] == Inf,K] = '$\infty$'
colnames(data_52_sum_for_printing)[2:6] <- c("$\\lambda_0^c$","$\\mu$","$K$","$\\gamma$","$\\lambda^a$")
colnames(data_52_sum_for_printing)[c(10,12)] <- c('AIC weight','BIC weight')
print(xtable(data_52_sum_for_printing, type = 'latex', digits = c(0,0,2,2,2,4,0,2,0,2,2,2,2)),
      file = paste0(dir,'data_52_sum.tex'),
      sanitize.text.function = function(x){x},
      include.rownames = FALSE)

loglik_CS <- data_52_sum[3,'LL'] #-208.672047620003
loglik_IW <- data_52_sum[5,'LL'] # -215.097677999268
CSminusIWdata <- loglik_CS - loglik_IW

parsCS <- data_52_sum[3,4:8] #c(0.4368767699838290, 0.1126373232769410, 36.4500023291870008, 0.0002187464194299,	0)
parsIW <- data_52_sum[5,4:8] #c(0.4012298101438560, 0.1699521120864120, 131.9595140113719935, 0.0003487955068717,	0)

m_pool <- 1000

boots <- read.table(paste0(dir,'out_simulation_M1000.txt'),header = TRUE)
boots <- boots[order(boots[,1]),]
boots <- boots[,1:15]
colnames(boots) <- c('the_island','the_dataset','island_age','M','the_init','model','model_type','methode','R','lam_c','mu','K','gam','lam_a','loglik')

logliks_CS <- matrix(nrow = max(boots$the_island),ncol = 26)
logliks_CS[,1] <- 1:max(boots$the_island)

logliks_IW <- matrix(nrow = max(boots$the_island),ncol = 26)
logliks_IW[,1] <- 1:max(boots$the_island)

# Subsetting and Removing duplicates
CS_to_CS <- subset(boots,the_dataset == 'CSdata' & model_type == 'CS')
CS_to_CS <- CS_to_CS[!duplicated(CS_to_CS$the_island),]
cat('There are ',length(CS_to_CS[,1]), 'CS inferences on CS sims \n')
which_CS_to_CS_NA <- which(is.na(CS_to_CS$loglik))
cat('Of these there are ',length(which_CS_to_CS_NA), 'NA \n')
cat('These are: ',sort(CS_to_CS$the_island[which_CS_to_CS_NA]), '\n')
IW_to_CS <- subset(boots,the_dataset == 'CSdata' & model_type == 'IW')
IW_to_CS <- IW_to_CS[!duplicated(IW_to_CS$the_island),]
cat('There are ',length(IW_to_CS[,1]), 'IW inferences on CS sims \n')
which_IW_to_CS_NA <- which(is.na(IW_to_CS$loglik))
cat('Of these there are ',length(which_IW_to_CS_NA), 'NA \n')
cat('These are: ',sort(IW_to_CS$the_island[which_IW_to_CS_NA]), '\n')
CS_to_IW <- subset(boots,the_dataset == 'IWdata' & model_type == 'CS')
CS_to_IW <- CS_to_IW[!duplicated(CS_to_IW$the_island),]
cat('There are ',length(CS_to_CS[,1]), 'CS inferences on IW sims \n')
which_CS_to_IW_NA <- which(is.na(CS_to_IW$loglik))
cat('Of these there are ',length(which_CS_to_IW_NA), 'NA \n')
cat('These are: ',sort(CS_to_IW$the_island[which_CS_to_IW_NA]), '\n')
IW_to_IW <- subset(boots,the_dataset == 'IWdata' & model_type == 'IW')
IW_to_IW <- IW_to_IW[!duplicated(IW_to_IW$the_island),]
cat('There are ',length(IW_to_IW[,1]), 'IW inferences on IW sims \n')
which_IW_to_IW_NA <- which(is.na(IW_to_IW$loglik))
cat('Of these there are ',length(which_IW_to_IW_NA), 'NA \n')
cat('These are: ',sort(IW_to_IW$the_island[which_IW_to_IW_NA]), '\n')

for (i in 1:max(boots$the_island)){
  logliks_CS[i,3:8]  <- as.numeric(CS_to_CS[which(CS_to_CS[,'the_island'] == i)[1],c('lam_c','mu','K','gam','lam_a','loglik')])
  logliks_CS[i,9:14] <- as.numeric(IW_to_CS[which(IW_to_CS[,'the_island'] == i)[1],c('lam_c','mu','K','gam','lam_a','loglik')])
}

for (i in 1:max(boots$the_island)){
  logliks_IW[i,3:8]  <- as.numeric(CS_to_IW[which(CS_to_IW[,'the_island'] == i)[1],c('lam_c','mu','K','gam','lam_a','loglik')])
  logliks_IW[i,9:14] <- as.numeric(IW_to_IW[which(IW_to_IW[,'the_island'] == i)[1],c('lam_c','mu','K','gam','lam_a','loglik')])
}

colz <- c('the_island','N_spec','lam_c_CS','mu_CS','K_CS','gam_CS','lam_a_CS','loglik_CS','lam_c_IW','mu_IW',
        'K_IW','gam_IW','lam_a_IW','loglik_IW','best_model','loglik_diff','loglik_diff2','< -2','between2','> 2','AICweights1','AICweights2',
        'N_cols','Specs_per_col','AIC_CS','AIC_IW')

colnames(logliks_CS) <- colz
colnames(logliks_IW) <- colz

load(paste(dir,'/frogs_sim_CS_T30_M1000_R5000.Rdata',sep=""))
not_present_CS <- c()
for(i in 1:length(frogs_sims))
{
  if(frogs_sims[[i]][[1]]$not_present == 1000)
  {
    not_present_CS <- append(not_present_CS, i)
  }
}

frogs_sims_CS <- frogs_sims[-not_present_CS]
number_spec_CS <- NULL
number_cols_CS <- NULL
for(i in 1:length(frogs_sims_CS))
{
  the_island <- frogs_sims_CS[[i]]
  s <- NULL
  number_cols_CS <- append(number_cols_CS, length(the_island) - 1)
  for(j in 2:length(the_island))
  {
    if(is.null(the_island[[j]]$all_colonisations))
    {
      s <- c(s, length(the_island[[j]]$branching_times) - 1)
    } else
    {
      for(k in 1:length(the_island[[j]]$all_colonisations))
      {
        s <- c(s, length(the_island[[j]]$all_colonisations[[k]]$event_times) - 1)
      }
    }
  }
  number_spec_CS <- append(number_spec_CS,sum(s))
}
logliks_CS[,'N_spec'] <- number_spec_CS[1:1000]
logliks_CS[,'N_cols'] <- number_cols_CS[1:1000]

load(paste(dir,'/frogs_sim_IW_T30_M1000_R5000.Rdata',sep=""))
not_present_IW <- c()
for(i in 1:length(frogs_sims))
{
  if(frogs_sims[[i]][[1]]$not_present == 1000)
  {
    not_present_IW <- append(not_present_IW, i)
  }
}
frogs_sims_IW <- frogs_sims[-not_present_IW]
number_spec_IW <- NULL
number_cols_IW <- NULL
for(i in 1:length(frogs_sims_IW))
{
  the_island <- frogs_sims_IW[[i]]
  s <- NULL
  number_cols_IW <- append(number_cols_IW, length(the_island) - 1)
  for(j in 2:length(the_island))
  {
    if(is.null(the_island[[j]]$all_colonisations))
    {
      s <- c(s, length(the_island[[j]]$branching_times) - 1)
    } else
    {
      for(k in 1:length(the_island[[j]]$all_colonisations))
      {
        s <- c(s, length(the_island[[j]]$all_colonisations[[k]]$event_times) - 1)
      }
    }
  }
  number_spec_IW <- append(number_spec_IW,sum(s))
}
logliks_IW[,'N_spec'] <- number_spec_IW[1:1000]
logliks_IW[,'N_cols'] <- number_cols_IW[1:1000]

for(i in 1:nrow(logliks_CS)){
  the_logliks <- c(logliks_CS[i,'loglik_CS'],logliks_CS[i,'loglik_IW'])
  logliks_CS[i,'loglik_diff'] <- logliks_CS[i,'loglik_CS'] - logliks_CS[i,'loglik_IW']
  logliks_CS[i,'best_model']  <- as.numeric(which(the_logliks == max(the_logliks)))

  if(logliks_CS[i,'loglik_diff'] < (-2)){
    logliks_CS[i,'< -2'] <- 1
    logliks_CS[i,'between2'] <- 0
    logliks_CS[i,'> 2'] <- 0
  }

  if(logliks_CS[i,'loglik_diff'] > -2 & logliks_CS[i,'loglik_diff'] < 2){
    logliks_CS[i,'< -2'] <- 0
    logliks_CS[i,'between2'] <- 1
    logliks_CS[i,'> 2'] <- 0
  }

  if(logliks_CS[i,'loglik_diff'] > 2){
    logliks_CS[i,'< -2'] <- 0
    logliks_CS[i,'between2'] <- 0
    logliks_CS[i,'> 2'] <- 1
  }

  AICweights <- ICweights(the_logliks,c(4,4),100,'AIC')
  AICss <- AIC(the_logliks,c(4,4))
  logliks_CS[i,'AICweights1'] <- AICweights[1]
  logliks_CS[i,'AICweights2'] <- AICweights[2]
  logliks_CS[i,'AIC_CS'] <- AICss[1]
  logliks_CS[i,'AIC_IW'] <- AICss[2]
}

for(i in 1:nrow(logliks_IW)){
  the_logliks <- c(logliks_IW[i,'loglik_CS'],logliks_IW[i,'loglik_IW'])
  loglik_diff <- logliks_IW[i,'loglik_IW'] - logliks_IW[i,'loglik_CS']
  loglik_diff2 <- logliks_IW[i,'loglik_CS'] - logliks_IW[i,'loglik_IW']

  logliks_IW[i,'loglik_diff'] <- loglik_diff
  logliks_IW[i,'loglik_diff2'] <- loglik_diff2

  logliks_IW[i,'best_model']  <- as.numeric(which(the_logliks == max(the_logliks)))

  if(logliks_IW[i,'loglik_diff'] < (-2)){
    logliks_IW[i,'< -2'] <- 1
    logliks_IW[i,'between2'] <- 0
    logliks_IW[i,'> 2'] <- 0
  }

  if(logliks_IW[i,'loglik_diff'] > -2 & logliks_IW[i,'loglik_diff'] < 2){
    logliks_IW[i,'< -2'] <- 0
    logliks_IW[i,'between2'] <- 1
    logliks_IW[i,'> 2'] <- 0
  }

  if(logliks_IW[i,'loglik_diff'] > 2){
    logliks_IW[i,'< -2'] <- 0
    logliks_IW[i,'between2'] <- 0
    logliks_IW[i,'> 2'] <- 1
  }

  AICweights <- ICweights(the_logliks,c(4,4),100,'AIC')
  AICss <- AIC(the_logliks,c(4,4))
  logliks_IW[i,'AICweights1'] <- AICweights[1]
  logliks_IW[i,'AICweights2'] <- AICweights[2]
  logliks_IW[i,'AIC_CS'] <- AICss[1]
  logliks_IW[i,'AIC_IW'] <- AICss[2]
}

logliks_CS[,'Specs_per_col'] <- logliks_CS[,'N_spec']/logliks_CS[,'N_cols']
logliks_IW[,'Specs_per_col'] <- logliks_IW[,'N_spec']/logliks_IW[,'N_cols']

cat('Percentage of CS sims where IW was chosen as best model:',length(which(logliks_CS[,'best_model'] == 2))/nrow(logliks_CS)*100,'\n')
cat('Percentage of IW sims where CS was chosen as best model:',length(which(logliks_IW[,'best_model'] == 1))/nrow(logliks_IW)*100,'\n')

####
### PLOT CS PARAMETER DISTRIBUTION (CS pars under CS sims)

cex_lab <- 1.5
cex_size <- 0.8

pic(name = paste0(dir,'Bootstrap_pars_under_CS'),pictype = pictype)
lam_c_min <- 0
lam_c_max <- 1.5
dens_min <- 0
dens_max <- 4
par(mfrow = c(2,2),cex = cex_size,cex.lab = cex_lab,mar = c(4,5,1,2))
hist(CS_to_CS$lam_c,
     freq = F, breaks = c(seq(lam_c_min,lam_c_max,length.out = 25),10^10),
     xlab = TeX('$\\lambda_0 ^ c$'),
     main = NULL,
     col  = 'cornsilk3',
     border = 'cornsilk3',
     ylim = c(dens_min, dens_max),
     xlim = c(lam_c_min,lam_c_max))
##
med <- median(CS_to_CS$lam_c, na.rm = TRUE)
obs <- as.numeric(parsCS[1])
arrows(med, dens_max, med, 0, col = 'black',length = 0,lwd = 1.5)
arrows(obs, dens_max/4, obs, 0, col = 'blue',length = 0.05,lwd = 1.5)
box(bty = 'l')

mu_min <- 0
mu_max <- 0.8
dens_min <- 0
dens_max <- 12
hist(CS_to_CS$mu,
     freq = FALSE, breaks = c(seq(mu_min,mu_max,length.out = 25),10^10),
     xlab = TeX('$\\mu$'),
     main = NULL,
     col = 'cornsilk3',
     border = 'cornsilk3',
     ylim = c(dens_min, dens_max),
     xlim = c(mu_min,mu_max),
     axes = FALSE)
axis(side = 1, at = seq(mu_min,mu_max,0.2), labels = seq(mu_min,mu_max,0.2))
axis(side = 2, at = seq(dens_min,dens_max,2), labels = c(0,2,4,6,8,10,12))
med <- median(CS_to_CS$mu,na.rm = TRUE)
obs <- as.numeric(parsCS[2])
arrows(med, dens_max, med, 0, col = 'black',length = 0,lwd = 1.5)
arrows(obs, dens_max/4, obs, 0, col = 'blue',length = 0.05,lwd = 1.5)
box(bty = 'l')

K_min <- 0
K_max <- 100
dens_min <- 0
dens_max <- 0.1
hist(CS_to_CS$K,
     freq = FALSE, breaks = c(seq(K_min,K_max,length.out = 25),10^10),
     xlab = TeX('$\\textit{K}$'),
     main = NULL,
     col = 'cornsilk3',
     border  = 'cornsilk3',
     ylim = c(dens_min, dens_max),
     xlim = c(K_min,K_max))
med <- median(CS_to_CS$K,na.rm = T)
obs <- as.numeric(parsCS[3])
arrows(med, dens_max, med, 0, col = 'black',length = 0,lwd = 1.5)
arrows(obs, dens_max/4, obs, 0, col = 'blue',length = 0.05,lwd = 1.5)
box(bty = 'l')

gam_min <- 0
gam_max <- 0.002
dens_min <- 0
dens_max <- 4000
hist(CS_to_CS$gam,
     freq = FALSE, breaks = c(seq(gam_min,gam_max,length.out = 25),10^10),
     xlab = TeX('$\\gamma_0$'),
     main = NULL,
     col = 'cornsilk3',
     border = 'cornsilk3',
     ylim = c(dens_min, dens_max),
     xlim = c(gam_min,gam_max),
     axes = FALSE)
axis(side = 1, at = c(0,0.0005,0.001,0.0015,0.002), labels = c(0,'0.0005',0.001,0.0015,0.002))
axis(side = 2, at = c(0,1000,2000,3000,4000), labels = c(0,'',2000,'',4000))
med <- median(CS_to_CS$gam,na.rm = TRUE)
obs <- as.numeric(parsCS[4])
arrows(med, dens_max, med, 0, col = 'black',length = 0,lwd = 1.5)
arrows(obs, dens_max/4, obs, 0, col = 'blue',length = 0.05,lwd = 1.5)
box(bty = 'l')

dev.off()

### PLOT IW parameters under CS sims

pic(name = paste0(dir,'Bootstrap_IW_pars_under_CS'),pictype = pictype)
par(mfrow = c(2,2),cex = cex_size,cex.lab = cex_lab,mar = c(4,5,1,2))
lam_c_min <- 0
lam_c_max <- 1.5
dens_min <- 0
dens_max <- 3.5
hist(IW_to_CS$lam_c,
     freq = FALSE, breaks = c(seq(lam_c_min,lam_c_max,length.out = 25),10^10),
     xlab = TeX('$\\lambda_0 ^ c$'),
     main = NULL,
     col  = 'cornsilk3',
     border = 'cornsilk3',
     ylim = c(dens_min, dens_max),
     xlim = c(lam_c_min,lam_c_max))
med <- median(IW_to_CS$lam_c, na.rm = TRUE)
arrows(med, dens_max, med, 0, col = 'black',length = 0,lwd = 1.5)
box(bty = 'l')

mu_min <- 0
mu_max <- 0.8
dens_min <- 0
dens_max <- 7
hist(IW_to_CS$mu,
     freq = FALSE, breaks = c(seq(mu_min,mu_max,length.out = 25),10^10),
     xlab = TeX('$\\mu$'),
     main = NULL,
     col = 'cornsilk3',
     border = 'cornsilk3',
     ylim = c(dens_min, dens_max),
     xlim = c(mu_min,mu_max))
med <- median(IW_to_CS$mu, na.rm = TRUE)
arrows(med, dens_max, med, 0, col = 'black',length = 0,lwd = 1.5)
box(bty = 'l')

K_min <- 0
K_max <- 400
dens_min <- 0
dens_max <- 0.006
hist(IW_to_CS$K,
     freq = FALSE, breaks = c(seq(K_min,K_max,length.out = 25),10^10),
     xlab = TeX('$\\textit{K}$'),
     main = NULL,
     col = 'cornsilk3',
     border  = 'cornsilk3',
     ylim = c(dens_min, dens_max),
     xlim = c(K_min,K_max))
med <- median(IW_to_CS$K, na.rm = TRUE)
arrows(med, dens_max, med, 0, col = 'black',length = 0,lwd = 1.5)
box(bty = 'l')

gam_min <- 0
gam_max <- 0.003
dens_min <- 0
dens_max <- 2000
hist(IW_to_CS$gam,
     freq = FALSE, breaks = c(seq(gam_min,gam_max,length.out = 25),10^10),
     xlab = TeX('$\\gamma_0$'),
     main = NULL,
     col = 'cornsilk3',
     border = 'cornsilk3',
     yaxt = 'n',
     ylim = c(dens_min, dens_max),
     xlim = c(gam_min,gam_max),
     axes = FALSE)
axis(side = 1, at = seq(gam_min,gam_max,0.0005), labels = c(0,'',0.001, '',0.002,'',0.003))
axis(side = 2, at = c(dens_min,dens_max,1000), labels = c(dens_min,1000,dens_max))
med <- median(IW_to_CS$gam, na.rm = TRUE)
arrows(med, dens_max, med, 0, col = 'black',length = 0,lwd = 1.5)
box(bty = 'l')

dev.off()

### PLOT IW PARAMETER DISTRIBUTION (IW pars under IW sims)

pic(name = paste0(dir,'Bootstrap_pars_under_IW'),pictype = pictype)
par(mfrow = c(2,2),cex = cex_size,cex.lab = cex_lab,mar = c(4,5,1,2))
lam_c_min <- 0
lam_c_max <- 1.5
dens_min <- 0
dens_max <- 3.5
hist(IW_to_IW$lam_c,
     freq = FALSE, breaks = c(seq(lam_c_min,lam_c_max,length.out = 25),10^10),
     xlab = TeX('$\\lambda_0 ^ c$'),
     main = NULL,
     col = 'cornsilk3',
     border = 'cornsilk3',
     ylim = c(dens_min, dens_max),
     xlim = c(lam_c_min,lam_c_max))
med <- median(IW_to_IW$lam_c,na.rm = TRUE)
obs <- as.numeric(parsIW[1])
arrows(med, dens_max, med, 0, col = 'black',length = 0,lwd = 1.5)
arrows(obs, dens_max/4, obs, 0, col = 'blue',length = 0.05,lwd = 1.5)
box(bty = 'l')

mu_min <- 0
mu_max <- 0.8
dens_min <- 0
dens_max <- 8
hist(IW_to_IW$mu,
     freq = FALSE, breaks = c(seq(mu_min,mu_max,length.out = 25),10^10),
     xlab = TeX('$\\mu$'),
     main = NULL,
     col = 'cornsilk3',
     border = 'cornsilk3',
     ylim = c(dens_min, dens_max),
     xlim = c(mu_min,mu_max))
med <- median(IW_to_IW$mu,na.rm = TRUE)
obs <- as.numeric(parsIW[2])
arrows(med, dens_max, med, 0, col = 'black',length = 0,lwd = 1.5)
arrows(obs, dens_max/4, obs, 0, col = 'blue',length = 0.05,lwd = 1.5)
box(bty = 'l')

K_min <- 0
K_max <- 300
dens_min <- 0
dens_max <- 0.02
hist(IW_to_IW$K,
     freq = FALSE, breaks = c(seq(K_min,K_max,length.out = 25),10^10),
     xlab = TeX('$\\textit{K}$'),
     main = NULL,
     col = 'cornsilk3',
     border = 'cornsilk3',
     ylim = c(dens_min, dens_max),
     xlim = c(K_min,K_max))
med <- median(IW_to_IW$K,na.rm = TRUE)
obs <- as.numeric(parsIW[3])
arrows(med, dens_max, med, 0, col = 'black',length = 0,lwd = 1.5)
arrows(obs, dens_max/4, obs, 0, col = 'blue',length = 0.05,lwd = 1.5)
box(bty = 'l')

gam_min <- 0
gam_max <- 0.003
dens_min <- 0
dens_max <- 2000
hist(IW_to_IW$gam,
     freq = FALSE, breaks = c(seq(gam_min,gam_max,length.out = 25),10^10),
     xlab = TeX('$\\gamma_0$'),
     main = NULL,
     col = 'cornsilk3',
     border = 'cornsilk3',
     yaxt = 'n',
     ylim = c(dens_min, dens_max),
     xlim = c(gam_min,gam_max),
     axes = FALSE)
axis(side = 1, at = seq(gam_min,gam_max,0.0005), labels = c(0,'',0.001,'',0.002,'',0.003))
axis(side = 2, at = seq(dens_min,dens_max,1000), labels = seq(dens_min,dens_max,1000))
med <- median(IW_to_IW$gam,na.rm = TRUE)
obs <- as.numeric(parsIW[4])
arrows(med, dens_max, med, 0, col = 'black',length = 0,lwd = 1.5)
arrows(obs, dens_max/4, obs, 0, col = 'blue',length = 0.05,lwd = 1.5)
box(bty = 'l')

dev.off()

### Plot CS pars under IW sims

pic(name = paste0(dir,'Bootstrap_CS_pars_under_IW'),pictype = pictype)
par(mfrow = c(2,2),cex = cex_size,cex.lab = cex_lab,mar = c(4,5,1,2))
lam_c_min <- 0
lam_c_max <- 1.5
dens_min <- 0
dens_max <- 3.5
hist(CS_to_IW$lam_c,
     freq = FALSE, breaks = c(seq(lam_c_min,lam_c_max,length.out = 25),10^10),
     xlab = TeX('$\\lambda_0 ^ c$'),
     main = NULL,
     col = 'cornsilk3',
     border = 'cornsilk3',
     ylim = c(dens_min,dens_max),
     xlim = c(lam_c_min,lam_c_max))
med <- median(CS_to_IW$lam_c, na.rm = TRUE)
arrows(med, dens_max, med, 0, col = 'black',length = 0,lwd = 1.5)
box(bty = 'l')

mu_min <- 0
mu_max <- 0.8
dens_min <- 0
dens_max <- 6
hist(CS_to_IW$mu,
     freq = FALSE, breaks = c(seq(mu_min,mu_max,length.out = 25),10^10),
     xlab = TeX('$\\mu$'),
     main = NULL,
     col = 'cornsilk3',
     border = 'cornsilk3',
     ylim = c(dens_min,dens_max),
     xlim = c(mu_min,mu_max))
med <- median(CS_to_IW$mu, na.rm = TRUE)
arrows(med, dens_max, med, 0, col = 'black',length = 0,lwd = 1.5)
box(bty = 'l')

K_min <- 0
K_max <- 300
dens_min <- 0
dens_max <- 0.01
hist(CS_to_IW$K,
     freq = FALSE, breaks = c(seq(K_min,K_max,length.out = 25),10^10),
     xlab = TeX('$\\textit{K}$'),
     main = NULL,
     col = 'cornsilk3',
     border = 'cornsilk3',
     ylim = c(dens_min,dens_max),
     xlim = c(K_min,K_max),
     axes = FALSE)
axis(side = 1, at = seq(K_min,K_max,50), labels = seq(K_min,K_max,50))
axis(side = 2, at = seq(0,0.01,0.002), labels = c(0,'',0.004,'',0.008,''))
med <- median(CS_to_IW$K, na.rm = TRUE)
arrows(med, dens_max, med, 0, col = 'black',length = 0,lwd = 1.5)
box(bty = 'l')

gam_min <- 0
gam_max <- 0.003
dens_min <- 0
dens_max <- 4000
hist(CS_to_IW$gam,
     freq = FALSE, breaks = c(seq(gam_min,gam_max,length.out = 25),10^10),
     xlab = TeX('$\\gamma_0$'),
     main = NULL,
     col = 'cornsilk3',
     border = 'cornsilk3',
     ylim = c(dens_min,dens_max),
     xlim = c(gam_min,gam_max),
     axes = FALSE)
axis(side = 1, at = seq(gam_min,gam_max,0.0005), labels = c(0,'',0.001,'',0.002,'',0.003))
axis(side = 2, at = c(0,1000,2000,3000,4000), labels = c(0,'',2000,'',4000))
med <- median(CS_to_IW$gam, na.rm = TRUE)
arrows(med, dens_max, med, 0, col = 'black',length = 0,lwd = 1.5)
box(bty = 'l')

dev.off()


### PLOT SATURATION CS and IW

pic(name = paste0(dir,'Saturation'),pictype = pictype, height = 4)
par(mfrow = c(1,2),cex = cex_size,cex.lab = cex_lab,cex.main = 1.3,mar = c(4,5,3,2))
dens_min <- 0
dens_max <- 200
statistic <- logliks_CS[,'N_spec']/(ceiling(CS_to_CS$K) * logliks_CS[,'N_cols'])
statistic <- subset(statistic, statistic < 2)
hist(statistic,breaks = 25,
     xlab = TeX('Total richness / Total $\\textit{K}$'),
     col = 'cornsilk3',
     main = 'CS',
     border = 'cornsilk3',
     ylim = c(dens_min,dens_max))
obs <- 65/(as.numeric(parsCS[3]) * 5)
med <- median(statistic, na.rm = TRUE)
arrows(med, dens_max, med, 0, col = 'black',length = 0,lwd = 1.5)
arrows(obs, dens_max/4, obs, 0, col = 'blue',length = 0.05,lwd = 1.5)
box(bty = 'l')

statistic <- logliks_IW[,'N_spec']/ceiling(IW_to_IW$K)
statistic <- subset(statistic, statistic < 2)
hist(statistic,breaks = 25,
     xlab = TeX('Total richness / $\\textit{K}$'),
     col = 'cornsilk3',
     main = 'IW',
     border = 'cornsilk3',
     ylim = c(0,200))
obs <- 65/as.numeric(parsIW[3])
med <- median(statistic, na.rm = TRUE)
arrows(med, dens_max, med, 0, col = 'black',length = 0,lwd = 1.5)
arrows(obs, dens_max/4, obs, 0, col = 'blue',length = 0.05,lwd = 1.5)
box(bty = 'l')

dev.off()

#hist(statistic,breaks = 25,
#     main = TeX('Size largest clade / $\\textit{K}$'),col = 'cornsilk3',xlab = 'Saturation',
#     border = 'cornsilk3')
#med <- median(statistic, na.rm = TRUE)
#arrows(med, arrow_bottom,med, 0,col = 'blue',length = 0.05,lwd = 1.5)
#arrows(28/36, arrow_bottom,28/36, 0,col = 'black',length = 0.05,lwd = 1.5)

#statistic <- IW_to_IW$size_largest_clade_IW/IW_to_IW$K
#hist(statistic,breaks = 25,
#     main = TeX('Size largest clade / \\textit{K}'),col = 'cornsilk3',xlab = 'Saturation',
#     border = 'cornsilk3')
#med <- median(statistic, na.rm = TRUE)
#arrows(med, arrow_bottom,med, 0,col = 'blue',length = 0.05,lwd = 1.5)
#arrows(28/131, arrow_bottom,28/131, 0,col = 'black',length = 0.05,lwd = 1.5)

####
## PLOT Loglik delta vs species richness

par(mfrow = c(1,2))
plot(logliks_CS[,'N_spec'],logliks_CS[,'loglik_diff'],
     xlab = 'Number of species',ylab = 'loglik CS - loglik IW',
     main = 'CS datasets',
     pch = 16,cex = 0.6)
abline(h = CSminusIWdata,col = 'black',lwd = 1)
abline(h = 0,lty = 1, col = 'blue')
abline(h = 2,lty = 2, col = 'blue')
abline(h = -2,lty = 2, col = 'blue')
plot(logliks_IW[,'N_spec'],-logliks_IW[,'loglik_diff'],
     xlab = 'Number of species',ylab = 'loglik IW - loglik CS',
     main = 'IW datasets',
     pch = 16,cex = 0.6)
abline(h = -CSminusIWdata,col = 'black',lwd = 1)
abline(h = 0,lty = 1, col = 'blue')
abline(h = 2,lty = 2, col = 'blue')
abline(h = -2,lty = 2, col = 'blue')
box(bty = 'l')

####
## PLOT Loglik delta vs Ncols

par(mfrow = c(1,1))
plot(logliks_CS[,'N_cols'],logliks_CS[,'loglik_diff'],
     xlab = 'Number of colonisations',ylab = 'loglik CS - loglik IW',
     main = 'Loglik CS minus loglik IW, CS datasets',
     cex = 0.5)
abline(h = CSminusIWdata,col = 'black',lwd = 1)
abline(h = 0)
abline(h = 2,lty = 2)
abline(h = -2,lty = 2)
box(bty = 'l')

####
## PLOT Loglik delta vs Specs per col

par(mfrow = c(1,1))
plot(logliks_CS[,'Specs_per_col'],logliks_CS[,'loglik_diff'],xlab = 'Number of species per colonisation',
     ylab = 'loglik CS - loglik IW',main = 'Loglik CS minus loglik IW, CS datasets',
     pch = 16,cex = 0.5)
abline(h = CSminusIWdata,col = 'black',lwd = 1)
abline(h = 0)
abline(h = 0)
abline(h = 2,lty = 2)
abline(h = -2,lty = 2)
box(bty = 'l')

####
## HISTS Loglik delta OVERLAPPED
pic(name = paste0(dir,'LL'),pictype = pictype)
par(mfrow = c(1,1))
ycol <- rgb(0, 0, 50, maxColorValue = 255, alpha = 125, names = "green")

hist(logliks_CS[,'loglik_diff'],breaks = 30,xlab = 'Loglik CS - Loglik IW',
     ylim = c(0,250),col = 'cornsilk3',main = '',xlim = c(-20,40))
hist(logliks_IW[,'loglik_diff2'],breaks = 20,add = T,col = ycol)

arrows(quantile(logliks_CS[,'loglik_diff'],0.05), -10, quantile(logliks_CS[,'loglik_diff'],0.05), -2, xpd = TRUE,col = 'cornsilk3',length = 0.05,lwd = 2)
arrows(quantile(logliks_IW[,'loglik_diff2'],0.95), -10, quantile(logliks_IW[,'loglik_diff2'],0.95), -2, xpd = TRUE,col = ycol,length = 0.05,lwd = 2)
arrows(CSminusIWdata, -10, CSminusIWdata, -2, xpd = TRUE,col = 'black',length = 0.05,lwd = 2)
box(bty = 'l')

xl <- 11
yl <- 230
legend(x = xl,y = yl,legend = c('CS data', 'IW data'),pch = 15,bty = 'n',
       col = c('cornsilk3',ycol),cex = 1.2)
arrows(xl + 2, yl - 73, xl + 2, yl - 73 + 8, xpd = TRUE,col = 'black',length = 0.05,lwd = 2)
text(x = xl + 4, y = yl - 70, adj = 0, 'empirical value')
arrows(xl + 2, yl - 91.5, xl + 2, yl - 91.5 + 8, xpd = TRUE,col = 'cornsilk3',length = 0.05,lwd = 2)
text(x = xl + 4, y = yl - 88, adj = 0, TeX('5^{th} percentile CS simulations'))
arrows(xl + 2, yl - 110, xl + 2, yl - 110 + 8, xpd = TRUE,col = ycol,length = 0.05,lwd = 2)
text(x = xl + 4, y = yl - 106, adj = 0, TeX('95^{th} percentile IW simulations'))
box(bty = 'l')

dev.off()

#Similar with ggplot2

#library(ggplot2)
#CS <- logliks_CS[,'loglik_diff']
#IW <- logliks_IW[,'loglik_diff2']

#q5 <- quantile(logliks_CS[,'loglik_diff'],0.05)
#q5
#cat('5% quantile in CS logliks:',q5,'\n')

#q95 <- quantile(logliks_IW[,'loglik_diff2'],0.95)
#q95
#cat('95% quantile in IW logliks:', q95,'\n')

#obs <- CSminusIWdata
#dati <- data.frame(loglik_difference = factor(rep(c("CS data","IW data"), c(length(CS),length(IW)))),
#                   rating = c(CS,IW))
#pic(name = paste0(dir,'LL2'),pictype = pictype)
#ggplot(dati, aes(x = rating, fill = loglik_difference))  +
#  geom_histogram(binwidth = 2, position = "dodge")  +
#  theme(panel.grid.major = element_blank(), panel.grid.minor = element_blank(),
#        panel.background = element_blank(), axis.line = element_line(colour = "black"),
#        text = element_text(size = 18),axis.title.y = element_text(margin = margin(t = 0, r = 10, b = 0, l = 0)),
#        axis.title.x = element_text(margin = margin(t = 10, r = 0, b = 0, l = 0)),
#        axis.text.x = element_text(size = 18),
#        axis.text.y = element_text(size = 18),
#        legend.position = c(0.7, 0.5),legend.title = element_text(colour = "white"))  +
#  scale_fill_brewer(palette = "Pastel1")  +
#  ylab("Frequency")  +
#  xlab("Loglik CS - Loglik IW")  +
#  geom_segment(aes(x = q5, y = -25, xend = q5, yend = 0),
#    lineend = "butt", linejoin = "round", color = 'LightGrey',
#    size = 1, arrow = arrow(length = unit(0.1, "inches"))
#  )  +
#  geom_segment(aes(x = q95, y = -25, xend = q95, yend = 0),
#      lineend = "butt", linejoin = "round", color = 'DarkGrey',
#      size = 1, arrow = arrow(length = unit(0.1, "inches"))
#  )  +
#  geom_segment(aes(x = obs, y = -25, xend = obs, yend = 0),
#               lineend = "butt", linejoin = "round", color = 'Black',
#               size = 1, arrow = arrow(length = unit(0.1, "inches"))
#  )
#dev.off()

#p value
p_value <- length(which(logliks_IW[,'loglik_diff2'] > CSminusIWdata))/1000

#Power to detect CS (5% error):
pow_CS <- 1 - length(which(logliks_CS[,'loglik_diff'] < quantile(logliks_IW[,'loglik_diff2'],0.95)))/length(logliks_CS[,'loglik_diff'])
cat('Power to detect CS:', pow_CS,'\n')

#Power to detect IW (5% error):
pow_IW <- 1 - length(which(logliks_IW[,'loglik_diff2'] > quantile(logliks_CS[,'loglik_diff'],0.05)))/length(logliks_IW[,'loglik_diff2'])
cat('Power to detect IW:',pow_IW,'\n')

## Incorrectly preferring IW with CS data
#simply compare likelihoods
logdiff <- logliks_CS[,'loglik_CS'] - logliks_CS[,'loglik_IW']
error_IW_on_CS_highest_likelihood <- length(which(logdiff < 0))/nrow(logliks_CS)*100
cat('Error of choosing IW on CS sims using highest likelihood:',error_IW_on_CS_highest_likelihood,'\n')

#at least 2 loglik difference
logdiff2 <- logliks_CS[,'loglik_CS'] - logliks_CS[,'loglik_IW']
#incorrectly inferred:
error_IW_on_CS_2_units_likelihood <- length(which(logdiff2 < (-2)))/nrow(logliks_CS)*100
cat('Error of choosing IW on CS sims using 2 units difference in likelihood:',error_IW_on_CS_2_units_likelihood,'\n')
#selected correctly
correct_IW_on_CS_2_units_likelihood <- length(which(logdiff2 > (2)))/nrow(logliks_CS)*100
cat('Correctly choosing CS using 2 units difference likelihood:',correct_IW_on_CS_2_units_likelihood,'\n')
#undecided
undecided_IW_on_CS_2_units_likelihood <- 100 - error_IW_on_CS_2_units_likelihood - correct_IW_on_CS_2_units_likelihood
cat('Undecided using 2 units difference likelihood:',undecided_IW_on_CS_2_units_likelihood,'\n')

## Incorrectly preferring CS with IW data
#simply compare likelihoods
logdiff <- logliks_IW[,'loglik_IW'] - logliks_IW[,'loglik_CS']
error_CS_on_IW_highest_likelihood <- length(which(logdiff < 0))/nrow(logliks_IW)*100
cat('Error of choosing CS on IW sims using highest likelihood:',error_CS_on_IW_highest_likelihood,'\n')

#at least 2 loglik difference
logdiff2 <- logliks_IW[,'loglik_IW'] - logliks_IW[,'loglik_CS']
error_CS_on_IW_2_units_likelihood <- length(which(logdiff2 < (-2)))/nrow(logliks_IW)*100
cat('Correctly choosing IW using 2 units difference likelihood:',error_CS_on_IW_2_units_likelihood,'\n')
#se`lected correctly
correct_CS_on_IW_2_units_likelihood <- length(which(logdiff2 > (2)))/nrow(logliks_IW)*100
cat('Correctly choosing IW using 2 units difference likelihood:',correct_CS_on_IW_2_units_likelihood,'\n')
#undecided
undecided_CS_on_IW_2_units_likelihood <- 100 - error_CS_on_IW_2_units_likelihood - correct_CS_on_IW_2_units_likelihood
cat('Undecided using 2 units difference likelihood:',undecided_CS_on_IW_2_units_likelihood,'\n')

cat(
100 * length(which(logdiff2 > (2)))/nrow(logliks_IW),
100 * sum(logliks_CS[,'loglik_diff'] < CSminusIWdata)/length(logliks_CS[,1]),
100 * sum(logliks_IW[,'loglik_diff2'] < CSminusIWdata)/length(logliks_IW[,1]),'\n'
)

### Analyses of 100 trees

boots_emp <- read.table(paste0(dir,'out_empirical_M1000.txt'), header = TRUE)
boots_emp <- boots_emp[order(boots_emp[,1]),1:15]
colnames(boots_emp) <- c('the_island','the_dataset','island_age','M','the_init','model','model_type','methode','R','lam_c','mu','K','gam','lam_a','loglik')

logliks_emp <- matrix(nrow = max(boots_emp$the_island),ncol = 22)
logliks_emp[,1] <- 1:max(boots_emp$the_island)

# Subsetting and removing duplicates
CS_to_emp <- subset(boots_emp,the_dataset == 'empdata' & model_type == 'CS')
CS_to_emp <- CS_to_emp[!duplicated(CS_to_emp$the_island),]
IW_to_emp <- subset(boots_emp,the_dataset == 'empdata' & model_type == 'IW')
IW_to_emp <- IW_to_emp[!duplicated(IW_to_emp$the_island),]

for (i in 1:max(boots_emp$the_island)){
  logliks_emp[i,2:7]  <- as.numeric(CS_to_emp[which(CS_to_emp[,'the_island'] == i)[1],c('lam_c','mu','K','gam','lam_a','loglik')])
  logliks_emp[i,8:13] <- as.numeric(IW_to_emp[which(IW_to_emp[,'the_island'] == i)[1],c('lam_c','mu','K','gam','lam_a','loglik')])
}

colz <- c('the_island','lam_c_CS','mu_CS','K_CS','gam_CS','lam_a_CS','loglik_CS','lam_c_IW','mu_IW',
          'K_IW','gam_IW','lam_a_IW','loglik_IW','best_model','loglik_diff','< -2','between2','> 2','AICweights1','AICweights2',
          'AIC_CS','AIC_IW')

colnames(logliks_emp) <- colz

for(i in 1:nrow(logliks_emp)){
  the_logliks <- c(logliks_emp[i,'loglik_CS'],logliks_emp[i,'loglik_IW'])
  logliks_emp[i,'loglik_diff'] <- logliks_emp[i,'loglik_CS'] - logliks_emp[i,'loglik_IW']
  logliks_emp[i,'best_model']  <- as.numeric(which(the_logliks == max(the_logliks)))

  if(logliks_emp[i,'loglik_diff'] < (-2)){
    logliks_emp[i,'< -2'] <- 1
    logliks_emp[i,'between2'] <- 0
    logliks_emp[i,'> 2'] <- 0
  }

  if(logliks_emp[i,'loglik_diff'] > -2 & logliks_emp[i,'loglik_diff'] < 2){
    logliks_emp[i,'< -2'] <- 0
    logliks_emp[i,'between2'] <- 1
    logliks_emp[i,'> 2'] <- 0
  }

  if(logliks_emp[i,'loglik_diff'] > 2){
    logliks_emp[i,'< -2'] <- 0
    logliks_emp[i,'between2'] <- 0
    logliks_emp[i,'> 2'] <- 1
  }

  AICweights <- ICweights(the_logliks,c(4,4),100,'AIC')
  AICss <- AIC(the_logliks,c(4,4))
  logliks_emp[i,'AICweights1'] <- AICweights[1]
  logliks_emp[i,'AICweights2'] <- AICweights[2]
  logliks_emp[i,'AIC_CS'] <- AICss[1]
  logliks_emp[i,'AIC_IW'] <- AICss[2]
}

####
### PLOT CS PARAMETER DISTRIBUTION

row_max_IW_LL <- which.max(logliks_emp[,'loglik_IW'])

arrow_bottom <- -100
cex_lab <- 1.5
cex_size <- 0.8

library(latex2exp)

pic(name = paste0(dir,'CS_parameters_for_100_empirical_trees'),pictype = pictype)
par(mfrow = c(2,2),cex = cex_size,cex.lab = cex_lab,mar = c(4,5,1,2))
lam_c_min <- 0.35
lam_c_max <- 0.46
dens_min <- 0
dens_max <- 15
hist(CS_to_emp$lam_c,
     freq = TRUE, breaks = c(seq(lam_c_min,lam_c_max,length.out = 25)),
     xlab = TeX('$\\lambda_0 ^ c$'),
     main = NULL,
     col  = 'cornsilk3',
     border = 'cornsilk3',
     ylim = c(dens_min, dens_max),
     xlim = c(lam_c_min,lam_c_max))
##
med <- median(CS_to_emp$lam_c, na.rm = TRUE)
obs <- as.numeric(parsCS[1])
arrows(med, dens_max, med, 0, col = 'black',length = 0,lwd = 1.5)
arrows(obs, dens_max/4, obs, 0, col = 'blue',length = 0.05,lwd = 1.5)
obs_max_IW_LL <- as.numeric(logliks_emp[row_max_IW_LL,'lam_c_CS'])
arrows(obs_max_IW_LL, dens_max/4, obs_max_IW_LL, 0, col = 'red',length = 0.05,lwd = 1.5)
box(bty = 'l')

mu_min <- 0.08
mu_max <- 0.12
dens_min <- 0
dens_max <- 15
hist(CS_to_emp$mu,
     freq = TRUE, breaks = c(seq(mu_min,mu_max,length.out = 25)),
     xlab = TeX('$\\mu$'),
     main = NULL,
     col = 'cornsilk3',
     border = 'cornsilk3',
     ylim = c(dens_min, dens_max),
     xlim = c(mu_min,mu_max))
med <- median(CS_to_emp$mu,na.rm = TRUE)
obs <- as.numeric(parsCS[2])
arrows(med, dens_max, med, 0, col = 'black',length = 0,lwd = 1.5)
arrows(obs, dens_max/4, obs, 0, col = 'blue',length = 0.05,lwd = 1.5)
obs_max_IW_LL <- as.numeric(logliks_emp[row_max_IW_LL,'mu_CS'])
arrows(obs_max_IW_LL, dens_max/4, obs_max_IW_LL, 0, col = 'red',length = 0.05,lwd = 1.5)
box(bty = 'l')

K_min <- 35
K_max <- 38
dens_min <- 0
dens_max <- 15
hist(CS_to_emp$K,
     freq = TRUE, breaks = c(seq(K_min,K_max,length.out = 25)),
     xlab = TeX('$\\textit{K}$'),
     main = NULL,
     col = 'cornsilk3',
     border  = 'cornsilk3',
     ylim = c(dens_min, dens_max),
     xlim = c(K_min,K_max))
med <- median(CS_to_emp$K,na.rm = T)
obs <- as.numeric(parsCS[3])
arrows(med, dens_max, med, 0, col = 'black',length = 0,lwd = 1.5)
arrows(obs, dens_max/4, obs, 0, col = 'blue',length = 0.05,lwd = 1.5)
obs_max_IW_LL <- as.numeric(logliks_emp[row_max_IW_LL,'K_CS'])
arrows(obs_max_IW_LL, dens_max/4, obs_max_IW_LL, 0, col = 'red',length = 0.05,lwd = 1.5)
box(bty = 'l')

gam_min <- 0.00020
gam_max <- 0.00025
dens_min <- 0
dens_max <- 25
hist(CS_to_emp$gam,
     freq = TRUE, breaks = c(seq(gam_min,gam_max,length.out = 25)),
     xlab = TeX('$\\gamma_0$'),
     main = NULL,
     col = 'cornsilk3',
     border = 'cornsilk3',
     ylim = c(dens_min, dens_max),
     xlim = c(gam_min,gam_max))
med <- median(CS_to_emp$gam,na.rm = TRUE)
obs <- as.numeric(parsCS[4])
arrows(med, dens_max, med, 0, col = 'black',length = 0,lwd = 1.5)
arrows(obs, dens_max/4, obs, 0, col = 'blue',length = 0.05,lwd = 1.5)
obs_max_IW_LL <- as.numeric(logliks_emp[row_max_IW_LL,'gam_CS'])
arrows(obs_max_IW_LL, dens_max/4, obs_max_IW_LL, 0, col = 'red',length = 0.05,lwd = 1.5)
box(bty = 'l')

dev.off()


### PLOT IW PARAMETER DISTRIBUTION

row_max_IW_LL <- which.max(logliks_emp[,'loglik_IW'])

arrow_bottom <- -100
cex_lab <- 1.5
cex_size <- 0.8

library(latex2exp)

pic(name = paste0(dir,'IW_Parameters_for_100_empirical_trees'),pictype = pictype)
par(mfrow = c(2,2),cex = cex_size,cex.lab = cex_lab,mar = c(4,5,1,2))
lam_c_min <- 0.30
lam_c_max <- 0.45
dens_min <- 0
dens_max <- 15
hist(IW_to_emp$lam_c,
     freq = TRUE, breaks = c(seq(lam_c_min,lam_c_max,length.out = 25)),
     xlab = TeX('$\\lambda_0 ^ c$'),
     main = NULL,
     col  = 'cornsilk3',
     border = 'cornsilk3',
     ylim = c(dens_min, dens_max),
     xlim = c(lam_c_min,lam_c_max))
med <- median(IW_to_emp$lam_c, na.rm = TRUE)
obs <- as.numeric(parsIW[1])
arrows(med, dens_max, med, 0, col = 'black',length = 0,lwd = 1.5)
arrows(obs, dens_max/4, obs, 0, col = 'blue',length = 0.05,lwd = 1.5)
obs_max_IW_LL <- as.numeric(logliks_emp[row_max_IW_LL,'lam_c_IW'])
arrows(obs_max_IW_LL, dens_max/4, obs_max_IW_LL, 0, col = 'red',length = 0.05,lwd = 1.5)
box(bty = 'l')

mu_min <- 0.1
mu_max <- 0.2
dens_min <- 0
dens_max <- 15
hist(IW_to_emp$mu,
     freq = TRUE, breaks = c(seq(mu_min,mu_max,length.out = 25)),
     xlab = TeX('$\\mu$'),
     main = NULL,
     col = 'cornsilk3',
     border = 'cornsilk3',
     ylim = c(dens_min, dens_max),
     xlim = c(mu_min,mu_max))
med <- median(IW_to_emp$mu,na.rm = TRUE)
obs <- as.numeric(parsIW[2])
arrows(med, dens_max, med, 0, col = 'black',length = 0,lwd = 1.5)
arrows(obs, dens_max/4, obs, 0, col = 'blue',length = 0.05,lwd = 1.5)
obs_max_IW_LL <- as.numeric(logliks_emp[row_max_IW_LL,'mu_IW'])
arrows(obs_max_IW_LL, dens_max/4, obs_max_IW_LL, 0, col = 'red',length = 0.05,lwd = 1.5)
box(bty = 'l')

K_min <- 125
K_max <- 140
dens_min <- 0
dens_max <- 15
hist(IW_to_emp$K,
     freq = TRUE, breaks = c(seq(K_min,K_max,length.out = 25)),
     xlab = TeX('$\\textit{K}$'),
     main = NULL,
     col = 'cornsilk3',
     border  = 'cornsilk3',
     ylim = c(dens_min, dens_max),
     xlim = c(K_min,K_max))
med <- median(IW_to_emp$K,na.rm = T)
obs <- as.numeric(parsIW[3])
arrows(med, dens_max, med, 0, col = 'black',length = 0,lwd = 1.5)
arrows(obs, dens_max/4, obs, 0, col = 'blue',length = 0.05,lwd = 1.5)
obs_max_IW_LL <- as.numeric(logliks_emp[row_max_IW_LL,'K_IW'])
arrows(obs_max_IW_LL, dens_max/4, obs_max_IW_LL, 0, col = 'red',length = 0.05,lwd = 1.5)
box(bty = 'l')

gam_min <- 0.00025
gam_max <- 0.00040
dens_min <- 0
dens_max <- 15
hist(IW_to_emp$gam,
     freq = TRUE, breaks = c(seq(gam_min,gam_max,length.out = 25)),
     xlab = TeX('$\\gamma_0$'),
     main = NULL,
     col = 'cornsilk3',
     border = 'cornsilk3',
     ylim = c(dens_min, dens_max),
     xlim = c(gam_min,gam_max))
med <- median(IW_to_emp$gam,na.rm = TRUE)
obs <- as.numeric(parsIW[4])
arrows(med, dens_max, med, 0, col = 'black',length = 0,lwd = 1.5)
arrows(obs, dens_max/4, obs, 0, col = 'blue',length = 0.05,lwd = 1.5)
obs_max_IW_LL <- as.numeric(logliks_emp[row_max_IW_LL,'gam_IW'])
arrows(obs_max_IW_LL, dens_max/4, obs_max_IW_LL, 0, col = 'red',length = 0.05,lwd = 1.5)
box(bty = 'l')

dev.off()
```
