## Supplementary material for "The limits to ecological limits to diversification": Figures Tables and Supplementary Data and Scripts: Plot_Goodness_fit_frogs.R

```
summarize_bootstrap_results <- function(simulation_dataset, empirical_summary_statistics, m_pool = 1000, time = 30)
{
  load(simulation_dataset)
  replicates <- length(frogs_sims)

  ###### Calculate overall species richness and colonization statistics across all islands
  number_colonists <- c()
  number_spec <- c()
  number_stac0 <- c()
  number_stac1 <- c()
  number_stac2 <- c()
  number_stac3 <- c()
  number_stac4 <- c()
  number_stac2_singletons <- c()
  size_largest_clade <- c()
  size_smallest_clade <- c()
  rank_largest_clade <- c()

  for(i in 1:replicates) {
    the_island <- frogs_sims[[i]]
    number_colonists <- append(number_colonists, m_pool - the_island[[1]]$not_present)
    number_stac0 <- append(number_stac0, the_island[[1]]$not_present)
    unlist(the_island) -> u_island

    if(the_island[[1]]$not_present == m_pool) {
      number_spec <- append(number_spec, 0)
      number_stac1 <- append(number_stac1, 0)
      number_stac2 <- append(number_stac2, 0)
      number_stac3 <- append(number_stac3, 0)
      number_stac4 <- append(number_stac4, 0)
      number_stac2_singletons <- append(number_stac2_singletons, 0)
      size_largest_clade <- append(size_largest_clade, NA)
      size_smallest_clade <- append(size_smallest_clade, NA)
    } else {
      #number_spec <- append(number_spec, sum(the_island[[1]]$stt_all[nrow(the_island[[1]]$stt_all), c(2, 3, 4)]))
      s <- NULL
      cc <- NULL
      for(j in 2:length(the_island))
      {
        if(is.null(the_island[[j]]$all_colonisations))
        {
          s <- c(s, length(the_island[[j]]$branching_times) - 1)
          cc <- c(cc, the_island[[j]]$branching_times[2])
        } else
        {
          for(k in 1:length(the_island[[j]]$all_colonisations))
          {
            s <- c(s, length(the_island[[j]]$all_colonisations[[k]]$event_times) - 1)
            cc <- c(cc, the_island[[j]]$all_colonisations[[k]]$event_times[2])
          }
        }
      }
      scc <- sort(cc, decreasing = TRUE)
      rank_largest_clade <- c(rank_largest_clade,which(scc == cc[which.max(s)]))
      number_spec <- append(number_spec,sum(s))
      number_stac1 <- append(number_stac1, length(which(u_island[which(names(u_island) == "stac")] == 1)))
      number_stac2 <- append(number_stac2, length(which(u_island[which(names(u_island) == "stac")] == 2)))
      number_stac3 <- append(number_stac3, length(which(u_island[which(names(u_island) == "stac")] == 3)))
      number_stac4 <- append(number_stac4, length(which(u_island[which(names(u_island) == "stac")] == 4)))
      if(length(which(u_island[which(names(u_island) == "stac")] == 2)) > 0)
      {
        store_stac2s_brts <- list()
        store_all_brts <- list()
        for (isl in 2:length(the_island))
        {
          if(the_island[[isl]]$stac == 2) {store_stac2s_brts[[length(store_stac2s_brts) + 1]] <- the_island[[isl]]$branching_times}
          store_all_brts[[length(store_all_brts) + 1]] <- the_island[[isl]]$branching_times
        }
        number_stac2_singletons <- append(number_stac2_singletons, length(which(unlist(lapply(store_stac2s_brts, length)) == 2)))
        #size_largest_clade <- append(size_largest_clade, max(unlist(lapply(store_stac2s_brts, length))) - 1)
        #size_smallest_clade <- append(size_smallest_clade, min(unlist(lapply(store_all_brts, length))) - 1)
        size_largest_clade <- append(size_largest_clade, max(s))
        size_smallest_clade <- append(size_smallest_clade, min(s))
      } else {
        number_stac2_singletons <- append(number_stac2_singletons, 0)
        size_largest_clade <- append(size_largest_clade, NA)
        size_smallest_clade <- append(size_smallest_clade, NA)
      }
    }
  }

  the_prop <- c()
  max_clade <- c()
  the_prop_youngest <- c()

  frogs5 <- list()
  for(i in 1:length(frogs_sims)) {
    if(length(frogs_sims[[i]]) == 6){
      pl <- length(frogs5) + 1
      frogs5[[pl]] <- frogs_sims[[i]] }
  }

  for(i in 1:length(frogs5)) {
    island <- frogs5[[i]]
    if(length(island) == 1) {
      the_prop <- append(the_prop, 0)
      the_prop_youngest <- append(the_prop_youngest, 0)
    } else {
      island[[1]] <- NULL
      stac_vec <- unlist(island)[which(names(unlist(island)) == "stac")]
      stac_age_known <- which(stac_vec == 2 | stac_vec == 3 |  stac_vec == 4)
      btimes <- list()
      colonisation_time <- c()
      diversity <- c()
      for (o in 1:length(stac_age_known)) {
        btimes[[o]] <- island[[stac_age_known[o]]]$branching_times[-1]
        colonisation_time[o] <- max(btimes[[o]])
        diversity[o] <- length(btimes[[o]]) + island[[stac_age_known[o]]]$missing_species
      }

      seque <- cbind(colonisation_time, diversity)
      seque <- seque[rev(order(seque[, 1])), ]

      if(is.matrix(seque)){
        the_prop <- append(the_prop, as.numeric(seque[1, 2]/sum(seque[, 2])))
        the_prop_youngest <- append(the_prop_youngest, as.numeric(seque[nrow(seque), 2]/sum(seque[, 2])))
      }

      if(is.numeric(seque)){
        the_prop <- append(the_prop, 1)
        the_prop_youngest <- append(the_prop_youngest, 1)
      }
    }
  }

  tt <- matrix(ncol = 2, nrow = 5)
  tt[1:5,1] <- 1:5
  tt[1:5,2] <- hist(x = rank_largest_clade,
                    breaks = seq(0.5,max(rank_largest_clade) + 0.5, by = 1),
                    plot = FALSE)$density[1:5]

  overall_results <- list(number_colonists = number_colonists,
                          number_spec = number_spec,
                          number_stac0 = number_stac0,
                          number_stac1 = number_stac1,
                          number_stac2 = number_stac2,
                          number_stac3 = number_stac3,
                          number_stac4 = number_stac4,
                          number_stac2_singletons = number_stac2_singletons,
                          size_largest_clade = size_largest_clade,
                          size_smallest_clade = size_smallest_clade,
                          tt = tt)

  return(overall_results = overall_results)
}

plot_bootstrap_results <- function(overall_results, sumstats = c(65,5,28,1), ylims = c(0.01, 0.2, 0.15, 0.7), title = NULL)
{
  hist(x = overall_results$number_spec,
       breaks = 30,
       freq = FALSE,
       col = "cornsilk3",
       border = "cornsilk3",
       main = title,
       xlab = "Number of species",
       ylim = c(0,ylims[1]))
  med <- median(overall_results$number_spec, na.rm = T)
  arrows(med, ylims[1], med, 0, col = "black", length = 0, lwd = 1.5)
  arrows(sumstats[1], ylims[1]/4, sumstats[1], 0, col = 'blue', length = 0.07, lwd = 1.5)
  box(bty = 'l')

  ### Colonizations
  hist(x = overall_results$number_colonists,
       breaks = 10,
       freq = FALSE,
       col = "cornsilk3",
       border = "cornsilk3",
       main = NULL,
       xlab = "Number of colonization events",
       ylim = c(0,ylims[2]))
  med <- median(overall_results$number_colonists, na.rm = T)
  arrows(med, ylims[2], med, 0, col = "black", length = 0, lwd = 1.5)
  arrows(sumstats[2], ylims[2]/4, sumstats[2], 0, col = 'blue', length = 0.07, lwd = 1.5)
  box(bty = 'l')

  ##### size largest clade
  hist(x = overall_results$size_largest_clade,
       breaks = 30,
       freq = FALSE,
       col = "cornsilk3",
       border = "cornsilk3",
       main = NULL,
       xlab = "Size largest clade",
       ylim = c(0,ylims[3]))
  med <- median(overall_results$size_largest_clade, na.rm = T)
  arrows(med, ylims[3], med, 0, col = "black", length = 0, lwd = 1.5)
  arrows(sumstats[3], ylims[3]/4, sumstats[3], 0, col = 'blue', length = 0.07, lwd = 1.5)
  box(bty = 'l')

  bar_colors <- rep('cornsilk3',5)
  bar_colors[sumstats[4]] <- 'blue'
  barplot(height = overall_results$tt[, 2],
          names.arg = c('1st', '2nd', '3rd', '4th', '5th'),
          col = bar_colors,
          border = bar_colors,
          bty = "L",
          xlab = 'Rank largest clade',
          ylab = 'Density',
          ylim = c(0, ylims[4]))
  box(bty = 'l')
}

get_stats_cppfun <- Rcpp::cppFunction('
NumericMatrix get_stats_cpp(const List& brts_list,
                            const NumericVector& tt,
                            const float alpha) {
  NumericMatrix output(tt.size(), 5);

  size_t min_index = brts_list.size() * alpha/2;
  size_t max_index = brts_list.size() * (1 - alpha/2);

  Rcout << "0--------25--------50--------75--------100" << std::endl;
  Rcout << "*";
  int updateFreq = tt.size() /  20;
  if(updateFreq < 1) updateFreq = 1;

  for (int i = 0; i < tt.size(); ++i) {
    if (i % updateFreq == 0) {
      Rcout << "**";
    }


    float focal_t = tt[i];
    std::vector<size_t> found_lin(brts_list.size());
    float mean = 0.f;
    NumericVector focal_brts;
    for (int j = 0; j < brts_list.size(); ++j) {
      focal_brts = brts_list[j]; // temp
      size_t cnt = 0;
      while(focal_brts[cnt] > focal_t) {
        cnt++;
      }
      found_lin[j] = cnt; // two at first timepoint, crown age.
      mean += found_lin[j];
    }
    mean *= 1.0 / brts_list.size();

    std::sort(found_lin.begin(), found_lin.end());
    float median = found_lin[ found_lin.size() / 2 + 1 ];

    if (found_lin.size() % 2 == 0) {
      median = 0.5 * (found_lin[ found_lin.size() / 2] +
                      found_lin[ found_lin.size() / 2 + 1]);
    }

    float minalpha = found_lin.at( min_index );
    float maxalpha = found_lin.at( max_index );

    NumericVector add(5);
    add(0) = focal_t;
    add(1) = median;
    add(2) = minalpha;
    add(3) = maxalpha;
    add(4) = mean;

    output(i, _) = add;
  }
  return output;
}')

prepare_plot_alpha_base_highres <- function(sim_dataset,
                                            emp_dataset,
                                            alpha = 0.05) {

  load(sim_dataset)

  cat("converting simulation output to branching times: \n")
  pb <- txtProgressBar(style = 3, max = length(frogs_sims))
  brts_list <- c()
  for (i in 1:length(frogs_sims)) {
    temp_brt_table <- DAISIE:::add_brt_table(frogs_sims[[i]])
    brts_list[[i]] <- temp_brt_table[[1]]$brts_table[, 1]
    setTxtProgressBar(pb, i)
  }

  max_t <- sapply(brts_list, function(x) max(x))
  max_t <- max(max_t)

  cat("\ncollecting unique branching times: \n")
  pb <- txtProgressBar(style = 3, max = length(brts_list))

  all_t <- c()
  for (i in 1:length(brts_list)) {
    all_t <- c(all_t, brts_list[[i]])
    setTxtProgressBar(pb, i)
  }
  all_t <- unique(all_t)
  all_t <- sort(all_t, decreasing = TRUE)

  cat("\ncalculating CI for", length(all_t), "timepoints: \n")

  sim_plot <- get_stats_cppfun(brts_list, all_t, alpha)

  cat("\n")

  colnames(sim_plot) <- c("t", "median", "minalpha", "maxalpha", "mean")

  load(emp_dataset)

  frogs <- DAISIE:::add_brt_table(frogs[[52]])
  x <- frogs[[1]]$brts_table[,1]
  y <- 1:length(x)
  emp_plot <- cbind(x, y)

  return(list(sim_plot = sim_plot, emp_plot = emp_plot))
}

plot_alpha_base_highres <- function(sim_plot,
                                    emp_plot,
                                    xlabel = "Time (Mya)",
                                    ylabel = "1 + # Lineages",
                                    plot_title = "CW",
                                    area_fill = 'cornsilk3',
                                    min_y = 2,
                                    max_y = max(sim_plot[, 4], emp_plot[, 2])) {

  plot(x = NA, y = NA,
       xlim = c(30, 0),
       ylim = c(min_y, max_y),
       log = "y",
       bty = "l",
       xlab = xlabel,
       ylab = ylabel,
       main = plot_title)

  polygon(x = c(sim_plot[, 1], rev(sim_plot[, 1])),
          y = c(sim_plot[, 4], rev(sim_plot[, 3])), col = area_fill, border = NA)

  lines(x = sim_plot[, 1], y = sim_plot[, 2], col = "black", type = "s", lwd = 2)
  lines(x = emp_plot[, 1], y = emp_plot[, 2], col = "blue", type = "s", lwd = 2)
}

library(Rcpp)
library(DAISIE)

# setwd("/Users/janzen/Dropbox/projects/ltt_rampal/")
setwd("d:/data/tex/Etienne et al 2019")

dataset_emp <- 'Frogs_100_30_miss_spec_added_M1000.Rdata'
dataset_CS <- 'frogs_sim_CS_T30_M1000_R5000.RData'
dataset_IW <- 'frogs_sim_IW_T30_M1000_R5000.RData'
overall_results_CS <- summarize_bootstrap_results(simulation_dataset = dataset_CS)
overall_results_IW <- summarize_bootstrap_results(simulation_dataset = dataset_IW)
pdf(file = 'd:/data/tex/Etienne et al 2019/Goodness_of_fit.pdf', width = 10, height = 5)
par(mfrow = c(2, 4), cex.lab = 1.5, cex.main = 1.5)
plot_bootstrap_results(overall_results = overall_results_CS,
                       title = 'Simulated under CS',
                       ylims = c(0.01, 0.2, 0.15, 0.7))
plot_bootstrap_results(overall_results = overall_results_IW,
                       title = 'Simulated under IW',
                       ylims = c(0.05, 0.25, 0.025, 0.7))
dev.off()

#pdf(file = 'd:/data/tex/Etienne et al 2019/Goodness_of_fit2.pdf', width = 10, height = 5)
#par(mfrow = c(1, 2), cex.lab = 1.5, cex.main = 1.5)
#DAISIE::DAISIE_plot_sims(island_replicates = frogs_sims, sample_freq = 25)
#DAISIE::DAISIE_plot_sims(island_replicates = frogs_sims, sample_freq = 25)

pdf(file = 'LTT.pdf', width = 10, height = 5)
par(mfrow = c(1, 2), cex.lab = 1.5, cex.main = 1.5)

plots_CS <- prepare_plot_alpha_base_highres(dataset_CS,
                                            dataset_emp)
plots_IW <- prepare_plot_alpha_base_highres(dataset_IW,
                                            dataset_emp)
plot_alpha_base_highres(plots_CS[[1]],
                        plots_CS[[2]],
                        plot_title = "CS",
                        min_y = 1,
                        max_y = 150)
plot_alpha_base_highres(plots_IW[[1]],
                        plots_IW[[2]],
                        plot_title = "IW",
                        min_y = 1,
                        max_y = 150)
dev.off()
```
