## Supplementary material for "The limits to ecological limits to diversification": Figures Tables and Supplementary Data and Scripts: Response letter Etienne et al.pdf

*Dear Editor,*

*We are pleased to submit our revision of the manuscript "The limits to ecological limits to diversification" which was submitted previously under ms ID USYB-2019-266. Because of COVID-19 and because we did a major overhaul of our code (see below), this revision took a lot longer than we had anticipated. But we hope it is in much better shape now. Below we provide a point-by-point response to the comments made by editors and reviewers. We hope it is now acceptable for publication in Systematic Biology.*

*Kind regards,, on behalf of all authors,*

*Rampal Etienne*

**Dr Bryan Carstens**
**Editor in Chief, Systematic Biology**

Thank you for your Systematic Biology submission. It has been reviewed by Associate Editor Professor Tanja Stadler, two expert reviewers (below), and myself. Our general consensus is that this work is potentially important and likely represents a useful advance. However, none of us believe that the submission should be published in its current form. In addition to the useful comments from the reviewers, particularly reviewer 2, I have a couple of concerns about the manuscript. First of all, it is too understated in its description of how this work represents an important advance to the field. As a result, I am not completely sold on the importance of this work to the field, although I suspect that I could be convinced. Consider adding a conclusion that makes a succinct and direct argument as to why this work matters, and as to how it will improve our understanding of island biogeography. Secondly, the readability of the manuscript could be improved, particularly in the introduction and the section of the methods that concerns the derivation of the equation. You might consider tightening this section up to reduce the overall manuscript length, and provide a complete derivation in supplemental materials. Finally, the Abstract is overly long and reads more as an introduction. Please consider shortening this and leaving out some of the background details which are largely repeated in the Introduction.

*Thank you for your comments. To answer your specific questions: we have shortened the abstract - indeed it read a bit as an introduction - and added a conclusion on why this work is important in the abstract and in the discussion. We have moved the equations to the supplementary material so we have more space to explain them.*

**Associate Editor: Professor Tanja Stadler**

Thank you for submitting your manuscript to Syst. Biol.

I received two independent reviewer reports. Overall, they rate your manuscript as very interesting, and I agree with them.

*Thank you for these kind words.*

Several severe concerns are raised and I again agree with them. Please carefully consider the reviewer comments. In particular:

"What is a little unclear to me is what to use this for. The methodology is clearly described, but not to a degree where other researchers are likely to use this model more broadly. Publishing it as an R package, like daisie, would open up this. As an empirical paper the single test example is a bit insufficient. Perhaps reimplementing the model in a faster language could make it possible to extend this to a broader example."

*The methodology IS available in DAISIE. We have made this clearer in the revision. In addition, to make the methodology more user-friendly, we have now written a step-by-step tutorial, which we provide as an Rmarkdown vignette/demo in the new version of the R package DAISIE. We attach a pdf version of this demo to the submission for easy access to the referees.*
*We reimplemented the computationally demanding parts of the model in C++ and now it runs a lot faster and can apply to more data sets. Regarding other empirical examples: our empirical example serves as an illustration of our method, and thus we show to the empiricists how they could apply our method to their dataset.*

With respect to applications, it seems that your IW model could also be adopted to be used for local epidemic outbreaks (hospitals, schools). This may serve as an additional empirical example, or could be mentioned in the discussion.

*This is an interesting suggestion that we now include in the discussion. As an empirical example it is less appropriate for this paper, because this paper is about the phylogenetic limits to ecological limits to diversification, whereas in the epidemic example it is clear that the IW model is appropriate because infections with multiple sources will all compete for susceptible hosts. But again, this is a really good example of another application of the IW model (see also our response to one of the reviewer comments below).*

Further, please make more effort to convince us that the implementation is correct! We appreciate the difficulty given the slow runtime, however, proceeding with code without a very good validation is problematic. I want to avoid later controversies, as we see with other phylogenetic methods which were not validated initially.

*We agree that correctness of the implementation is of utmost importance. We have checked that in the case of infinite carrying capacity the CS and IW model give the same result, and checked the code over and over again. Also, the fact that the generating parameter set falls well within the distribution of the estimated parameters provides more confidence in the implementation. Still, a bug is always possible, and detection is difficult because this is a new model for which no independent check exists.*

**Reviewer: 1**

This paper presents an extension to the well-established island biogeography "daisie" model, which allows it to extend diversity-dependence from within the evolving clade to all species (of a given taxon) on the island. They call the latter "island-wide", though it's more an arbitrary phylogenetic scale larger than the individual radiation. They test the model for a genus of frogs on Hispaniola, and find that the "island-wide" extension fits the data more poorly than the original per-clade model.

It's an interesting question at what phylogenetic scale diversity-dependence should act, and this model seems like it could give some indications on this. Intuitively I'd think that the strength of diversity-dependence would decrease more as a continuous function of phylogenetic distance, though I will agree with the authors that "clade-wise" and "island-wise" seem to be both often implicitly accepted scales.

*We agree that the strength of diversity-dependence is likely to decrease continuously with phylogenetic distance. This is, however, very difficult to implement in a likelihood framework. We believe that the two scales we consider here are a good first start. Species are either closely related, because they originate from the same immigrant, or they are distantly related because they originate from different immigrants which can be expected to be phylogenetically far apart compared to cladogenetic species on the island.*

The model itself looks correctly performed, though I should say I don't feel qualified to review the complex collection of equations here. The statistical modelling and analysis are also done with care and competence.

*Thank you for these kind words.*

What is a little unclear to me is what to use this for. The methodology is clearly described, but not to a degree where other researchers are likely to use this model more broadly. Publishing it as an R package, like daisie, would open up this. As an empirical paper the single test example is a bit insufficient. Perhaps reimplementing the model in a faster language could make it possible to extend this to a broader example.

*See our response to the Associate editor.*

**Reviewer: 2**

The authors compare a clade-specific diversity dependence model to an island-wide model that allows genetically distant species compete with each other for niches. The main challenge is calculating the likelihood of the island-wide model, which is expensive both in computing time and memory usage. Because of the cost of the model the implementation is not exhaustively validated here, although it appears to be work well enough on this dataset. I particularly like the inclusion of a goodness-of-fit test and moving away from using only AIC/BIC statistics to perform model selection. I think the methods section could be improved and I have a few issues with the empirical analysis.

*Thank you for these encouraging words. We have improved on the methods section by incorporating the comments of the reviewer.*

1.) I found the derivation of the IW model likelihood to be quite painful. This is mainly because of the number of different events that have to be kept track of. I think it would be easier to follow if the authors first properly define P and Q and then carefully label what event each term in the equations represents. Some terms in the equations and in the subscripts can also be simplified (e.g. M - 1 - m + 1). If they are intentionally left unsimplified it should be explained why.

*We wholeheartedly agree that the derivation IS indeed very painful! We have now moved the derivation to the supplementary material so we have more space to explain it in more detail. We simplified our notation where possible.*

I also found the Q superscripts to be confusing. The way I understand it is that the value of the superscript (1,2,3 etc.) denotes the order of the colonizing species (3 would be for the 3rd mainland species to colonise the island) and the ± indicates whether this species has speciated after colonisation (+) or not (-). In this way the likelihood keeps track of the exact order of colonization and speciation within the l-vector. Is this really necessary? Is there a reason why you can't write down the likelihood while only keeping track of m, e, l^{+} and l^{-}? That would simplify things and avoid needing the l_i vector/operator (which I also found confusing).

*The reviewer has understood the notation correctly. It is not necessary to keep track of the exact order of colonization, but it is necessary to keep track of which species has speciated and which has not after colonization, so the identity of the clade matters. This is because the data tells us the endemicity status of the clade and the colonization time of the clade. An earlier colonization is more likely to undergo speciation during its (longer) presence on the island, and hence the probability of endemicity is higher. When only keeping track of l^{+} and l^{-}, one loses this connection between colonization time and endemicity status. We have explained this in the revision.*
*We have, however, completely rewritten the likelihood derivation and moved it to the supplementary material to allow for a more elaborate explanation. We hope it is much clearer now.*

2.) I am a little sceptical about the method of randomly inserting missing species into subclades. I assume this was done because no genomic data are available for these species? I don't have access to Table S1 and cannot see the rationale for the placements of these species and how uncertain the placements are. Were the results any different if those 8 species were left out completely?

*Indeed there is no genetic data available on GenBank for these eight species (they have been recently described, are known from a single specimen or are possibly extinct). In table S1 we explain in detail the rationale for the placements with all the appropriate references, but we summarize it here, and we have now added this information to the main text. Four of the missing species were previously considered subspecies, and have recently been elevated to species, and we thus randomly inserted them (100 times) to a location along the tip branch of the species they were previously assigned to. Four species have been proposed to belong to well-defined terminal clades based on morphology, and we randomly inserted them at any position within those clades (100 times). The results are robust across all these 100 datasets.as we explained in the ms. The remaining species is* E. neiba*, for which no data is available for a topological assignment, and we already explain in the main text how we dealt with it (including it or excluding it does not affect the results).*
*Regarding the question of whether leaving those 8 species out completely alters the results, we have run an analysis excluding them. We find that the same model is preferred, and the parameters vary only slightly.*

Is there a reason why the IW and CS models can't be extended to include partial sampling (as was done in Etienne et al. 2012)? Unless I'm mistaken, partial sampling only affects the last step of the likelihood computation, so it should not introduce a large cost and since the proportion of sampled species is known it would not introduce any new parameters. This would be a better way of accounting for missing species than to randomly insert them into the tree.

*In the CS model this can indeed be done and indeed only concerns the last step of the likelihood computation where for every clade one can take the (n + 1)-th component of the Q-vector when there are n unsampled species (see Valente et al. 2015 where we explain this approach in the first implementation of CS in DAISIE). In the IW model, the index that corresponds to the missing endemic species is e, which keeps track of the missing number of endemics in ALL the clades, including clades for which the colonization times are unknown (if we had to keep track of the number of endemics for each colonizing clade, the dimensionality of the problem would explode). Thus we could use this to specify unsampled species if we assumed that the unsampled species could have originated anywhere. However, we did not do this because we know with high certainty to what exact clade the unsampled species belong; we just don't know their branching times. We believe that this additional information is important to use, and we accounted for the uncertainty in the branching time of these missing species by considering various possibilities of where the missing species branched off. We explained this better in the revision.*

3.) It is well-known that raw likelihood values are not a good way of comparing models with different levels of complexity. Since several of the models used here have the same number of parameters, the AIC and BIC just amounts to comparing likelihood values. However, there is still a difference in model complexity (and degrees of freedom) between, for example, the CS and IW models or the DI and CS without anagenesis models, even though they have the same number of parameters.

Therefore, I think the LRT from the bootstrapped datasets is the best way to proceed. However, only the BIC is used in the first model selection step (deciding between the DI and the two CS models to pick the forests with the best/worst likelihoods). In the second model selection step the DI model and the CS and IW models with anagenesis are all ignored. I agree that the models with anagenesis can probably be ignored, since it has already been shown previously that there is very little power in the model to detect anagenesis. Is there a good reason why the DI model was not used in the likelihood ratio test?

*We were interested in the comparison of the two types of diversity-dependence, CS and IW, because we are concerned with the question on the phylogenetic limits to ecological limits. Considering a model without ecological limits in detail is not so relevant in this light. Our ms is not so much about the question whether rain frogs experience CS or IW diversity-dependence, but to introduce the method and illustrate it with the rain frogs.*

4.) I think assuming the same carrying capacity for all clades in the CS model is a very strong assumption. This means assuming that K is at least 28 for all clades, which may not be the case for a clade consisting of species all competing for a very limited number of niches. I didn't see any exploratory analyses with LTT plots to see if there are indications that any of the clades are close to saturation. From eyeballing the trees it looks like clade 2 (21 species) is closer to saturation than clade 1 (28 species).

Would it be possible to try fitting a CS model with independent K's to the data? It would be interesting to see if such a model infers $K < 28$ for any of the clades and if it fits the data better. The likelihood computation would not be any more expensive or use significantly more memory. It might not be possible to identify K for clades 3-5, so they may need to be removed from the analysis. Since clades are assumed independent under the CS model and these clades do not contain many data points, the effect on speciation/extinction rate estimates should be small.

*We agree that assuming the same K across all clades in the CS model is a strong assumption. It is also technically possible to fit a model with independent K's for all of the clades. There are several reasons why we did not consider this: 1. We would need to assume a K for all possible colonists, including those that have no descendants on the island! That would be a very large number of parameters to add. 2. Removing data points because they have too few species will bias the results. 3. It is known that estimating K on single clades leads to biased estimates (Etienne et al. 2017); the merit of DAISIE with a single K value for all clades is that it reduces this bias. 4. In this ms we are proposing a new model (IW) that is compared to an existing model (CS); we did not want to change the existing model as well. 5. We*

*are considering elsewhere a model which assumes that K can be different for different clades by allowing K to be sampled from a distribution. This is, however, beyond the scope of this paper.*
*The associate editor mentioned an interesting application of our model, that of local spread of an epidemic, such as a virus in schools or hospitals. Interestingly, in these cases the number of children/patients represents the carrying capacity K. In the CS model this K is identical for different strains of the virus where the hosts can be infected multiple times, whereas in the IW model the K is the overall limit for all strains: once a host is infected, no other strain can infect the host, for example because the host builds up immunity. We discuss this example in the revision.*

5.) It looks like the figures/captions in Figure 4 and 5 are switched around. Figure 4 shows the results using the parameters estimated for the IW model and Figure 5 for the CS model. It would also be useful to add the results of fitting the CS model fitted to the 1600 datasets simulated under the IW model (and vice-versa) to the supplement.

*This is indeed a mistake. Thank you for noting this; we have corrected this. We have added the other results to the supplementary material as suggested.*

6.) Please number the equations.

*The equations have been numbered.*

7.) Could the colonisation process be seen as an ongoing process from the earliest appearance of a mainland species on the island until the first speciation event (until then any new colonisations by this species simply join the existing mainland population on the island)? Would this avoid having to prohibit any new colonisations by this species during this period?

*There are indeed two options: 1. Recolonization does nothing to the original population, i.e. it does not alter the colonization time. This is suggested by the reviewer. 2. Recolonizations are not allowed until speciation occurs because a selective sweep is expected that would reset the colonization time (assuming that mainland species are more fit than island species which can be argued from the success of invasive species on islands, and also theoretically from e.g. Rosindell et al. 2015. We chose the second option, which has been in the DAISIE model since the original paper on the CS model (Valente et al. 2015).*

8.) Line 231: N doesn't need to be a function of time here. In previous equations the subscript already denotes the total population size at a given time (e.g. m + e + 2 - 1). If N is a function of t, the subscripts in all previous equations can just become N, which would make the equations simpler to read, but in turn make it more difficult to figure out the effective colonisation/speciation rates at a glance.

*We have dropped the dependence on time as suggested.*

9.) Line 275: I don't understand the statement: "We did not want to use the maximum clade credibility tree, as this tree maximizes the credibility of each clade but does not necessarily result in a tree with high posterior probability." The maximum clade credibility tree was used as the starting point and was only modified by adding the missing species. At no point is the posterior probability of the 100 trees calculated and the MCC tree is still being used.

*Thank you for spotting this. Indeed this does not make much sense. We removed the confusing sentence..*

10.) When fitting the DAISIE models to the Hispaniola clades extracted from the BEAST MCC tree the models are fitted to a forest, not a tree, since the dataset contains 5 separate trees.

*The reviewer is right. We have changed our wording to "set of trees".*

11.) Where does the number of 300 mainland frog species come from? Earlier it is stated that 148 species comprise 89% of Caribbean diversity. Are there another 140 continental Eleutherodactylus frog species and is there reason to believe that continental species as well as Caribbean species are able to colonise Hispaniola?
*The value of M has very little effect on the parameters, except for colonization rate if M is large enough (which it is in this case). In fact, what we are estimating is M*gamma, so the total colonization rate, and what we see is that when we increase M, we decrease the estimated colonization rate. But in simulations the results depend largely on M*gamma and hence these are not changed. We changed the mainland pool size to 1000 which has become the standard in DAISIE analyses, and this confirms that very little changes.*

12.) Does the model assume that the number of mainland species stays constant through time? Is this realistic for the frog dataset?

*Yes, the model assumes that this number is constant. The mainland pool size may have grown in time for the frogs. If this were very important, we would have seen a poorer fit of the models to the data than what we observe in our goodness-of-fit plots.*

13.) On the model selection step on the 100 forests, did all 100 forests have the same best-fitting model?

*Yes, for all 100 forests the CS model (with no anagenesis) was the preferred model, with both AIC and BIC. We have added this information to the methods.*

14.) Would it be possible to get 95% CIs for parameter estimates? (perhaps using an LRT and a chi-squared approximation).

*The bootstrap results can be used for this. The frequency distribution of the parameter estimates provides an approximation of the uncertainty distribution. These histograms were already provided in the ms.*

15.) Out of the simulated replicates that could not be correctly assigned using the LRT, how many selected the wrong model and how many were indecisive?

*We added these numbers in the revised ms.*

16.) I don't think it is at all surprising that the first (oldest) clade is most often the largest under the CS model. The first clade has been in existence the longest (extinct clades are not observed). Since all clades have the same speciation/extinction rates and carrying capacities, the oldest clade is more likely to have more species. If different clades had different carrying capacities this would no longer be the case, since it would then be possible for two clades with the same number of species to have different effective speciation rates.

*We fully agree. We would have expected, however, that this priority effect is much more pronounced in the IW model. This does not seem to be the case. We have added some discussion on this point.*
