## Supplementary figures and images for "The limits to ecological limits to diversification"

### Bootstrap_CS_pars_under_IW.pdf

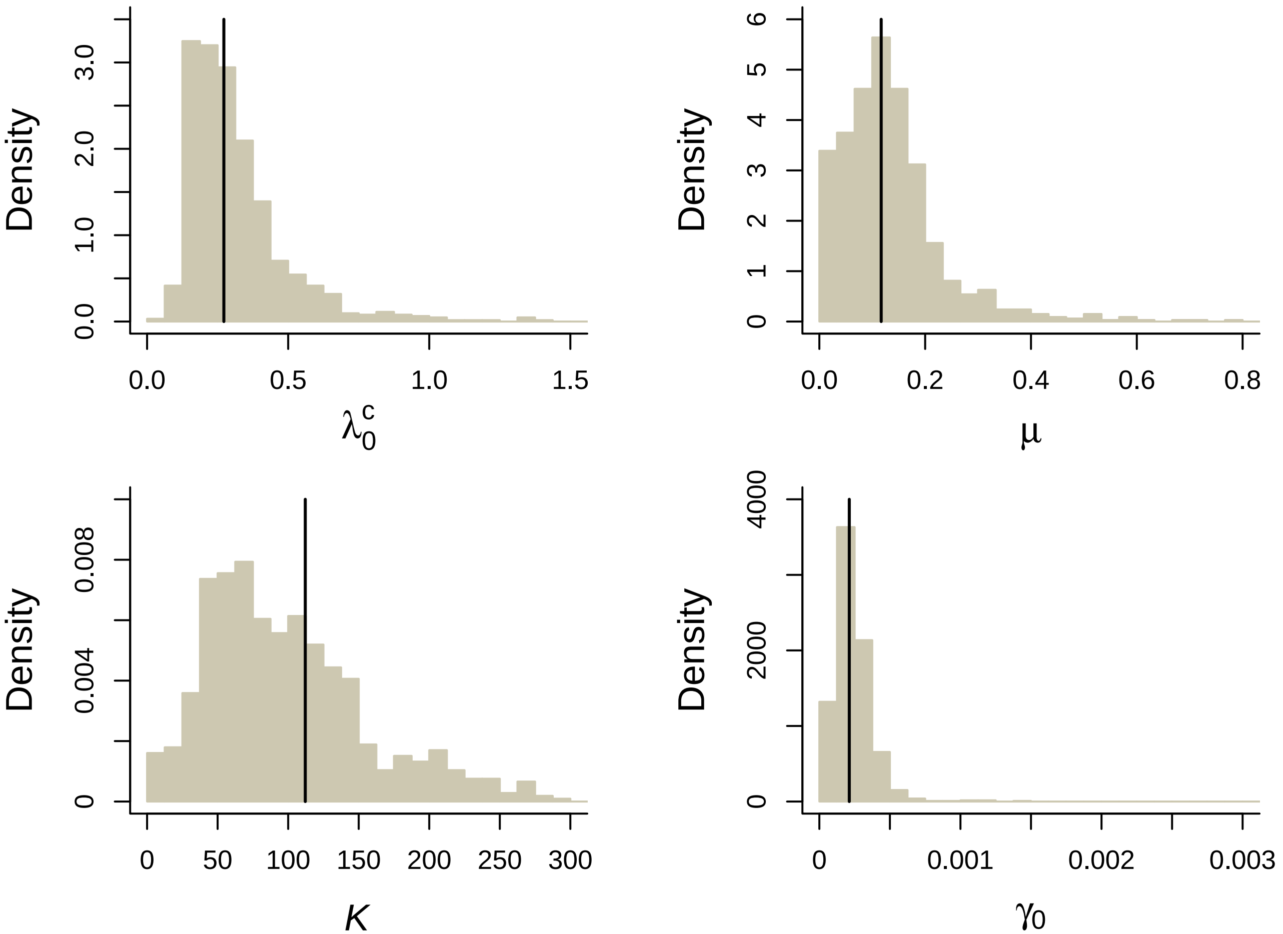

### Bootstrap_IW_pars_under_CS.pdf

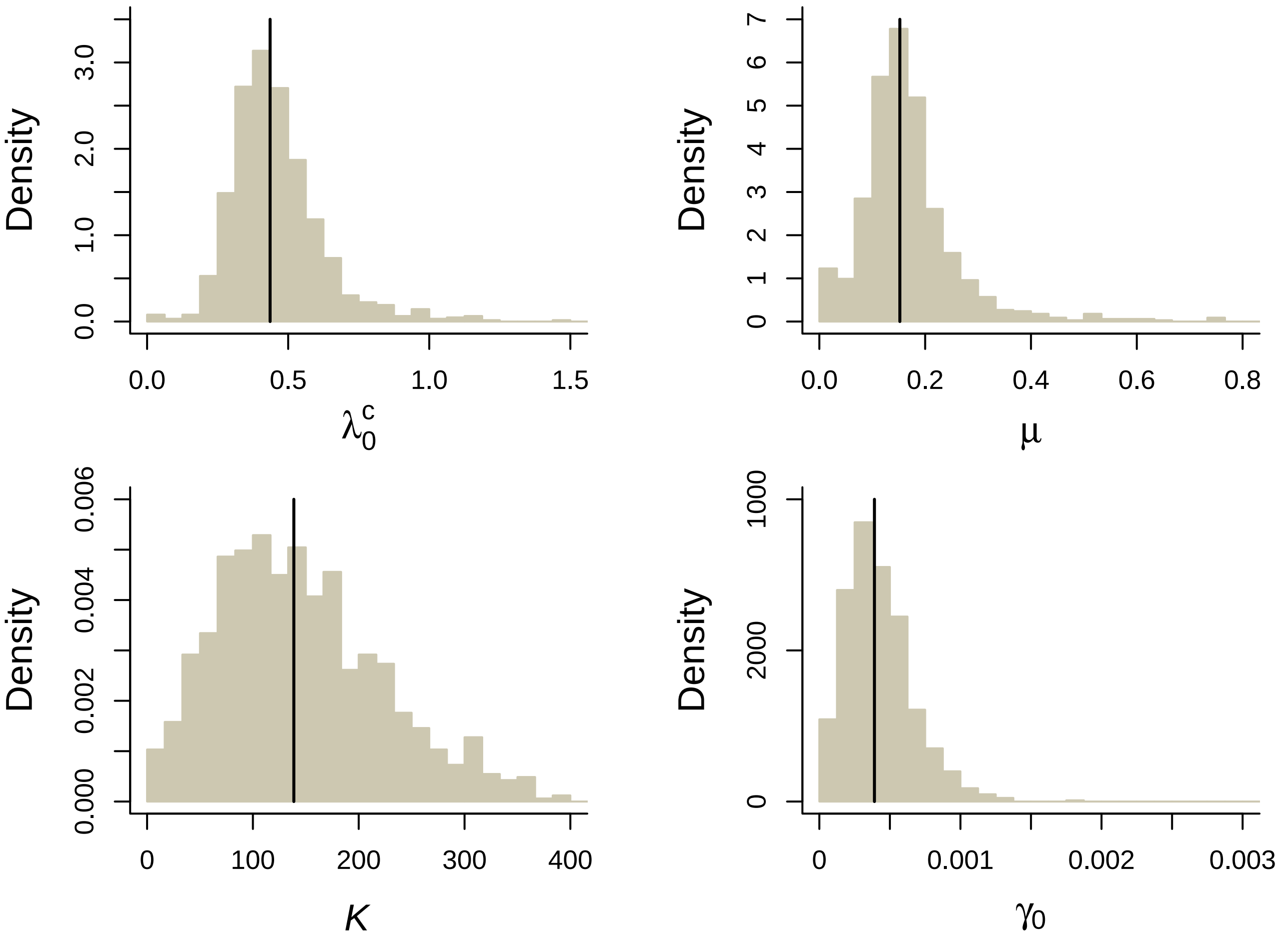

### Bootstrap_pars_under_CS.pdf

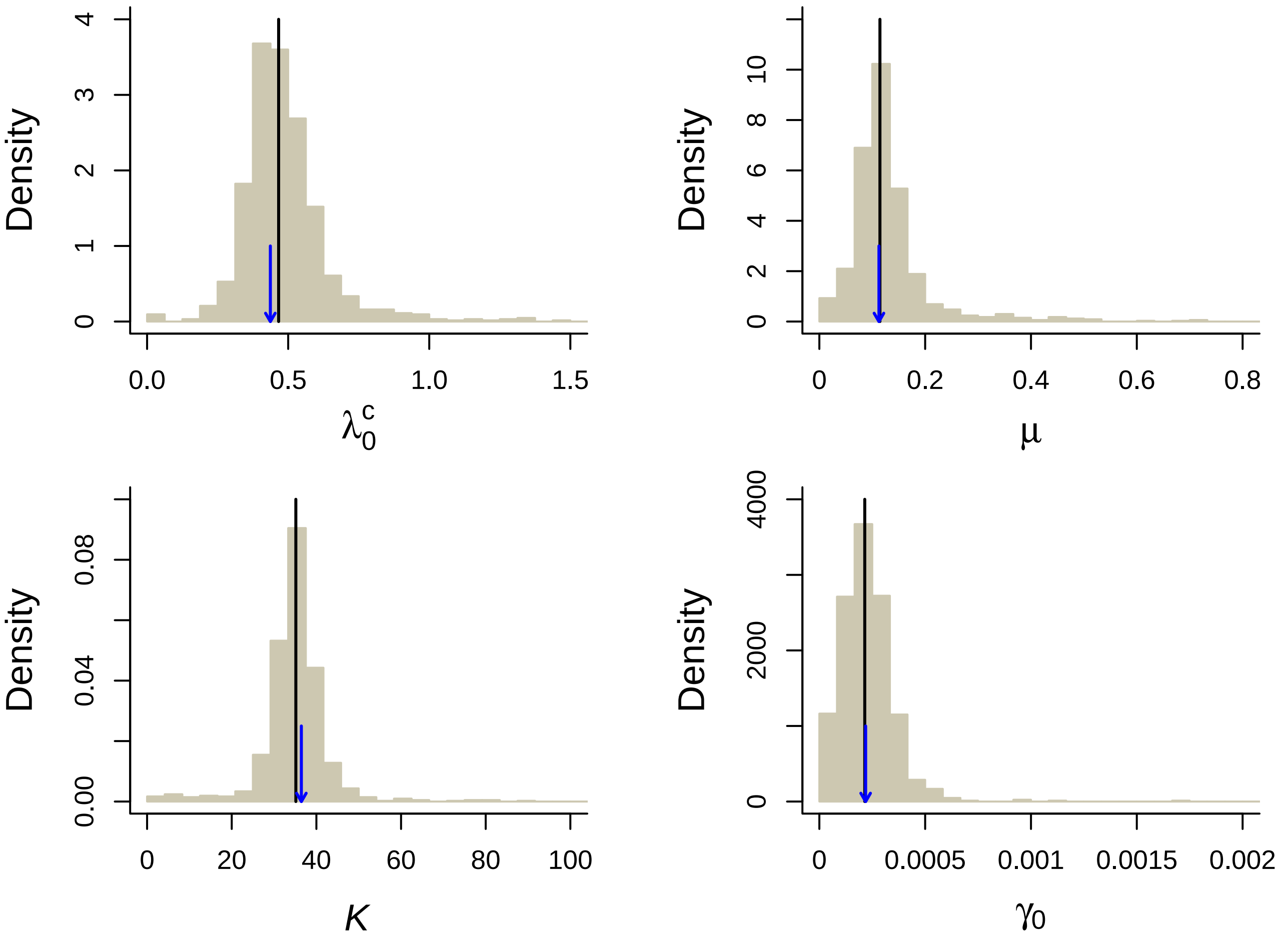

### Bootstrap_pars_under_IW.pdf

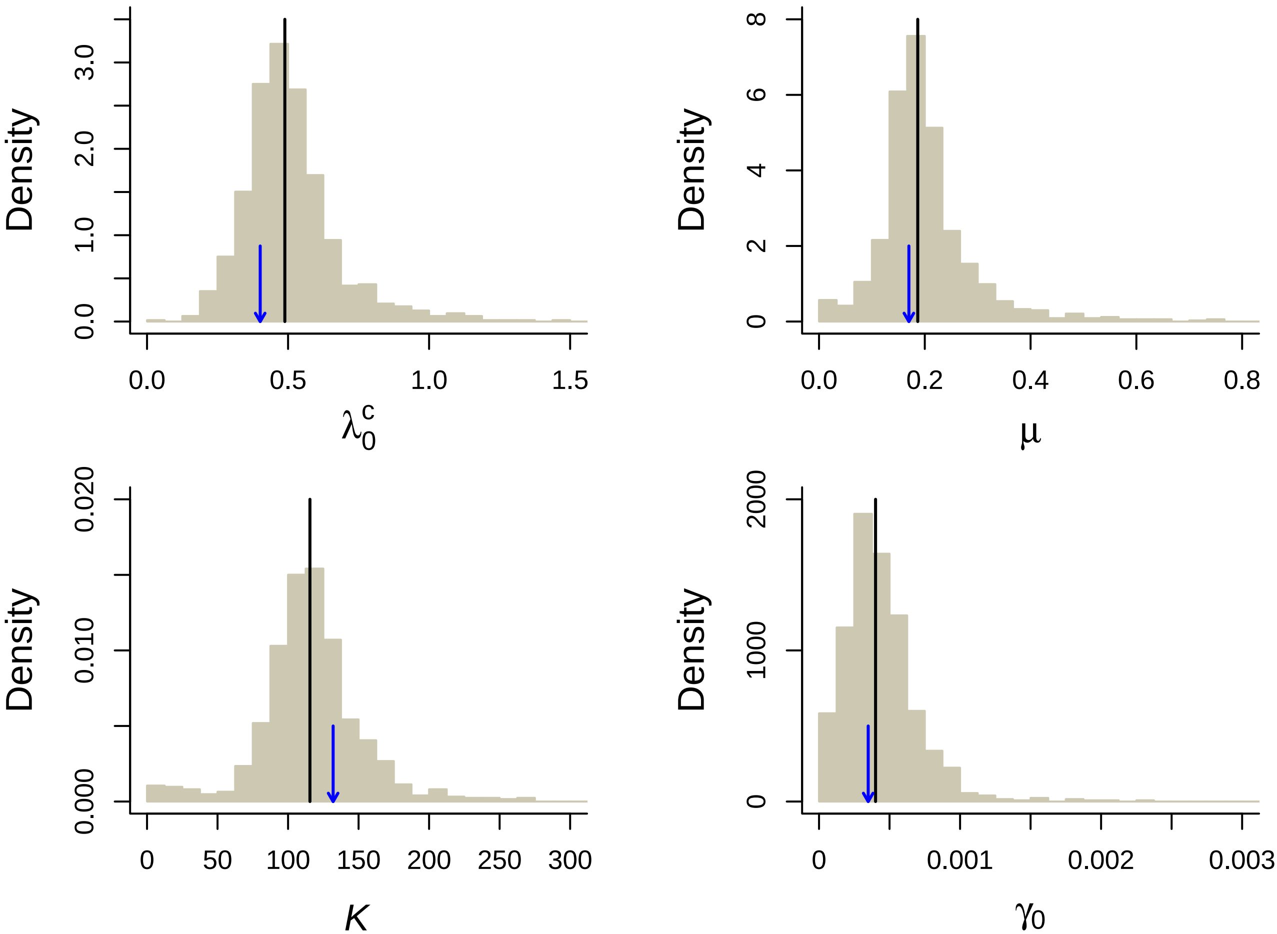

### diagram_likelihood.png

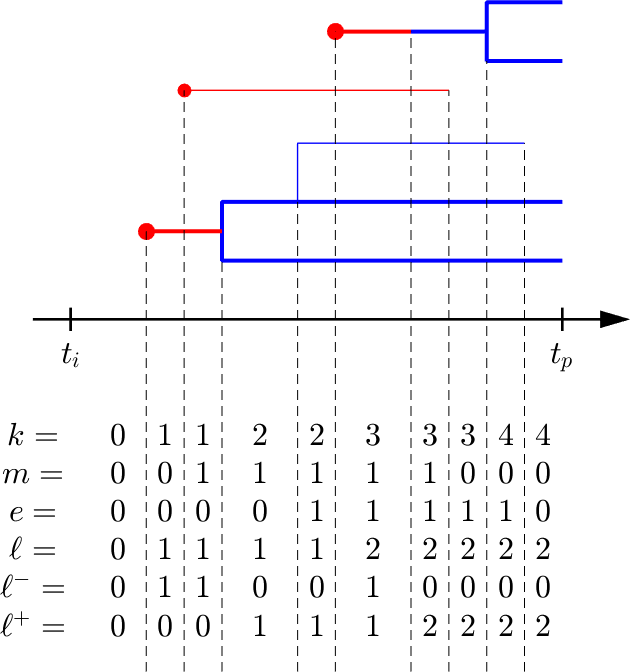

### Figure 2_Modified.png

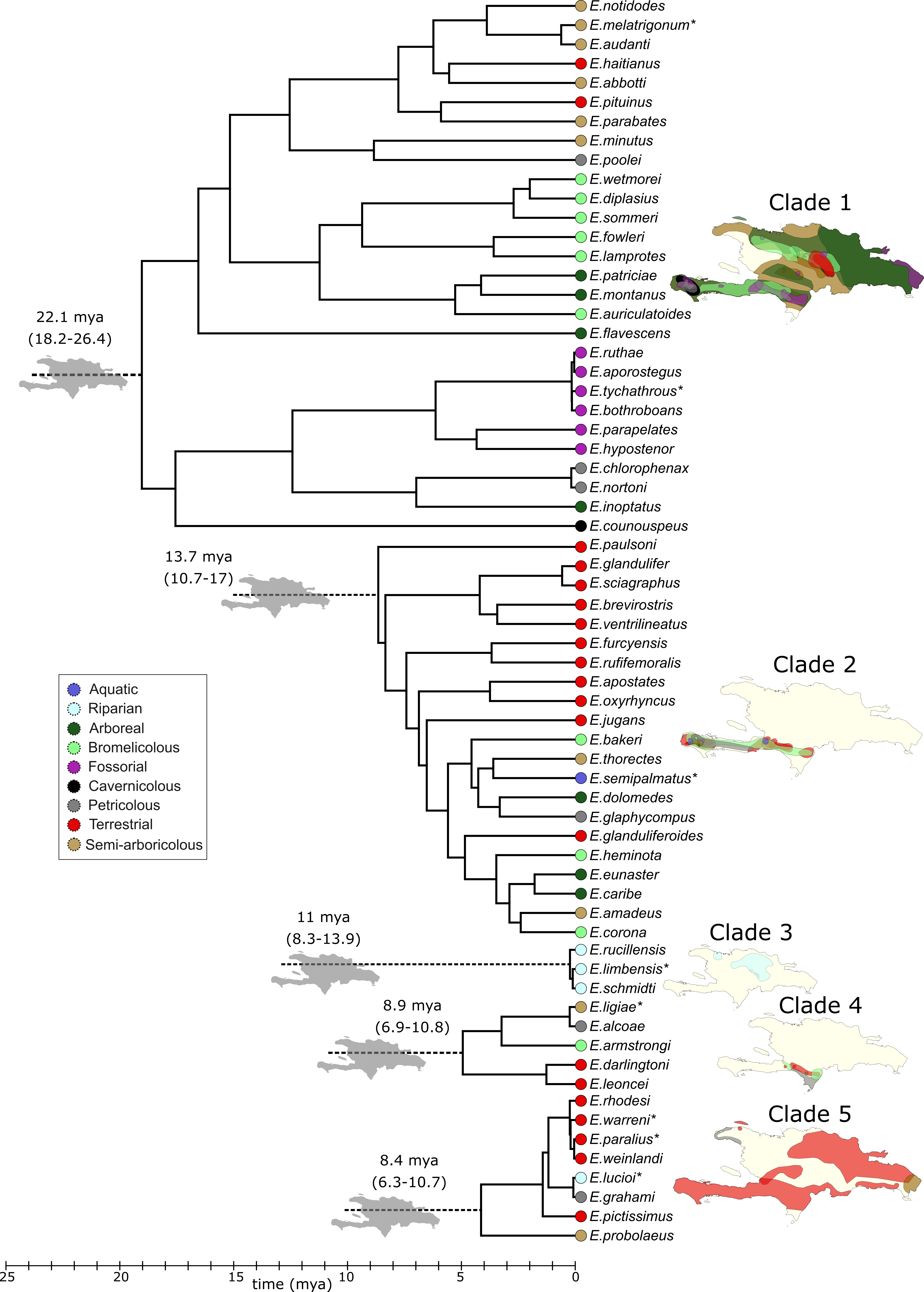

### Goodness_of_fit.pdf

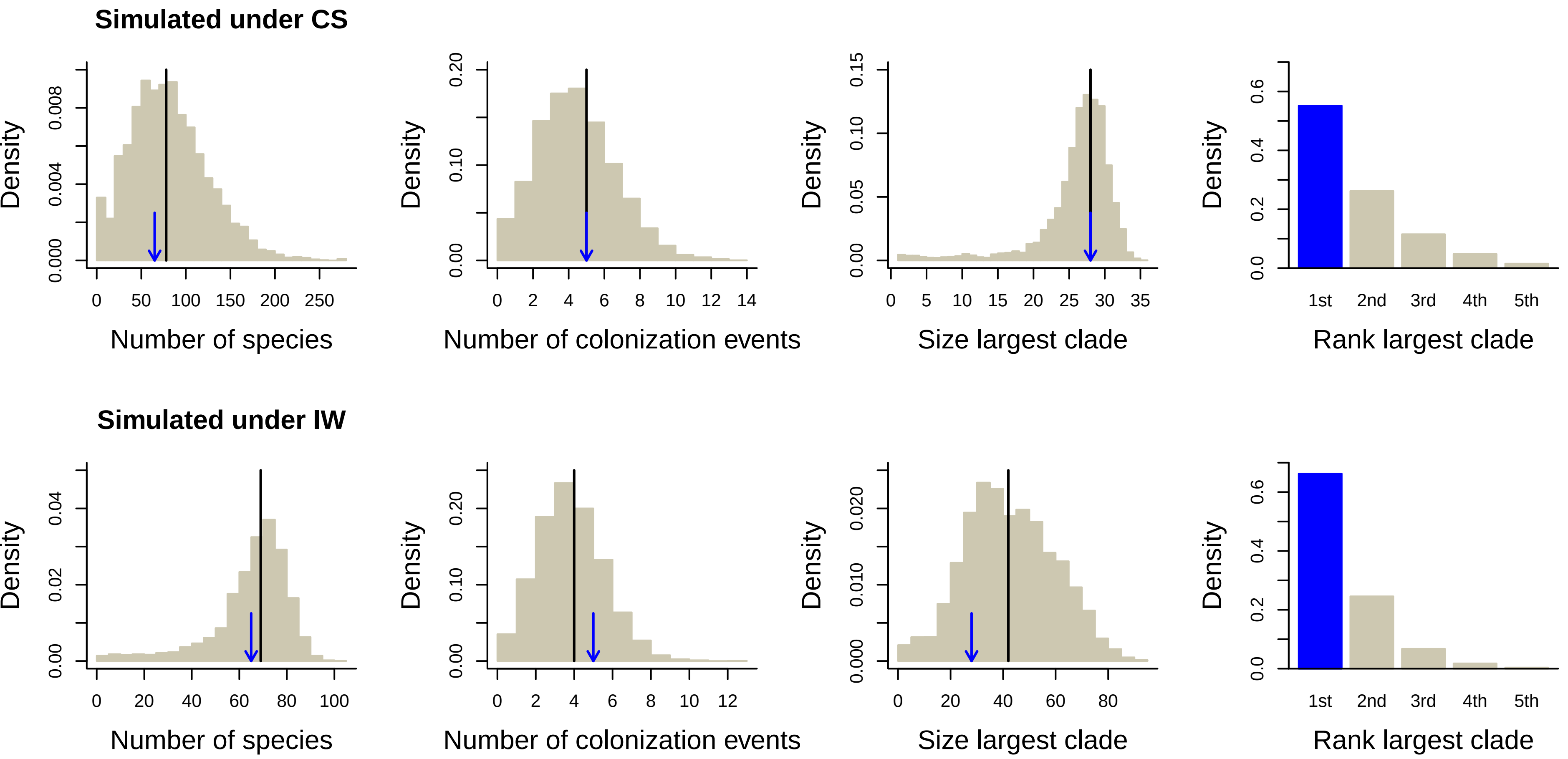

### likelihood_example_fig1.png

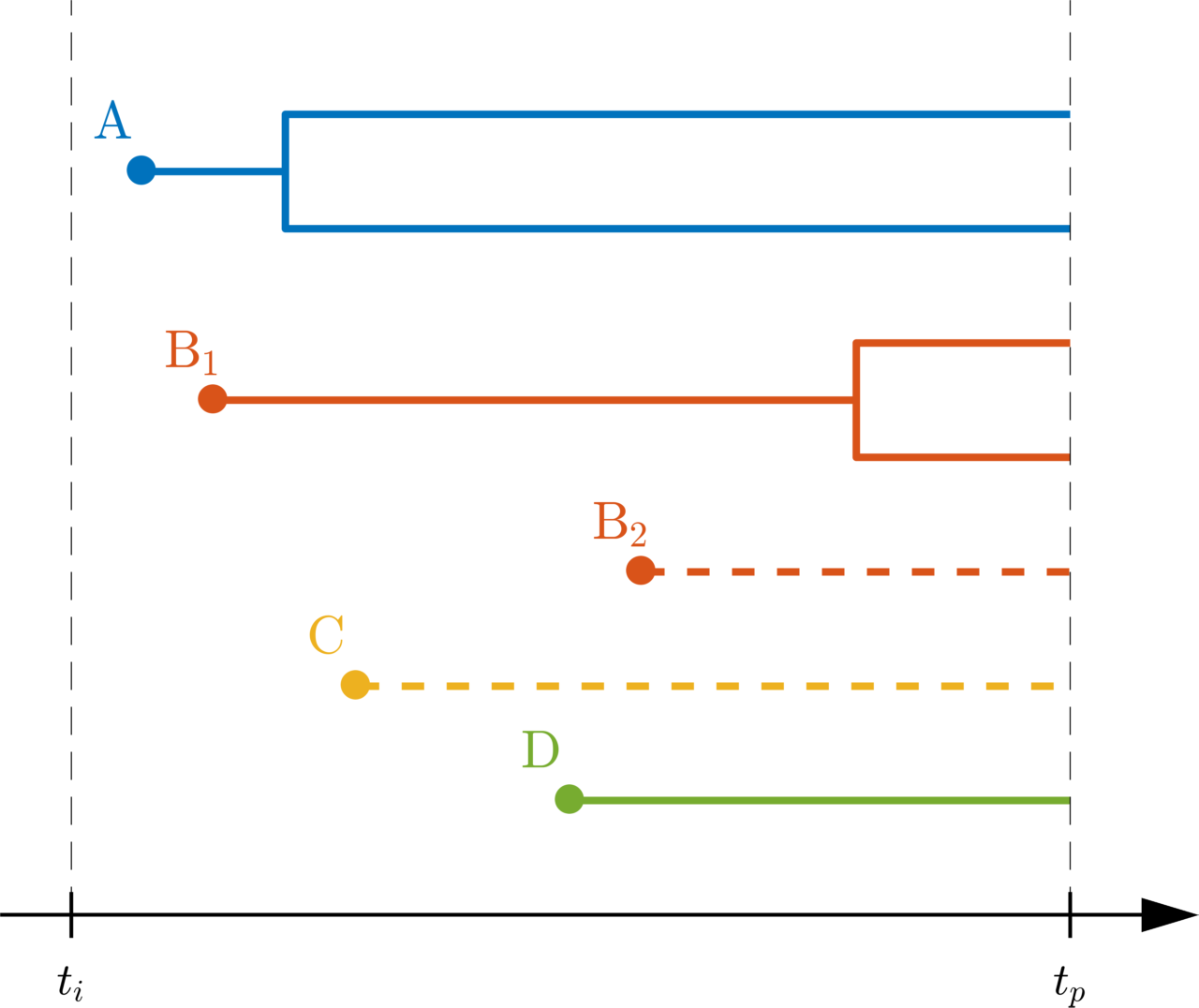

### likelihood_example_fig2.png

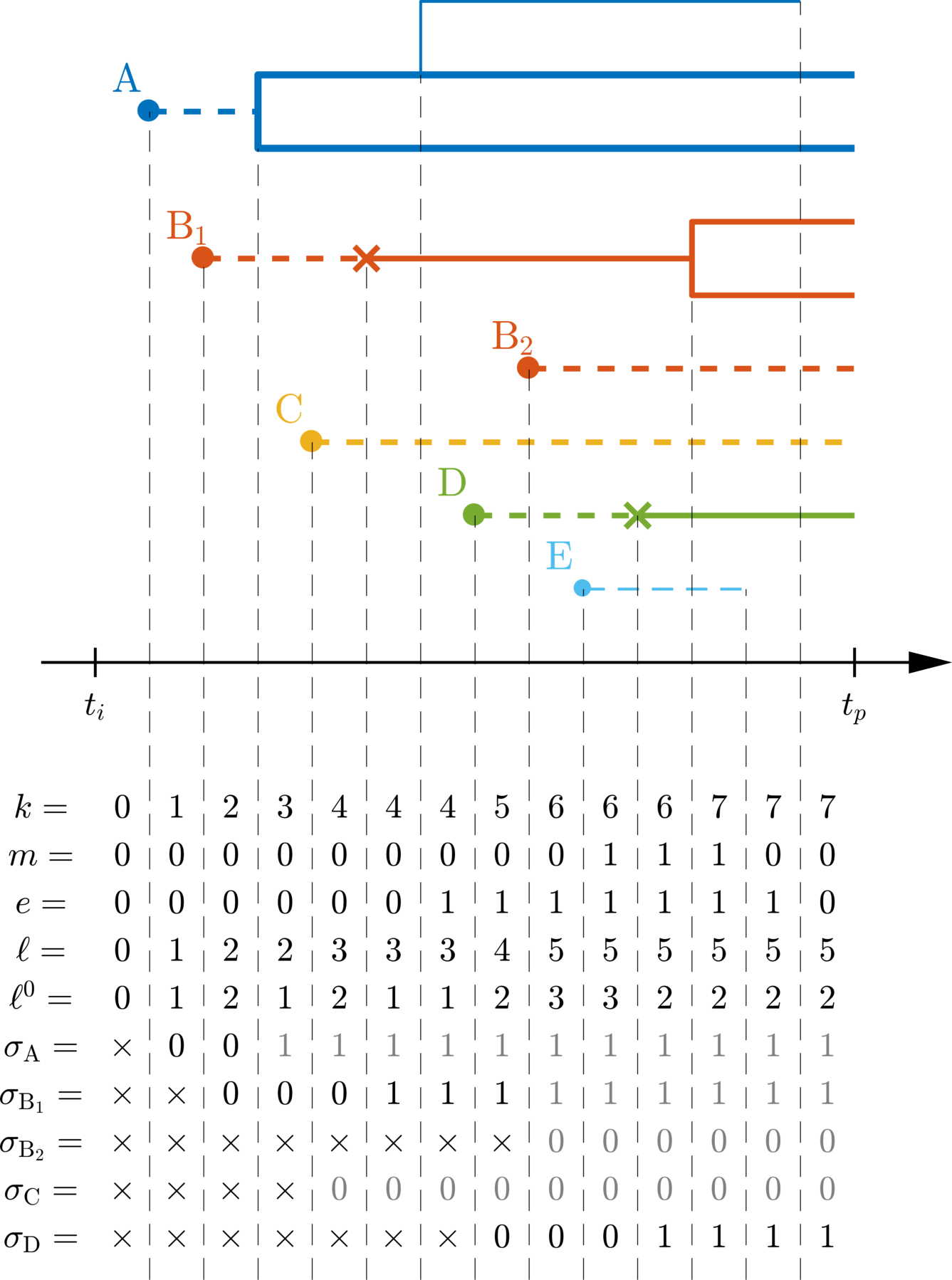

### LL.pdf

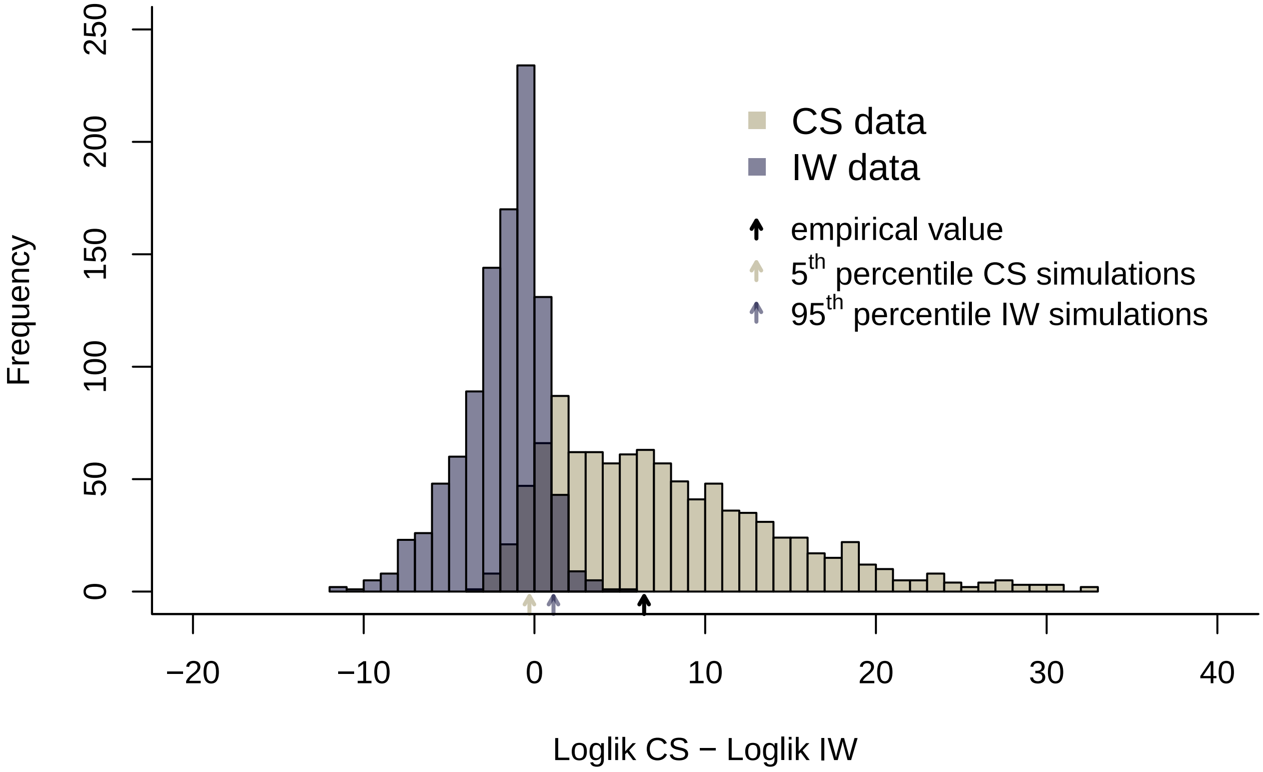

### LTT.pdf

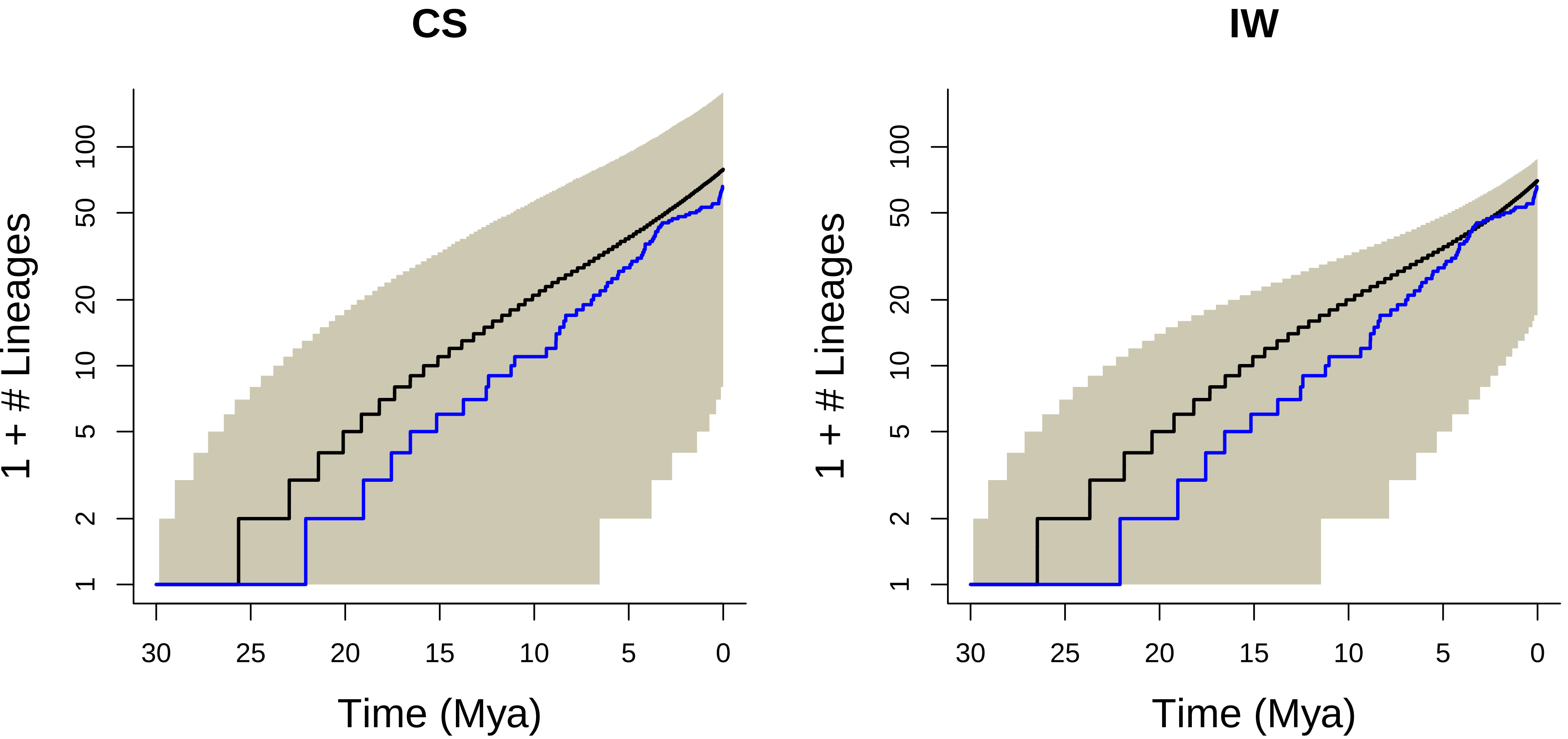

### Saturation.pdf

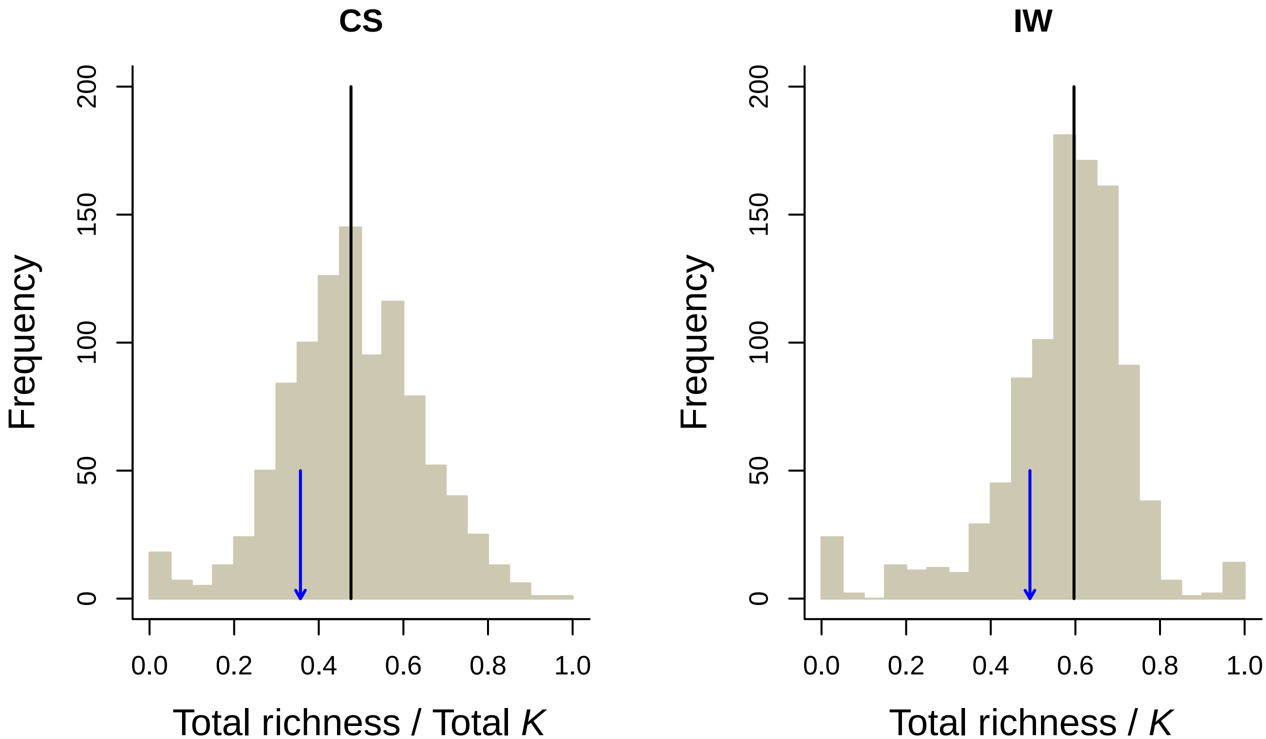
